# Deciding when to decide: how recency, urgency, risk, and bias shape human sequential decision-making - a case study across the obsessive–compulsive spectrum

**DOI:** 10.64898/2026.08.21.746184

**Authors:** Ahmed H. Abdelrazik, Peter Dayan

## Abstract

Deciding when to stop gathering information and commit to a choice is a fundamental challenge in decision-making under uncertainty. Normative characterizations such as Partially Observable Markov Decision Processes (POMDPs) prescribe mathematically optimal stopping rules; however, human evidence gathering systematically departs from optimality. Pathological departures – such as the excessive indecisiveness characteristic of obsessive-compulsive disorder (OCD) – offer an important opportunity to investigate the cognitive mechanisms involved in stopping. We extend a POMDP framework to incorporate key candidate suboptimalities: a biased prior belief, transient evidence exaggeration, progressive forgetting, boosted costs of error, temporal regulation (patience and urgency), and misperception of a deadline. We evaluate this model in a pre-existing dataset comprising 105 participants spanning healthy controls, generalised anxiety disorder, and the OCD spectrum performing an information gathering task with controlled, stochastic, deadlines. Model comparison reveals that human sequential choices are broadly governed by subjective risk penalties and time-dependent urgency, with a smaller and less certain contribution from an over-weighting of recent evidence, which a random-effects comparison does not support at the population level. Individuals differ in how that overweighting is implemented: in one deadline condition, subjects divide almost evenly between models carrying a transient exaggeration of the newest sample, models carrying progressive forgetting of older evidence, and models carrying no recency mechanism at all. Crucially, while risk sensitivity and choice stochasticity act as shared mechanisms across conditions, mechanisms such as belief bias and patience are more variable. Finally, using OCD as a clinical case study, we demonstrate that simulating choices from the fitted exaggeration model reproduces model-agnostic regression signatures of clinical indecision, which the forgetting and no-recency accounts do not. These findings offer a generative foundation for dissecting clinical departures in information gathering across the obsessive-compulsive spectrum.

**Author summary:** When people gather information before deciding, how do they choose when to stop, how does this depart from the theoretically optimal strategy, and how are the answers to these questions different in disorders involving indecision such as obsessive-compulsive disorder (OCD)? We reanalyze data from a recent study in which a large number of participants, including some on an OCD spectrum, exhibited a specific suboptimality. We examine the relative importance of a small number of interpretable departures from optimality, including a biased prior, over-weighting of most recent evidence, exponential forgetting of old evidence, excess costs for an incorrect choice, and a patience and urgency signal.

## Introduction

Decision-making in the face of uncertainty involves a number of steps. Along with gathering and assimilating information relevant to choices are the critical questions: when should we stop gathering evidence and commit to which of the options? In other words, deciding when to decide is itself a form of decision-making [1, 2]. The answers to these questions determine the balance between the speed of making the choice and the accuracy that results. A malfunction in any of these processes can produce disruptive psychiatric symptoms, from pathological indecision to premature, poorly evidenced commitments [3–5].

In decision-theoretic terms [6], this class of problems can typically be framed as Partially Observable Markov Decision Processes (POMDPs) [7]. Venerable decision-making methods involving the sequential accumulation of evidence toward a decision boundary, from Wald’s sequential probability ratio test [2] to the drift-diffusion model (DDM) [8–12], are optimal or sometimes approximately optimal policies for the underlying POMDPs [7, 13], providing answers to the questions above. Factors such as urgency – a progressive unwillingness to continue sampling [14] – turn out to be appropriate under various circumstances such as stochastic deadlines [15] or when the quality of the available evidence varies in an unsignalled manner [16]

However, suboptimalities have been documented across many settings, and are cast into stark light by the POMDP framing. One form of suboptimality is excess urgency – perhaps consistent with the proposal that urgency is a dissociable component of the decision process, separate from the accumulation of evidence itself [17]. A second suboptimality concerns the accumulation of evidence: rather than being integrated linearly, there is often a recency bias, whereby the most recently sampled information is over-weighted relative to what came before [17–19], and sometimes a primacy bias favouring early samples [20].

These mechanisms, together with more basic departures from optimality such as a biased conversion of evidence into belief and a subjective degree of risk sensitivity associated with being wrong, are natural candidates for explaining the suboptimal information gathering that is widely, though differently, observed in both health and diseases in which indecisiveness [3] or prematurity [4, 5] are evident. The differences exhibited between these populations make it important to assess the relative contributions of the various mechanisms.

Obsessive–compulsive disorder (OCD) offers a well-characterised setting in which to study exactly these questions [21]. OCD is characterised by recurrent, intrusive, and distressing thoughts (obsessions) and repetitive behaviours or mental acts (compulsions) performed to alleviate anxiety or neutralise obsessions; intolerance of uncertainty is higher in OCD patients with checking compulsions than in those without [22], and has been linked to collecting more information before committing, contributing to indecisiveness [23, 24]. Both the mechanisms above have separately been proposed to explain this indecisiveness. In terms of urgency, the signal that ordinarily grows with elapsed time appears to be attenuated in OCD, with patients continuing to sample past the point at which healthy control participants generally stop [3, 25]. Continuing to sample can look like the more accurate choice in isolation, since more evidence generally improves accuracy. It only becomes a genuine problem, when there is a deadline, a cost to continued sampling, or both.

In terms of recency, a magnetoencephalography (MEG) study by del Río and colleagues found that the normal over-weighting of the most recent sample is instead attenuated along the obsessive– compulsive spectrum, with individuals of more severe compulsivity relying less on the update in evidence strength carried by the most recent sample (Δ*ES*) [26]. Convergent evidence comes from other paradigms: a prospect-valence learning model of Iowa Gambling Task choices reported a lower recency parameter in OCD patients [27], and OCD patients have been found to accumulate more evidence while being less sensitive to it than controls [28]. The picture is not uniform, however, as one study found that OCD patients required significantly fewer draws than controls before committing in the famous ‘beads-in-the-jar’ information gathering task [29], whereas another found the opposite, in clear contrast to the delusional patients tested in the same study, who instead showed the classic “jumping to conclusions” bias of committing after too little evidence [30, 31]. Task design, sample composition, and different OC spectrum therefore shape what is observed, underscoring the need for a common, mechanistic account rather than just task-specific descriptive comparisons.

Here, we take advantage of the extensive body of data collected by del Río and colleagues [26] spanning OCD, generalised anxiety disorder (GAD), matched controls, and non-clinical high- and low-compulsivity groups. Their study began with a large crowd-sourced sample tested on a smartphone version of an information gathering task; this guided their subsequent lab-based MEG study that we analyse throughout. They performed a GLM/GLMM analysis to examine the factors encouraging participants to stop collecting evidence, showing that the weight on the most recent piece of evidence tracks obsessive-compulsive symptom severity dimensionally [26]. This association was specific: only the weighting of the most recent evidence update was related to symptoms, both behaviourally and neurally, whereas the overall tendency to decide, the number of draws taken, the horizon condition and the weighting of previously accumulated evidence showed no relation [26].

A model-agnostic regression of this kind, however, cannot by itself arbitrate between different mechanistic hypotheses – urgency attenuation, recency mis-weighting, or some combination of the two and other candidate mechanisms – nor verify that any of them arises from a coherent generative process of choice, nor determine whether a single mechanism set describes the whole population or whether different individuals rely on different combinations. We therefore modelled the task as a POMDP extended with the non-normative mechanisms described above, and fitted it to each of the 105 participants individually. We asked four questions: First, which mechanisms are actually necessary? We compared 78 candidate models spanning combinations of belief bias, recency, subjective risk and temporal regulation. Recency itself admits two distinct implementations: a transient exaggeration of the most recent sample, or an exponential forgetting of older evidence. Second, whether the two mechanisms most often proposed, recency and urgency, contribute comparably? We tested this by isolating each in turn using one further ablation model fitted for that purpose. Third, whether the structure of model describes everyone – with different parameters fitted for each subject, or whether individuals differ in which mechanisms they require? Fourth, whether the fitted mechanisms relate to obsessive-compulsive symptoms, and whether any such relation follows the continuous dimension or the diagnostic groups? Moreover, we benchmark the mechanistic account against the descriptive GLM/GLMM on identical data, check that each retained mechanism can be recovered from simulated behaviour, and quantify how well its parameters are identified. The result is a template for moving from a model-agnostic description of suboptimal information gathering to a mechanistic one.

## Behavioural task and data

### Task description

The task and dataset were adapted from [26], who developed an information-gathering paradigm to investigate sequential decision-making in obsessive–compulsive disorder (OCD). Participants completed the task while neural activity was recorded using magnetoencephalography (MEG); however, the present study analysed behavioural responses only.

Participants were required to judge which of two stimulus categories (yellow or blue cards) was more abundant in a sequence of observations. A display containing five cards (in random positions) was presented every 1,250 ms (Fig. 1). Each sample contained a varying combination of yellow and blue cards (generated as independent draws from an underlying probability that participants did not know), providing noisy evidence about the underlying majority category. Participants could press a button at any point to report whether yellow or blue cards were more abundant, or continue waiting for additional evidence up to a horizon, after which they were deemed to have failed to respond.

**Fig 1.**
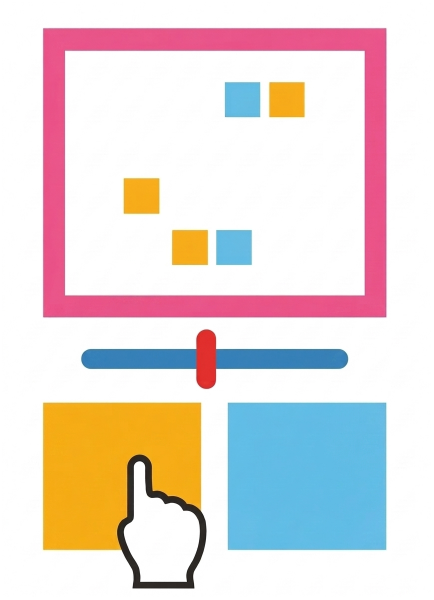
Example evidence sample: 5 yellow/blue cards per draw, in random positions.

The horizon was determined stochastically. In one, short-horizon condition, sequences lasted between 4 and 8 draws, whereas in the other, long-horizon condition, sequences lasted between 10 and 14 draws. The horizon condition was indicated by frame colour (green or pink, counterbalanced across participants who were instructed as to its meaning), and each participant completed 80 intermixed games per condition. Importantly, evidence presentation continued until the predetermined sequence endpoint, regardless of whether a participant had made an earlier decision. Correct responses were determined by the underlying generative probability of the stimulus category. Details of how the generative probabilities were defined and how sequences were generated are provided in S1 Appendix.

Participants first completed a 12-game practice block to familiarise themselves with the task and horizon manipulation; performance on this practice block was not remunerated. Subsequently, performance was incentivised through a points-based system: correct decisions yielded +2 points, incorrect decisions yielded −2 points, and failure to respond before the horizon resulted in a −1 point penalty. A visual progress bar provided trial-based performance feedback, beginning at 50% and increasing or decreasing by 10% following correct or incorrect responses, respectively. Participants received an additional monetary reward (£0.50) if the progress bar reached 100%.

### Data

Del Rio initially recruited 115 participants and excluded ten of them prior to analysis: two for not meeting the clinical criteria for GAD, one for comorbid OCD and GAD, one for completely missing questionnaire data, one for an excessive number of non-decision trials (approximately 55%), resulting in below-chance performance, and five due to technical difficulties and/or poor MEG data quality, yielding a final sample of 105 participants [26].

The final sample comprised: OCD patients (*n* = 29), GAD patients (*n* = 17), matched healthy controls (*n* = 19), and non-clinical participants with high (*n* = 20) or low (*n* = 20) compulsivity (Table 1). OCD and GAD patients were recruited through NHS services, charities, and advertisements, and controls were drawn from areas of comparable socioeconomic status to ensure similar backgrounds; the non-clinical high- and low-compulsivity groups were recruited from a population-based sample of young people (the U-CHANGE study) and stratified on the revised Padua Inventory (PI-WSUR). All participants underwent a structured clinical interview (SCID) administered by experienced researchers, and each patient met the diagnostic criteria for only one of the two disorders. Symptom severity in patients was assessed using Y-BOCS interviews, and OCI-R scores were recorded for all participants. Exclusion criteria, applied on the basis of the clinical interview, included current use of antipsychotic medication, severe learning disability, psychosis, bipolar disorder, autism spectrum disorder, substance abuse disorder, tics/Tourette disorder, or personality disorders other than obsessive-compulsive personality disorder [26].

**Table 1.** Demographic and clinical characteristics by group.

|  | Low Comp. | High Comp. | Control | OCD | GAD |
| --- | --- | --- | --- | --- | --- |
| *N* | 20 | 20 | 19 | 29 | 17 |
| Age | 21.4 ± 2.5 (18–26) | 20.8 ± 2.3 (18–26) | 29.6 ± 8.4 (18–44) | 29.1 ± 8.5 (18–45) | 31.8 ± 8.7 (20–48) |
| Sex (M/F) | 7/13 | 6/14 | 5/14 | 7/22 | 5/12 |
| IQ | 113.7 ± 9.7 | 113.5 ± 8.7 | 111.4 ± 10.3 | 106.4 ± 14.3 | 112.0 ± 8.9 |

Participants completed a total of 8,400 trials in each horizon condition. Accuracy was defined as the proportion of correct responses out of all trials, including those (‘missing’) in which no response was recorded:

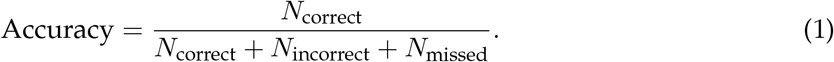

Overall task accuracy was 70.2% in the short-horizon condition and 77.1% in the long-horizon condition (Table 2). The missing rate differed between conditions (9.1% short vs. 2.0% long). The numbers of incorrect responses across conditions were 1, 743 (short) vs. 1, 756 (long). The higher accuracy and lower missing rate in the long-horizon condition are consistent with participants being able to gather more evidence before committing to a decision, in line with the POMDP framework’s prediction that extended observation opportunities improve belief precision. Fig 2 shows the aggregate choice behaviour across all subjects, giving for each evidence state the fraction of trials in which participants waited, chose yellow, or chose blue.

**Table 2.** Decision statistics broken down by horizon condition. Accuracy = Correct/Total; Missing Rate = Missed/Total.

| Condition | Total | Correct | Incorrect | Missed | Accuracy | Missing Rate |
| --- | --- | --- | --- | --- | --- | --- |
| **Short** | 8400 | 5894 | 1743 | 763 | 0.702 | 0.091 |
| **Long** | 8400 | 6473 | 1756 | 171 | 0.771 | 0.020 |

**Fig 2.**
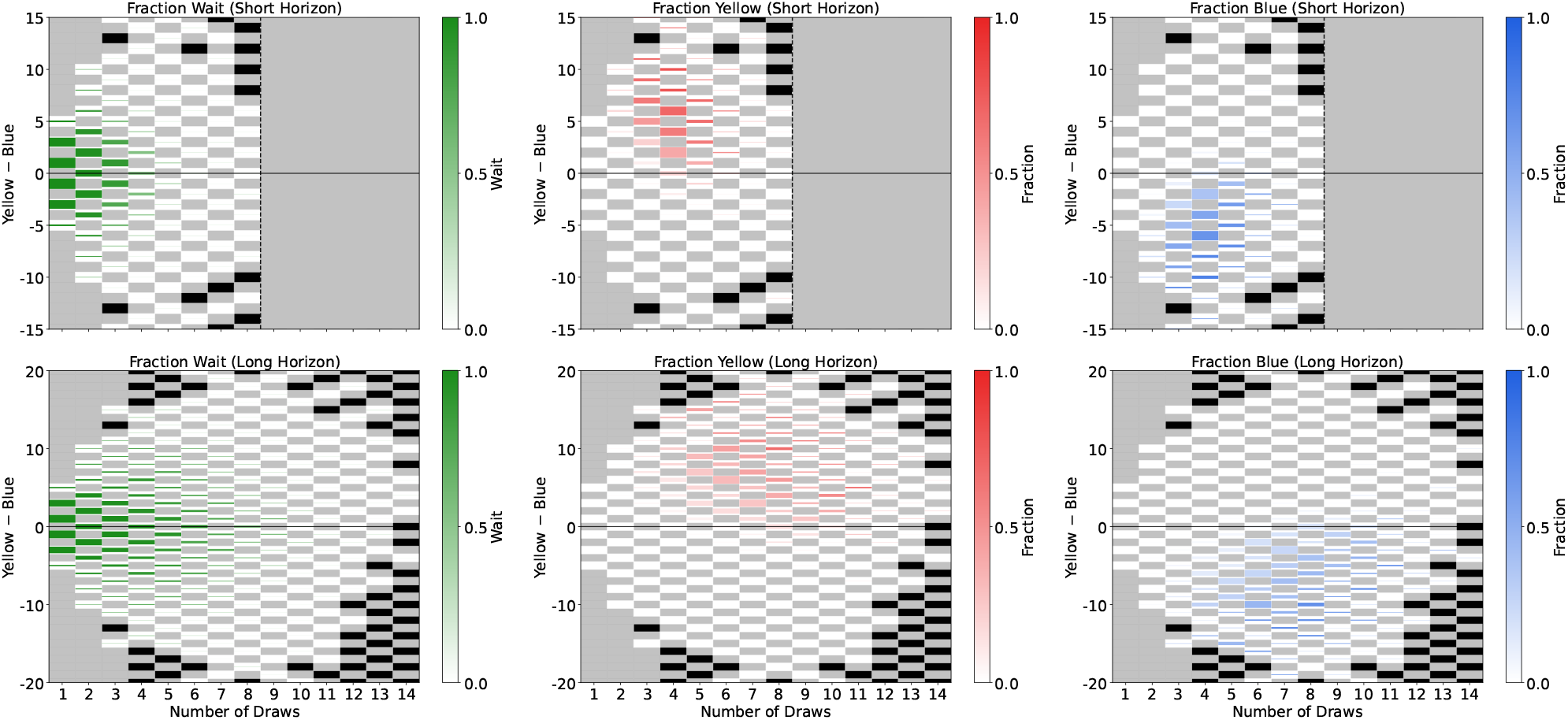
Empirical choice statistics aggregated across all subjects for the short (top) and long (bottom) horizon conditions. Each panel shows, for every visited state, the fraction of trials in which subjects chose to wait (left), chose yellow (centre; drawn in red, since a yellow ramp is hard to see against the pale cell backgrounds), or chose blue (right). Colour intensity encodes the choice fraction, so a vivid colour indicates that the corresponding behaviour dominated at that state. The cell background distinguishes three situations. Grey marks a state the task cannot produce, either because |*n*_*y*_ − *n*_*b*_| exceeds the 5*d* cards seen after *d* draws, or because it has the wrong parity, since *n*_*y*_ + *n*_*b*_ = 5*d* forces the difference to share the parity of 5*d* and only every other value is therefore attainable at a given draw; in the top row the same grey marks draws beyond the eighth, where the short-horizon condition has already ended. Black marks a state the task could produce but that no subject ever visited, so the black cells trace the frontier of the region participants actually explored. White marks a visited state at which that particular action was never chosen, so an action with zero occurrences stays visible as an empty cell rather than disappearing into the background. The area of colour in each cell is proportional to the raw count of that specific action at that state, normalised independently within each panel by the maximum raw count across all states in that panel, so a greater area reflects more absolute occurrences of that action. This dual encoding distinguishes states where a high fraction arises from many observations (large area, vivid cell) from states where it arises from only a handful of visits (small area, vivid cell). The *x*-axis represents the number of draws and the *y*-axis the difference between the cumulative difference between the number of yellow and blue cards observed (*n*_*y*_ − *n*_*b*_), where positive values indicate more yellow evidence and negative values more blue evidence. The vertical range is cropped to the evidence differences that actually occur; all short-horizon visits and 99.4% of long-horizon visits fall within the plotted range. The dashed vertical line in the top row marks the end of the short-horizon condition, and the horizontal line marks *n*_*y*_ − *n*_*b*_ = 0, where the evidence is balanced.

### Model

We modelled the decision-making task as a POMDP [7, 13]. This framework captures how individuals gather and process information when faced with incomplete or noisy evidence. We first calculated the objectively optimal policy, and then, based on observations from [26], considered suboptimalities associated with undue urgency to decide, forgetting past evidence and excess attention to immediate evidence.

### POMDP formulation

The belief states of the POMDP are defined by the sufficient statistics; here the number of yellow and blue cards, i.e., *s* = (*n*_*y*_, *n*_*b*_). At each draw there are *n* = 5 cards, so the number of draws can be calculated as *t* = (*n*_*y*_ + *n*_*b*_)*/n*. Three actions are available: choose yellow *Y*, choose blue *B*, or wait (i.e., continue sampling) *W*. The subjective transition probability of observing *i* ∈ {0, …, *n}* new yellow cards is assumed to come from a Beta-Binomial predictive distribution. As in the experiment, rewards are *R*_cor_ = −2 for a correct choice, *R*_inc_ = − 2 for an incorrect choice, and *R*_miss_ = −1 for missing the deadline. We also include two conventional suboptimalities: the policy follows a softmax function with temperature *τ* and lapse rate *ξ*. We fit these as two free parameters.

#### Belief calculation

To capture how participants accumulate evidence when judging which colour is more probable, we modelled them as Bayesian agents who maintain and update a belief over the generative probability *q* of the observed draws. On each trial, the decision reduces to comparing the posterior probability that the generative process favours one colour against the other. For yellow, this is the posterior probability that *q >* 0.5 given the *n*_*y*_ yellow and *n*_*b*_ blue cards revealed so far:

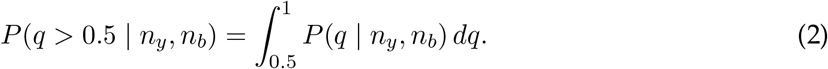

Using Bayes rule, the posterior distribution of *q* is proportional to the product of the prior belief and the likelihood of the data. We assume a Beta distribution for the prior, *P*_0_(*q*) = Beta(*q α, β*), and a Binomial distribution for the likelihood of the observed counts. A key property of this model is conjugacy, which ensures that the posterior distribution is also a Beta distribution, with updated parameters:

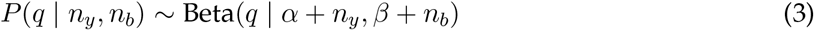

To allow for prior biases, we fix *α* = 1 and treat *β* as a free parameter to be estimated during the fitting procedure. We found that leaving both free led to problems with parameter recovery; and varying *β* – which we refer to as the belief bias – sufficed.

#### Transition probability

The subjective probability of reaching the next state *s*^*′*^ and observing *I* new yellow items depends on the current belief, characterised by the evidence (*n*_*y*_, *n*_*b*_). In our POMDP formulation, the transition dynamics specify the probability of drawing a given number of yellow cards. Since each draw consists of *n* = 5 cards, the transition probability is given by,

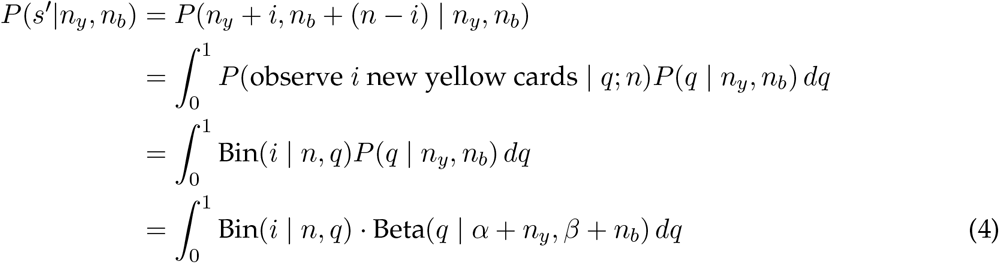

#### Generative hazard function

The generative hazard function *h*(*t*) is shown in Fig 3. Within each horizon window – draws 4 to 8 in the short condition and draws 10 to 14 in the long condition–the probability of termination is uniform, equal to 0.2 at each draw (one of five possible endpoints). The generative hazard at each time step is calculated using the standard formula:

**Fig 3.**
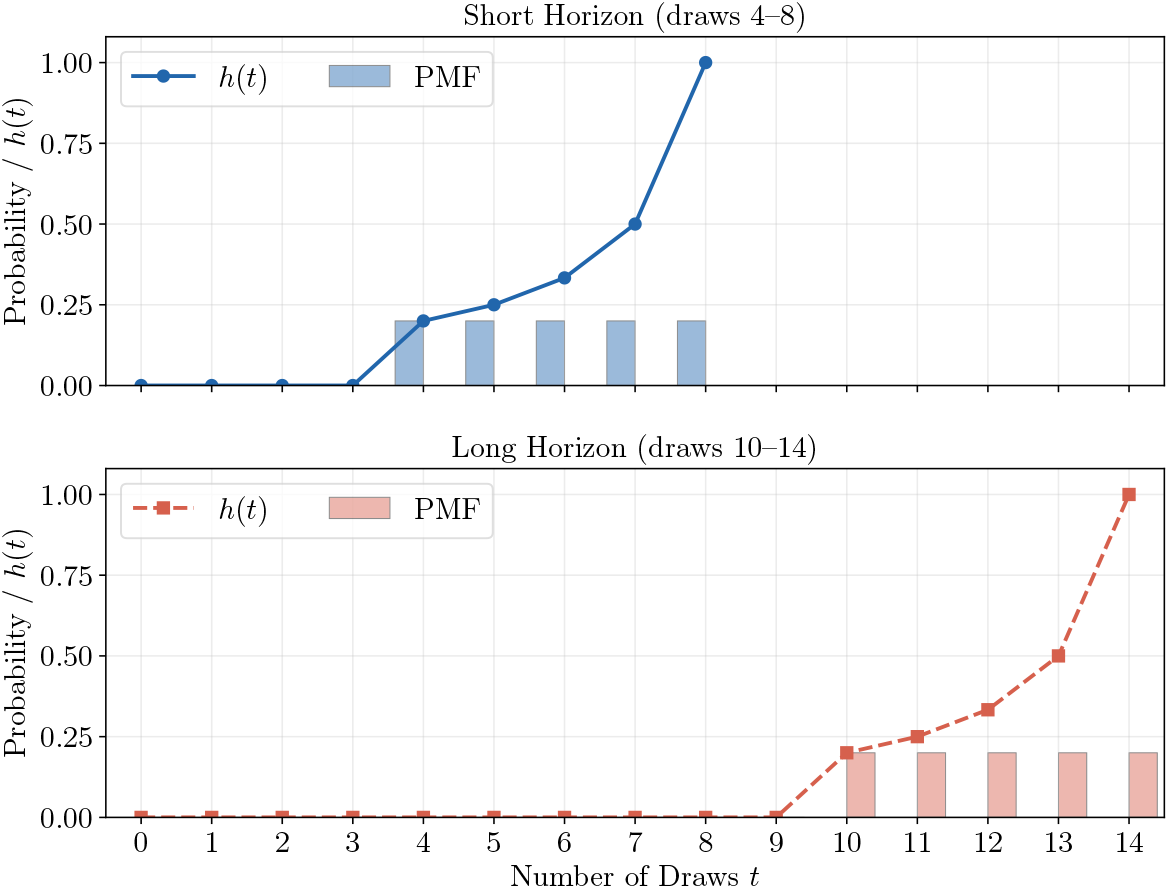
Discrete generative hazard function for the short (top, draws 4–8) and long (bottom, draws 10–14) horizon conditions. Within each horizon window the termination probability is uniform (*p* = 0.2), so the discrete hazard *h*(*t*) rises across the window as the remaining probability mass shrinks.

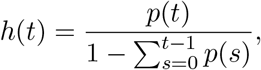

where *p*(*s*) denotes the uniform probability mass function (PMF).

#### Action values

The action value for choosing yellow, *Q*(*Y*|*n*_*y*_, *n*_*b*_), given the current state (*n*_*y*_, *n*_*b*_), is the expected reward for that decision. It is computed by weighting the potential outcomes by their posterior probabilities:

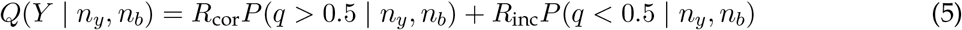

where *R*_cor_ and *R*_inc_ are the objective rewards for correct and incorrect decisions, respectively. The first probability term, *P* (*q >* 0.5 | *n*_*y*_, *n*_*b*_), is the agent’s belief that yellow is the more probable colour, as calculated in Eq 2. The second term, *P* (*q <* 0.5 | *n*_*y*_, *n*_*b*_), is the belief that blue is more probable. The action value for choosing blue, *Q*(*B*), is calculated symmetrically.

In the calculations of the action values for waiting, we have two cases: 1) to exclude the generative hazard from the calculation of the action value for waiting, and 2) to include the generative hazard in the action value for waiting. Both of them are calculated using backward induction.

#### Action value of waiting

The action value of waiting depends on the hazard the agent assumes to govern termination according to:

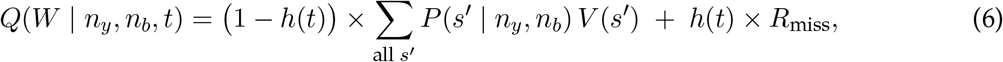

where the first term is the expected value of continuing, the second is the cost of the game ending before a decision is made, and

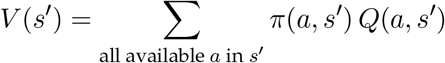

is the value of the future state *s*^*′*^, with *P* (*s*^*′*^ | *n*_*y*_, *n*_*b*_) the state transition probabilities and *R*_miss_ the cost of missing.

#### Policy function

The policy is modelled using a softmax function with a temperature parameter (*τ*) and a lapse rate (*ξ*). The lapse rate is applied *only* to the yellow (Y) and blue (B) actions, while the waiting action is left unaffected.

Let the softmax probabilities (before adding the lapse) be:

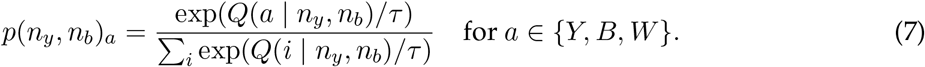

Then, the softmax policy, *π*(*a*), with lapse rate is defined as:

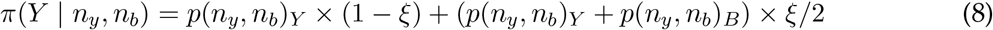

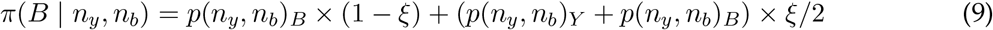

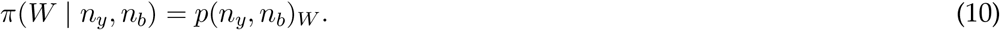

This construction ensures that, while the lapse *ξ* only redistributes probability mass between Yellow and Blue, the total probability remains preserved at 1.

Fig 4 shows the best actions for the normative model under the generative hazard function *h*(*t*), with temperature *τ* = 10^−8^ and lapse rate *ξ* = 0 (panel A). In this configuration the agent anticipates the stochastic termination of the game and commits from the fourth draw, at which the hazard starts. Panels B and C hold the hazard at this setting and perturb the policy, so both should be interpreted by comparison with panel A. When the temperature parameter is increased to *τ* = 0.3 (panel B), the policy becomes more stochastic, which effectively lowers the decision threshold and leads the agent to commit earlier (as in the analysis of jumping to conclusions in [31]). A similar effect is observed when increasing the lapse rate to *ξ* = 0.05 (panel C): the additional randomness also reduces the effective threshold, resulting in earlier commitment. In contrast, when the deterministic hazard function *G*(*t*) replaces *h*(*t*) (panel D), the game can end only at the deadline, and the optimal policy has a relatively high decision boundary, meaning the agent waits longer to accumulate evidence before committing to an action.

**Fig 4.**
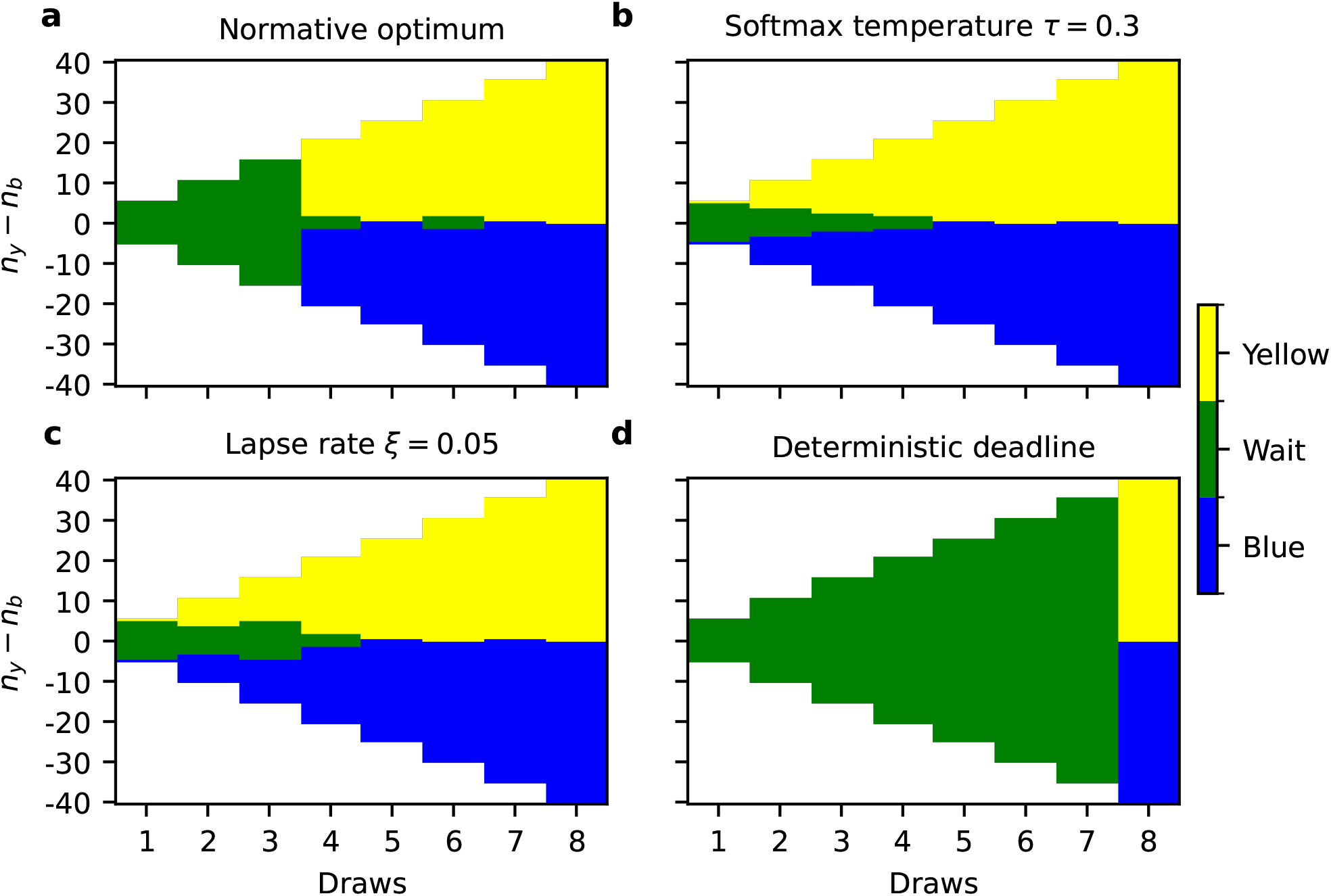
Effect of softmax temperature, lapse rate and the deadline on the best actions in the normative POMDP model. The x-axis represents the number of draws, ranging from the first to the eighth draw in the short-horizon condition. The y-axis represents the difference between the number of observed yellow and blue cards (*n*_*y*_ −*n*_*b*_), where positive values indicate more yellow evidence and negative values indicate more blue evidence. Colour gives the optimal action at each state: yellow and blue for committing to that colour, green for waiting (continue sampling). Although the heatmaps are rendered as continuous, the underlying state space is discrete, and not all (*t, n*_*y*_ − *n*_*b*_) combinations are reachable given the way the task is administered (e.g. *n*_*y*_ − *n*_*b*_ = 0 cannot occur at the odd draws). Other structurally impossible states are shown in white. a: The normative optimum for the task, under the generative hazard function *h*(*t*), which is active from the fourth draw, with a nearly deterministic policy (temperature *τ* = 10^−8^ and lapse rate *ξ* = 0). The agent anticipates the stochastic deadline and commits from the hazard onset rather than accumulating to a fixed boundary. b and c hold the hazard at this setting and perturb the policy, so both are read against a. b: Temperature is increased from *τ* = 10^−8^ to *τ* = 0.3. The policy becomes more stochastic, which effectively lowers the decision threshold and leads the agent to commit earlier. c: Lapse rate is increased from *ξ* = 0 to *ξ* = 0.05. The additional randomness reduces the effective threshold in the same direction, giving earlier and less conservative decisions. d: The same near-deterministic policy as a, but with the deterministic hazard function *G*(*t*) in place of *h*(*t*), leaving a fixed deadline. Without the prospect of a stochastic deadline, the agent is free to accumulate, and the wait region extends to a high decision boundary that it holds until the final draw.

### Heuristic mechanisms

The preceding section derives a policy directly from the task structure: Bayesian updating of the belief about the generating process, alongside dynamic programming over the payoffs, the deadline, and the true hazard function of the task. Table 3 lists a number of heuristic mechanisms that could perturb normative behaviour in the task (along with temperature and the lapse process described above).

**Table 3.** Heuristic mechanisms. “Acts on” names the part of the normative derivation the mechanism modifies.

| Mechanism | Acts on |
| --- | --- |
| Lapse rate $\xi$ | policy |
| Temperature $\tau$ | policy |
| Belief bias $\beta$ | belief |
| hazard $g(t)$ | value of waiting |
| Subjective cost $R_{\text{risk}}$ | reward/objective |
| Temporal regulation $\Phi(t)$ | value of waiting |
| Hazard lapse $L$ | task representation |
| Exaggeration $E$ | belief/transition |
| Forgetting $\gamma$ | belief/transition |

### Hazard

Although the participants faced a stochastic deadline, we considered the possibility that they might think it to be deterministically the maximum number of possible observations (8 for the short, and 14 for the long horizon conditions). This implies replacing the true hazard functions of Fig. 3 with

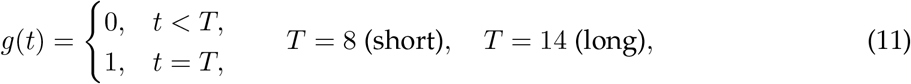

We also considered fitting the hazard function parameters, retaining its uniform termination distribution but estimating its onset and offset draws, which neither improved model fit nor recovered its own parameters.

### Risk parameter

The risk parameter *R*_risk_ captures individual differences in risk sensitivity. It is fitted per participant and acts as a subjective penalty on incorrect decisions, reflecting the degree to which an agent seeks to avoid errors. A larger |*R*_risk_| increases the perceived cost of committing to the wrong choice, thereby incentivising the agent to accumulate more evidence before deciding. The effective cost of an incorrect decision is thus defined as:

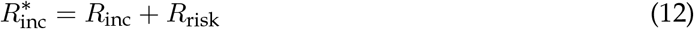

where *R*_risk_ ≤ 0, such that more negative values reflect greater risk aversion. Fig. 5 illustrates the effect of the risk parameter on the optimal policy. As *R*_risk_ becomes more negative, the effective cost of an incorrect decision 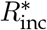 grows large enough to exceed the cost of missing the deadline, widening the decision boundary: the agent keeps accumulating evidence when the yellow-versus-blue difference is small, committing only once the evidence is sufficiently unambiguous, or otherwise waiting until the game terminates.

**Fig 5.**
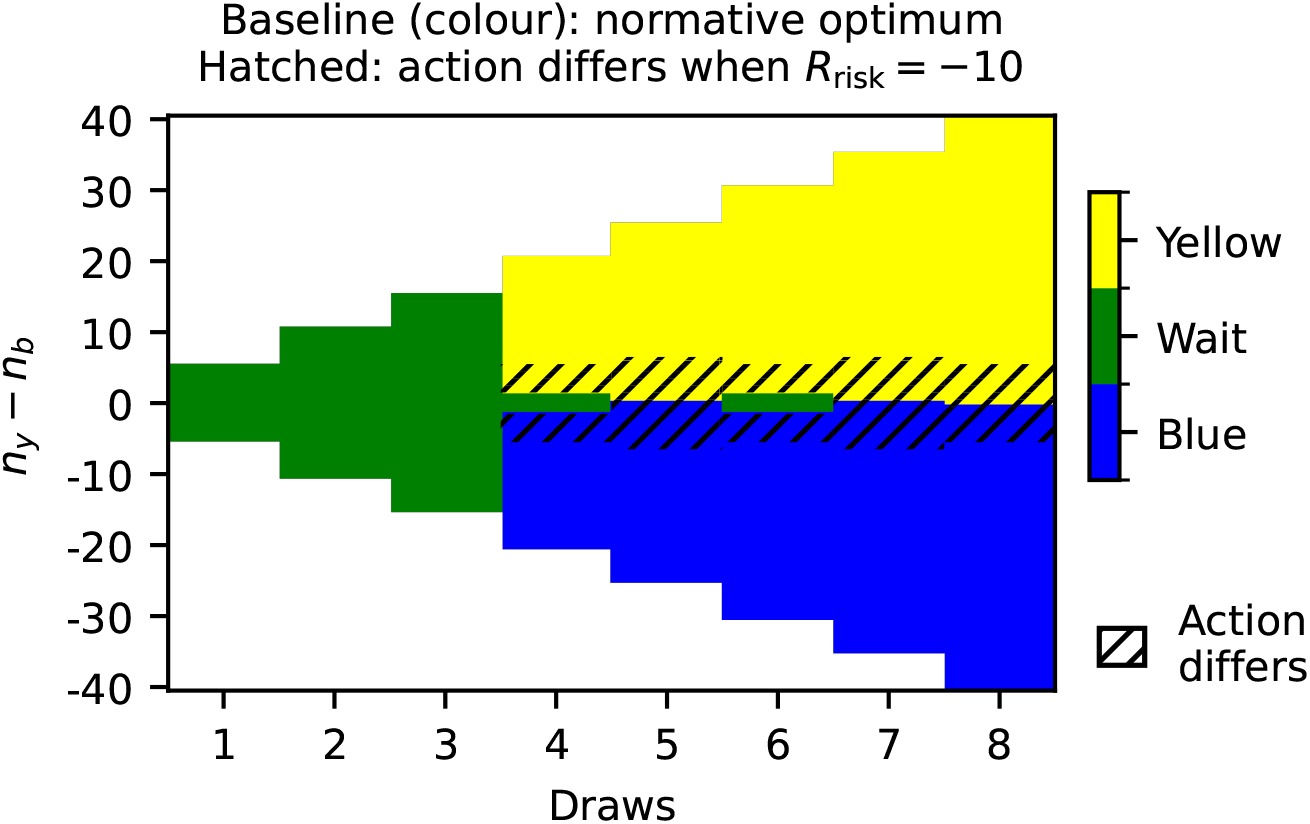
Effect of the risk parameter on the best actions in the normative POMDP model. Colour shows the normative optimum (panel a of Fig 4: *τ* ≈ 0, *ξ* = 0, *R*_risk_ = 0, with the generative hazard active); diagonal hatching marks states where the action changes once *R*_risk_ = − 10 is added, as also labelled directly on the figure. The large penalty on incorrect decisions raises the effective cost of committing to the wrong choice above the cost of missing the deadline 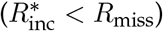, widening the decision boundary: in the hatched band the agent switches from committing to waiting, continuing to accumulate evidence until it is sufficiently unambiguous, or otherwise waiting until the game terminates.

### Temporal regulation function

The temporal regulation function is introduced to capture deviations in the subjective value of time from the POMDP model and thereby characterise aspects of urgency that are not implied, for instance, by the stochastic deadline. We implemented a nonlinear temporal regulation function in the form of a generalised sigmoid with independent maximum and minimum bounds. This allows the function to transition from rewarding patience to penalising delay. The temporal adjustment value per step Φ(*t*) evolves as a function of the number of draws *t* ∈ {1, …, *T}*, where *T* is the maximum sequence length in each condition (*T* = 8 for the short horizon and *T* = 14 for the long horizon), as follows:

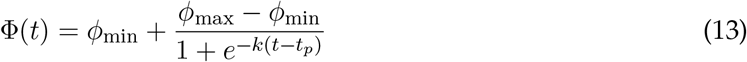

Here, Φ(*t*) partially modulates the utility of waiting by acting as either an incentive or a cost depending on the decision epoch. The parameter *ϕ*_max_ dictates the maximum positive incentive for waiting, creating a distinct patience region during early draws where Φ(*t*) *>* 0. In this region, the function acts as an explicit reward that encourages the agent to accumulate more evidence. The parameter *t*_*p*_ defines the patience time, serving as the inflection threshold where the agent transitions from patience to impatience. As *t* exceeds *t*_*p*_, the function crosses into the negative range (Φ(*t*) *<* 0), representing the onset of temporal regulation costs (or urgency penalties) that urges the agent to decide [14]. The parameter *ϕ*_min_ represents the asymptotic baseline cost—the maximum penalty per step incurred as the delay is prolonged. The parameter *k* controls the steepness of this transition–in our case it is negative since the sigmoid is inverted, as illustrated in Figure 6.

**Fig 6.**
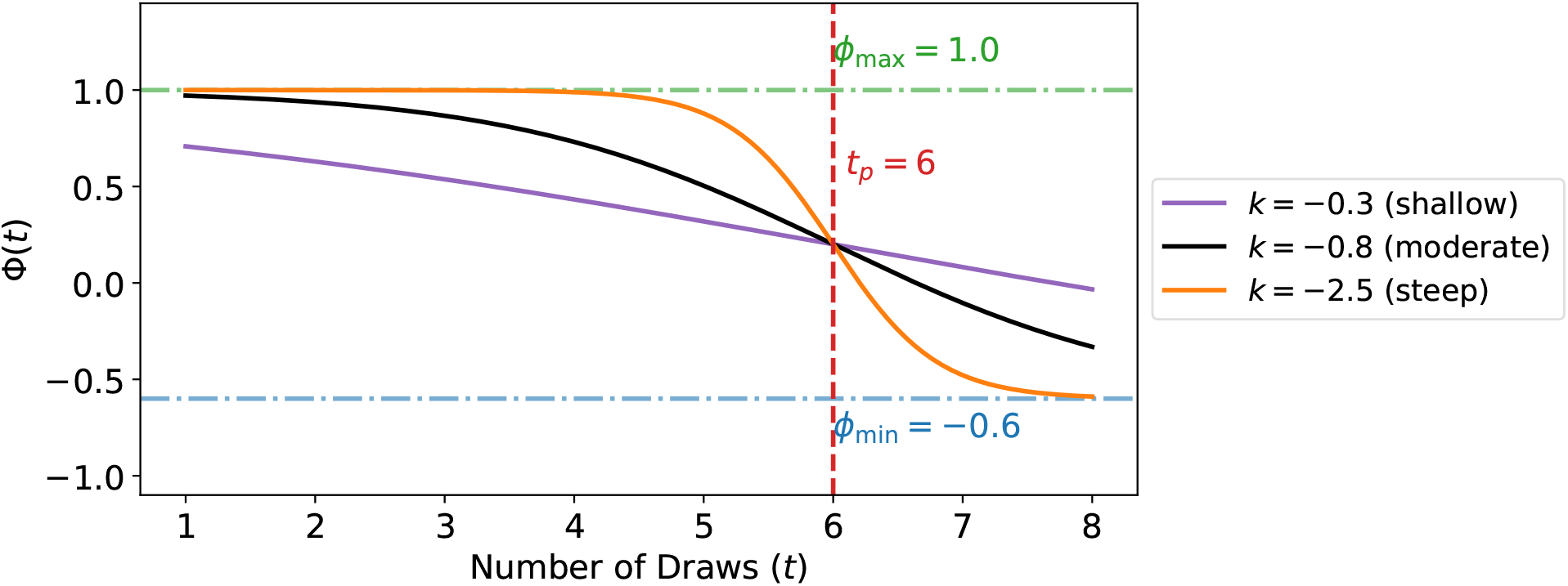
Illustration of the temporal regulation function Φ(*t*) for different slope values *k*. All curves share the same parameters *ϕ*_min_ = −0.6, *ϕ*_max_ = 1.0, and patience time *t*_*p*_ = 6, under the short-horizon condition. A shallow slope (*k* = −0.3) produces a gradual transition from the positive patience incentive to the negative temporal cost region, while a steep slope (*k* = −2.5) produces an abrupt switch. The vertical dashed line marks the patience time *t*_*p*_, before which Φ(*t*) *>* 0 (patience incentive region) and after which Φ(*t*) *<* 0 (temporal regulation costs).

The temporal regulation value is added directly to the action value for waiting in Eq. 6 as follows:

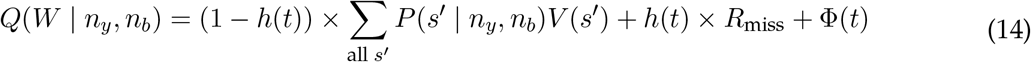

Fig. 7 illustrates the resulting effect on the optimal policy. The temporal regulation function and the hazard term *h*(*t*) *× R*_miss_ share a related contextual mechanism, though they differ in their early-draw behaviour. Both are incorporated into the action value for waiting and operate by modifying the utility of future waiting based on the decision epoch. The hazard function’s shape is illustrated in Fig. 3, while the temporal regulation function’s shape is depicted in Fig. 6. Notably, unlike the hazard term which purely penalises delay, the temporal regulation function features an initial region where waiting is actively valued as a benefit, requiring the agent to exhaust this patience utility before the action of waiting is perceived as a net cost.

**Fig 7.**
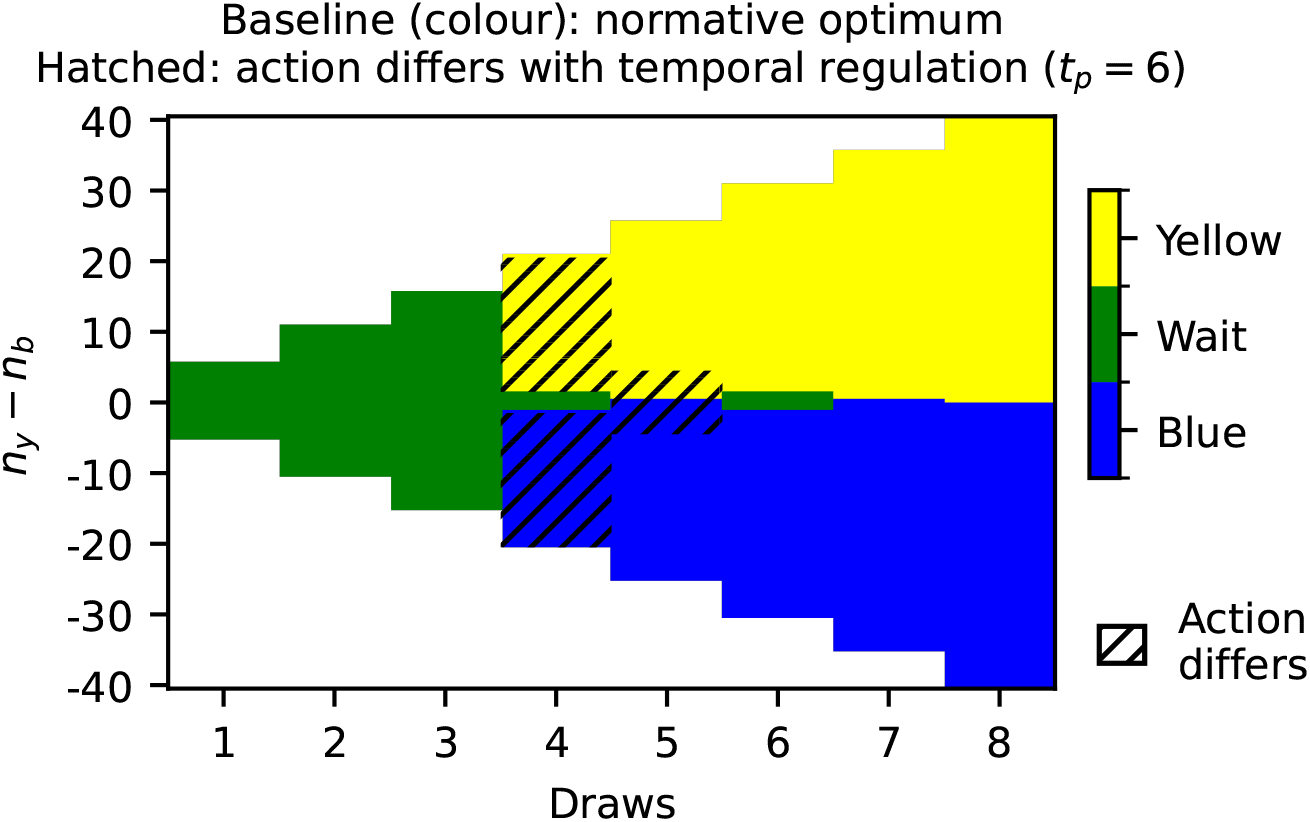
Effect of the temporal regulation function on the best actions when added to the normative model. Colour shows the normative optimum (panel a of Fig 4: *τ* ≈ 0, *ξ* = 0, no temporal regulation, with the generative hazard active); diagonal hatching marks states where the action changes once *ϕ*_min_ = − 0.6, *ϕ*_max_ = 1.0, *k* = − 0.8, and *t*_*p*_ = 6 are added, as also labelled directly on the figure. Before the patience time *t*_*p*_ = 6, the temporal regulation function is positive, providing an incentive that encourages the agent to wait and accumulate more evidence; after *t*_*p*_ = 6, it turns into a negative penalty, accelerating commitment. Accordingly, the hatched band runs along the decision boundary from the fourth draw, where the patience incentive holds the agent at waiting in states the optimum already commits in, and closes by the seventh draw once that incentive has expired.

### Hazard lapse parameter

Inspection of individual decision distributions revealed marked heterogeneity in how subjects adjusted their behaviour from short-to long-horizon trials. In the long-horizon condition, the generative hazard function begins to rise only from draw 10 onward (up to draw 14), so a well-calibrated observer should accumulate evidence until that window before committing. However, more than a third of the subjects showed decision distributions inconsistent with this: some peaked at draw counts characteristic of the *short*-horizon hazard (draws 3–5), suggesting they perceived themselves to be in the short-horizon condition despite the actual task structure; others showed bimodal distributions with peaks at both short- and long-horizon-consistent draw counts, indicating a mixture of beliefs across games; and others were consistently impulsive, responding very early irrespective of condition. Fig 8 illustrates four representative subjects with these characteristics.

**Fig 8.**
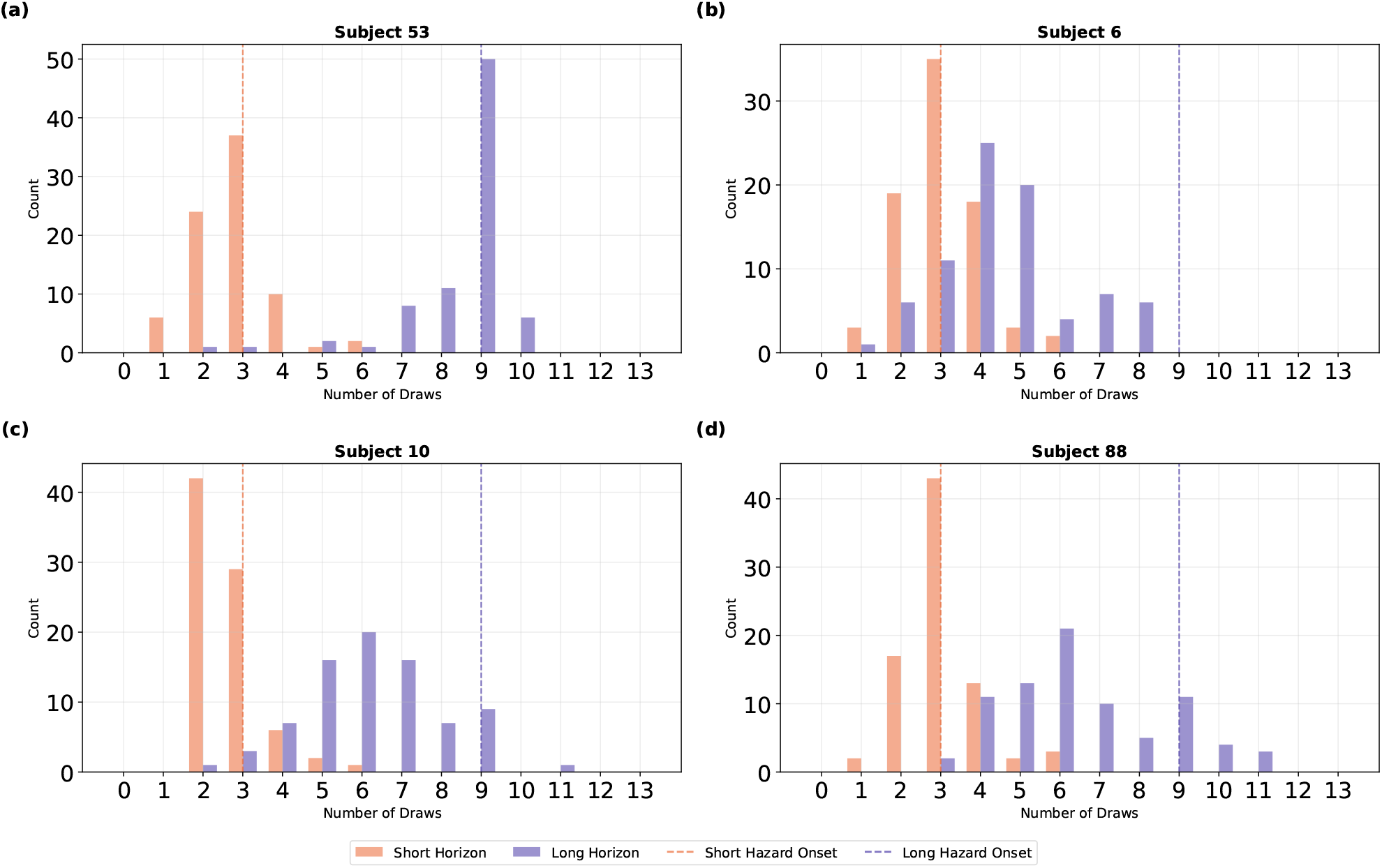
Decision distributions for four representative subjects illustrating the range of horizon-perception behaviour. The *x*-axis denotes the number of draws observed before the subject committed to a decision; the *y*-axis denotes game counts. Top panels show the short-horizon condition; bottom panels show the long-horizon condition. (**a**) Subject 53 is well calibrated, peaking just before the hazard onset in each condition (draw ≈ 3 for short, draw ≈ 9 for long). (**b**) Subject 6 apparently confuses the two conditions: although the long-horizon hazard rises only from draw 10, this subject’s decision distribution peaks at draws 4–5, consistent with short-horizon behaviour. (**c**) Subject 10 is consistently impulsive, committing at very early draw counts in both conditions. (**d**) Subject 88 shows a bimodal distribution in the long-horizon condition, with peaks at draws 6 and 9, indicating a mixture of short- and long-horizon beliefs across games.

This pattern can also be partially interpreted as a high evidence-accumulation cost: a subject who consistently decides around the 6th or 7th draw, despite the hazard not starting until draw 10, may be penalising further evidence collection rather than genuinely misperceiving the horizon. Both interpretations are therefore plausible, and they need not be mutually exclusive. To allow the model to decide between them, we fit an extended model that includes both the lapse parameter and temporal regulation function (accounting for time and accumulation costs), letting the complexity-controlled likelihood determine which account – or combination of accounts – best explains each subject’s data. This comparison is reported in the Model comparison section.

To accommodate this behavioural heterogeneity, we introduced a hazard lapse parameter *L* ∈ [0, 1] that linearly interpolates between the short- and long-horizon generative hazard functions:

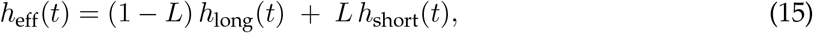

where *h*_long_ and *h*_short_ are the hazard functions for the long- and short-horizon conditions, respectively. A value of *L* = 0 indicates that the subject correctly perceives the long-horizon hazard; *L* = 1 indicates that the subject acts as though the hazard follows the short-horizon function, even when in the long-horizon condition. This parameter was estimated during long-horizon and combined-horizon model fitting, and left out of the short-horizon models. Eq 15 mixes the long-horizon hazard toward the short-horizon one, so *L* can express a single misperception: treating a long horizon as though it were short. Participants in the long-horizon condition did behave this way, committing at a median of the 7th draw with 89.7% of games ending before the hazard became active at draw 10 at all. The short-horizon condition showed no counterpart to this. Participants there committed at a median of the 4th draw, the first draw at which termination becomes possible, and nothing in the distribution of their decisions suggested they were waiting as if they are in the long horizon.

### Exaggeration

It was suggested in [26] that decision-makers may not weigh observations equally over time, but instead place disproportionate emphasis on the most recent evidence when forming beliefs and making decisions. We therefore considered mechanisms of *evidence exaggeration* that differ according to whether the amplification has a transient or sustained effect on accumulated belief.

In the *transient* variant, the current evidence is amplified only during the belief update at the present timestep; once incorporated, it returns to its original value in the accumulated evidence, so that past exaggerations do not persist. Because the agent plans over future observations, this variant admits a further distinction in how the transition probabilities are computed: the planner may either assume that it will continue to exaggerate at future timesteps, or assume that no exaggeration will occur in the future. The former treats exaggeration as a stable property of the agent that is anticipated during planning, whereas the latter treats each exaggeration as a momentary, unanticipated distortion. In the *sustained* variant, the amplified evidence remains integrated into the cumulative evidence, thereby affecting all subsequent belief updates.

We evaluated all three configurations–transient exaggeration with future exaggeration anticipated, transient exaggeration with no future exaggeration anticipated, and sustained exaggeration. In the following, we focus on the best-performing variant, in which only the current evidence is temporarily amplified during belief updating and the planner assumes no future exaggeration when computing transition probabilities.

We formalise this process through an exaggeration factor *E*, applied to the most recent draw. Although the sufficient statistics remain the total numbers of yellow and blue cards observed, for implementation purposes we distinguish between evidence accumulated before the current draw and evidence observed in the current draw. The total number of yellow observations is given by

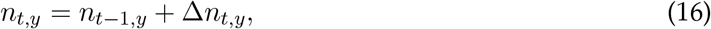

where *n*_*t*−1,*y*_ is the cumulative number of yellow cards observed before the current draw, and Δ*n*_*t,y*_ = *n*_*t,y*_ −*n*_*t*−1,*y*_ is the number of yellow cards observed in the current draw. The corresponding blue-card counts are denoted by *n*_*t*−1,*b*_ and Δ*n*_*t,b*_ = *n*_*t,b*_ − *n*_*t*−1,*b*_.

#### Belief calculation

The belief calculation follows the same Bayesian framework as the normative model (Eq. 3), with conjugate Beta-Binomial updating. The only modification is that the contribution of the current draw is scaled by the exaggeration factor *E*. The posterior distribution over *q* is thus:

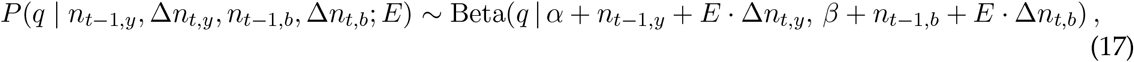

When *E* = 1, the amplified counts reduce to the standard cumulative counts *n*_*t,y*_ = *n*_*t*−1,*y*_ + Δ*n*_*t,y*_ and *n*_*t,b*_ = *n*_*t*−1,*b*_ + Δ*n*_*t,b*_, and the posterior (Eq. 17) reduces to the normative posterior (Eq. 3).

#### Transition probability

The transition probability follows the same derivation as in the normative model (Eq. 4), but with the current evidence amplified by *E*. Suppose that *i* ∈ {0, …, *n}* yellow cards are observed in the next draw, where *n* = 5 is the draw size. The transition probability is given by:

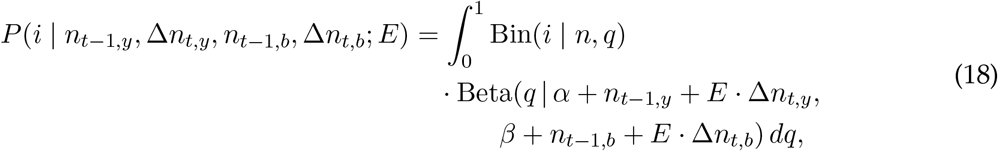

where the Beta distribution now uses the amplified sufficient statistics instead of the standard cumulative counts *n*_*t,y*_ and *n*_*t,b*_. When *E* = 1, this reduces to the normative transition probability.

### Forgetting

Another potential mechanism that can account for a recency bias in evidence accumulation is forgetting. Unlike the exaggeration model which produces a transient amplification of the current observation during belief updating before reverting to the raw count for subsequent draws, the forgetting model introduces a forgetting parameter *γ* ∈ (0, 1] that causes the influence of older draws to decay exponentially over time. We hypothesised that this mechanism could increase the relative weight of recent evidence with respect to the total accumulated evidence. Because the forgetting mechanism produces non-integer effective counts, the POMDP is solved on a continuous-valued state grid with step size 0.2, using linear and bilinear interpolation for beliefs and value functions at off-grid states.

#### Belief calculation

As a consequence of forgetting, the sufficient statistics of the belief state are no longer the total number of yellow and blue cards observed, but instead the effective (forgetting-weighted) counts 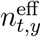 and 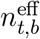 —continuous-valued quantities that reflect a decaying contribution of past draws updated recursively at each timestep. The belief follows the same conjugate Beta-Binomial framework as the normative model (Eq. 3), with these effective counts taking the place of the raw card totals:

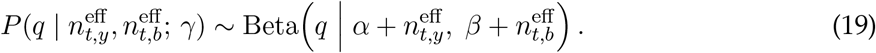

When *γ* = 1, the effective counts coincide with the integer card totals and the expression reduces to the normative posterior (Eq. 3).

#### Transition probability

In the normative model, the sufficient statistics after observing *i* new yellow cards simply accumulate: *n*_*t*+1,*y*_ = *n*_*t,y*_ + *i* and *n*_*t*+1,*b*_ = *n*_*t,b*_ + (*n* − *i*). In the forgetting model, the accumulated evidence is first discounted before the new draw is added. Specifically, when the agent is in state 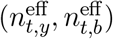 and observes *i* yellow cards in the next draw, the updated effective state is:

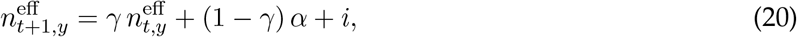

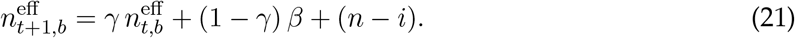

The offsets (1 − *γ*)*α* and (1 − *γ*)*β* ensure that as *γ* → 0 the effective accumulated past evidence decays towards *α* = 1 for the yellow counts and towards *β* for the blue counts.

When *γ* = 1, Eqs. (20–21) reduce to the standard normative accumulation.

The transition probability is computed from the effective past evidence at the moment of transitioning. From state 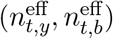, the predictive distribution over the number of yellow cards *i* in the next draw is:

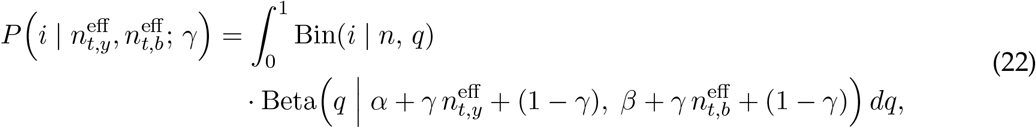

where 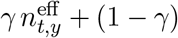 is the effective past yellow evidence after discounting.

### Model fitting

#### Log-likelihood

The negative log-likelihood was used as the objective function for model fitting. At each draw within a trial, the participant either waits or commits to a decision (yellow or blue). The model assigns action probabilities according to the softmax policy *π*(*a* | *n*_*y*_, *n*_*b*_) defined in Eqs. 8–10, where *a* ∈ *{Y, B, W}* corresponds to choosing yellow, choosing blue, or waiting given the observed evidence. Let 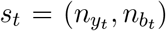 denote the state of evidence at observation *t*, characterised by the number of yellow and blue cards revealed. The total log-likelihood across all observations is

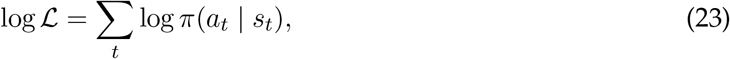

where *a*_*t*_ denotes the action taken by the participant at observation *t*. The negative log-likelihood, − log ℒ, was minimised during parameter estimation.

#### Commit-versus-wait likelihood

As a second check on model performance, we asked whether data simulated from the fitted model reproduces the GLM and GLMM results reported for this dataset [26]. Those regressions predict only whether a participant commits on a given draw or waits; they say nothing about which of the two colours is then chosen. To put the two accounts on an equal footing we therefore refitted the model against a second likelihood that predicts only *p*(commit). This likelihood is used only for the models that enter the comparison with the GLM and GLMM (S1 Appendix, “Commit likelihood and comparison with the GLM”); every other result reported here comes from models fitted against the full likelihood of Eq 23.

Writing *p*_*Y*_, *p*_*B*_ and *p*_*W*_ for the three choice probabilities of the policy *π*(*a* | *n*_*y*_, *n*_*b*_) in Eqs. 8–10, at each observed draw these are collapsed into a single commit probability,

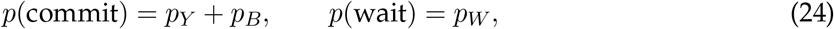

Writing *ã*_*t*_ ∈ {commit, wait} for the participant’s observed action under this grouping, the commit log-likelihood is

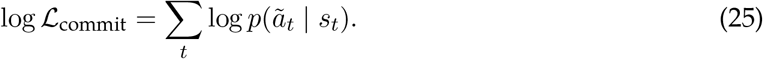

Every candidate model was refitted under this objective rather than having its full-likelihood parameters reused. Maximising log ℒ_commit_ is not exactly the same as maximising log ℒ over *a* ∈ *{Y, B, W}* and summing *p*_*Y*_ and *p*_*B*_ afterwards. We also report the differences in parameters between the full-fit and commit models in S1 Appendix, “Commit likelihood and comparison with the GLM”.

#### Fitting and optimisation procedure

Each participant was fitted individually, yielding a separate set of parameters per subject, by minimising the negative log-likelihood. Everything below applies equally to the full likelihood of Eq 23 and to the commit likelihood of Eq 25.

We considered models including different subsets of the belief bias *β*, the exaggeration factor *E*, the forgetting factor *γ*, the subjective cost *R*_risk_, the temperature *τ*, the lapse rate *ξ*, and the temporal regulation parameters (*ϕ*_min_, *ϕ*_max_, *k, t*_*p*_). This allowed us to examine whether behaviour is better explained by a simpler parameterisation or whether additional parameters are required to capture the observed decision dynamics. Because behavioural dynamics differed across horizons, we fitted each model three ways: to the short-horizon condition only, to the long-horizon condition only, and jointly to both. In the combined case, data from the short- and long-horizon conditions were concatenated in the order obtained and fitted with a single set of parameters per subject.

#### Fitting strategy

Our aim was to understand the decision-making process in humans in terms of the three components introduced above (how evidence is gathered, how it is incorporated into belief, and when to stop and commit): in what specific ways each of these departs from the theoretically optimal strategy, and which mechanisms different individuals rely on to produce those departures. We first carried out a data exploration, running a range of model-agnostic analyses of decision timing and accuracy to identify what kind of departures from optimality the task and the subject data could plausibly support, before committing to any specific mechanism. We then started from the normative POMDP with no additional mechanism and built the model up iteratively: fitting, simulating from the fitted parameters, comparing the simulated and observed behaviour, and running a parameter-recoverability analysis, repeating this cycle each time a systematic mismatch pointed to a candidate mechanism. Each such mismatch motivated one candidate mechanism: belief bias, a recency bias implemented as either transient exaggeration or forgetting of past evidence, a subjective cost of an incorrect choice, or a time-varying regulation of patience and urgency. In addition to building the model bottom-up, we also performed an ablation analysis, evaluating both directions by simplifying a full model top-down and expanding a simple baseline model bottom-up.

We refit the model each time a mechanism was added, and before treating any mechanism as part of the model we required it to pass three checks. First, it had to improve fit over the model without it, under the comparison criteria described below (*Model comparison*). Second, we verified parameter recovery using parameter values obtained from fitting the model to the behavioural data: we used these realistic values to generate synthetic datasets, then refit the model blind with the same procedure to ensure the parameters can be recovered. Third, we simulated the mechanism in isolation to confirm it behaved as intended, for example verifying that the exaggeration factor increased the relative weight of the most recent sample specifically with respect to total evidence, ensuring the implementation matched the intended computation.

Once this process had produced a model complex enough to capture the data, we ran a more precise, personalised analysis to identify the correct combination of mechanisms for each subject individually, rather than assuming the same mechanism set fitted every subject equally well. We fitted the complete set of 78 candidate models spanning every valid combination of mechanisms and horizon condition (Candidate models, below; full set in S1 Appendix, “Full candidate model set”) and asked, independently for each subject, which combination best explained their own behaviour (Personalised (per-subject) model selection, in Results). This full candidate-space comparison, evaluated with multiple criteria (Model comparison, below), also let us confirm that the mechanisms retained by the iterative build-up were both necessary given the full space of simpler and more complex alternatives.

In the forgetting model, *γ* was not searched continuously but evaluated over a grid of candidate values with a step size of 0.02, with each proposed *γ* snapped to the nearest grid point during optimisation. This parameter grid is distinct from the continuous belief grid on which the value function is solved; see the “Forgetting” section). We chose this discretisation due to the substantial computational complexity associated with fitting the forgetting model to the behavioural data.

For the exaggeration and forgetting models, the recency mechanism can enter the fitting procedure at different points, and we compared three procedures to isolate where the distortion is best located. In the first, the candidate model (exaggeration or forgetting) was fitted directly to the participants’ original behavioural responses, so that the recency mechanism is treated as part of the agent’s internal belief updating. In the second, the observations were amplified and the amplified sequences were fitted with the same recency model. In the third, the amplified observations were instead fitted to the normative model, asking whether a distortion at the level of the observed evidence alone–without any change to the updating rule–could account for behaviour. Comparing these procedures allowed us to distinguish whether the recency bias is best captured as a property of belief updating, of the perceived evidence, or of both. In what follows we report, as justified by the model fits, the transient variant of the exaggeration model fitted to the original behavioural data (the first procedure), and the variant of the forgetting model in which forgetting acts on both the agent’s internal belief updating and the observed evidence.

#### Optimisation strategy

Because the negative log-likelihood objective function lacks a closed-form expression, yields no analytical gradients, and contains local optima alongside flat regions, we employed gradient-free global optimisation strategies. We initially used genetic algorithm.

Although genetic algorithms successfully identified the relevant regions of parameter space, they frequently failed to achieve precise local convergence around exact optima. We systematically detected this non-convergence by exploiting the nesting structure across models. Whenever a larger model’s search space encompasses a nested submodel’s configuration, the larger model can reproduce the nested fit exactly by adopting its parameter values. At the true optimum, the fit must therefore satisfy log ℒ_*i*_(*M*) ≥ log ℒ_*i*_(*N*) for every subject *i*, where model *M* nests model *N*. If the larger model fits any subject worse, at least one of the two fits has failed to converge–a condition we define as a “nesting violation.” Under the genetic algorithm optimisation, 66 of the 190 checkable model pairs contained at least one nesting violation.

To resolve these convergence failures, we refitted all models using differential evolution (`scipy.optimize.differential_evolution`) with the `best1bin` strategy, a population size of 15 per parameter, up to 300 iterations, mutation drawn from (0.5, 1.0), a recombination rate of 0.9, and a Sobol-sequence initialisation to cover the search space uniformly. We set the convergence tolerance to zero to ensure every run exhausted its full iteration budget, and enabled a final local polish using L-BFGS-B to refine the best candidate point. Differential evolution demonstrated faster convergence and achieved superior log-likelihood values compared to the genetic algorithms.

Although differential evolution substantially reduced nesting violations, it did not eliminate them entirely. We therefore implemented a second optimisation pass that leveraged the nesting structure directly. For each model and subject, we identified the best-fitting nested neighbour, mapped its parameters into the coordinate space of the larger model, and used this configuration as the search seed for the larger model. By construction, the seed point lies within the larger model’s search space; hence, a converged fit cannot perform worse than the nested model it contains. Because seeding can occasionally cause an optimiser to trap prematurely in a local basin, we executed both seeded and unseeded runs for every subject and retained, for each subject independently, whichever fit achieved the higher log-likelihood. This two-start procedure provides a performance guarantee via the seeded run while protecting against premature convergence. All reported results reflect these finalised, merged fits.

#### Candidate models’ naming

The model names, given in Table 4, indicate the horizon condition and which parameters were free. The first character is `S`, `L`, or `C`, denoting the short, long, or combined horizon condition, respectively. The remaining characters occupy fixed positions following the order `BEXTGRPHCLUK` (Table 5): a letter indicates that the corresponding parameter was free (fitted), whereas a dash (`-`) indicates that it was fixed to its default value or set to zero (excluded). The generative-hazard position is the one exception to this two-way scheme, because the hazard takes three states rather than two: an uppercase `H` means it was free (fitted; either the true generative hazard function, *h*(*t*), or the deterministic deadline hazard function, *g*(*t*)), a lowercase `h` means the mechanism was on but held fixed at *h*(*t*), and a dash means it was held fixed at the deterministic deadline hazard *g*(*t*). The single-letter codes are fixed labels and do not always coincide with the parameter symbol. Note that the exaggeration factor (`E`) and the forgetting factor (`G`) are alternative recency mechanisms and were never free simultaneously.

**Table 4.** The three best-supported models, one per horizon condition. For each free parameter the fitted range is given as (min, max); a single value means the parameter was held fixed at that value rather than fitted, and True/False gives the state of the generative hazard when it was not itself fitted. *N*_params_ counts the free parameters. The full set of candidate models is given in S1 Appendix, “Full candidate model set”.

| Model | $N_{\text{params}}$ | $\beta$ | $E$ | $\xi$ | $\tau$ | $\gamma$ | $R_{\text{risk}}$ | $t_p$ | $H$ | $\phi_{\text{max}}$ | $L$ | $\phi_{\text{min}}$ | $k$ |
| --- | --- | --- | --- | --- | --- | --- | --- | --- | --- | --- | --- | --- | --- |
| SB-XT-RPh---- | 5 | (0.01, 5) | 1 | (0, 1) | (0, 100) | 1 | (–300, 0) | (0, 8) | True | 50 | 0 | –10 | –2 |
| LBE-T-RPhCL-- | 7 | (0.01, 5) | (0.1, 4) | 0 | (0, 100) | 1 | (–300, 0) | (0, 14) | True | (0, 80) | (0, 1) | –10 | –2 |
| C-EXT-RPHC-UK | 9 | 1 | (0.1, 4) | (0, 1) | (0, 100) | 1 | (–300, 0) | (0, 14) | (0, 1) | (0, 80) | 0 | (–30, 0) | (–20, 0) |

**Table 5.** Parameter codes and their corresponding model parameters.

| Code | Parameter | Symbol |
| --- | --- | --- |
| T | Temperature | $\tau$ |
| X | Lapse rate | $\xi$ |
| B | Belief bias | $\beta$ |
| H | Hazard | $H$ |
| R | Subjective cost | $R_{\text{risk}}$ |
| L | Hazard lapse | $L$ |
| E | Exaggeration factor | $E$ |
| G | Forgetting factor | $\gamma$ |
| P | Patience time | $t_p$ |
| C | Maximum regulation | $\phi_{\text{max}}$ |
| U | Minimum regulation | $\phi_{\text{min}}$ |
| K | Regulation slope | $k$ |

For example, `LBEXT-RPHCLUK` denotes a long-horizon model in which every parameter is free except the forgetting factor *γ* (the dash in the `G` position), corresponding to the full exaggeration model with forgetting disabled.

#### Model comparison

For each model, we assessed goodness of fit using the Bayesian Information Criterion (BIC), the Akaike Information Criterion (AIC), and the corrected AIC (AICc); AICc adds a finite sample correction penalty to AIC based on the number of trials per participant to adjust for small sample sizes [32, 33]. Across all criteria, lower values indicate a better fit after penalising for the number of free parameters. Because models were fitted per subject, the criteria were summed across participants for comparison. To facilitate comparison, we report each criterion as a difference relative to the best-fitting model (lowest in BIC score), ΔBIC, ΔAIC, and ΔAICc, so that the best model takes a value of zero and larger values indicate progressively worse fit.

#### GLM Fitting

In [26], the authors fitted a model-agnostic, data-driven generalised linear model (GLM) and general linear mixed model (GLMM) to the decision times of the participants. These include various regressors that influence these choices. We replicated this analysis, in order to compare the fidelity of the predictions of these accounts to our augmented POMDP model. We also applied the GLM and GLMM to data simulated from our models to compare the parameter values.

##### Total Evidence and Evidence Strength Update

Following [26], the total evidence at draw *t, ES*_*t*_, is defined as the absolute difference between the cumulative majority and minority card counts–that is, the net dominance of the leading colour accumulated up to the current draw. Let 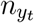 and 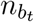 denote the cumulative counts of yellow and blue cards up to draw *t*, respectively, where each draw consists of *n* combinations of yellow and blue cards:

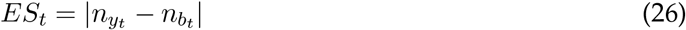

The last evidence strength update Δ*ES*_*t*_ captures how much the total evidence changed from the previous draw to the current one:

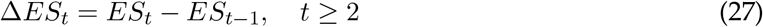

with Δ*ES*_1_ undefined (NaN), since no prior draw exists. A positive Δ*ES*_*t*_ indicates that the leading colour gained further ground, whereas a negative value indicates that the minority colour reduced the evidence gap. This distinction mirrors the relationship between prior expectation and prediction error. A worked example is provided in S1 Appendix, “Example *ES*_*t*_ and Δ*ES*_*t*_ calculations”.

##### GLM Setup

To measure the contribution of total evidence and last-evidence strength update in the decision-making process, we generated simulated datasets using the models fitted to each subject’s behavioural data, matching the number of samples to the actual behavioural data. To account for stochasticity in the simulated datasets, we generated an ensemble of 200 simulated datasets, fitted the GLM to each, and took the average of the resulting coefficients. Following [26], we used a logit-linked GLM predicting the probability of a subject committing to a decision, *p*(commit), using the same regressors and the same names they use (their `totevminus`, `deltaev`, `trial`, `termination`, totevminus**x**term and trial**x**term), written here as:

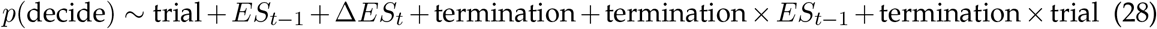

where regressors are defined in Table 6. The output variable *y*_*t*_ is binary:

**Table 6.** GLM regressors used to predict *p*(decide). Regressor names follow del Río and colleagues [26], whose *termination* variable codes the horizon condition; *ES*_*t*−1_ and Δ*ES*_*t*_ are their `totevminus` and `deltaev`, and the two interactions their totevminus**x**term and trial**x**term. Note that this regressor is distinct from the generative termination probability *h*(*t*) of the POMDP, which shares the word but is a different quantity. An intercept is fitted in addition to the six regressors listed.

| Regressor | Description |
| --- | --- |
| `trial` | Number of draws until a decision, $[1, 2, \dots, T]$ |
| `$ES_{t-1}$` | Total evidence at the previous draw |
| `$\Delta ES_t$` | Evidence strength update |
| `termination` | Horizon condition (1 = long, 2 = short) |
| `termination $\times$ $ES_{t-1}$` | Interaction between termination and previous evidence |
| `termination $\times$ trial` | Interaction between termination and draw number |

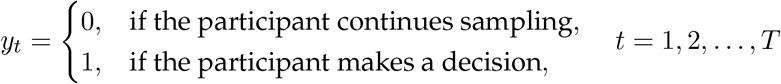

##### GLMM Setup

To account for subject-level variability in decision policies while still estimating group-level contributions of the regressors, we additionally fitted a Generalised Linear Mixed Model (GLMM), similar to [26]. The GLMM extends the GLM Section by including random effects across subjects and interaction terms with the second OCI-R factor score (FA2):

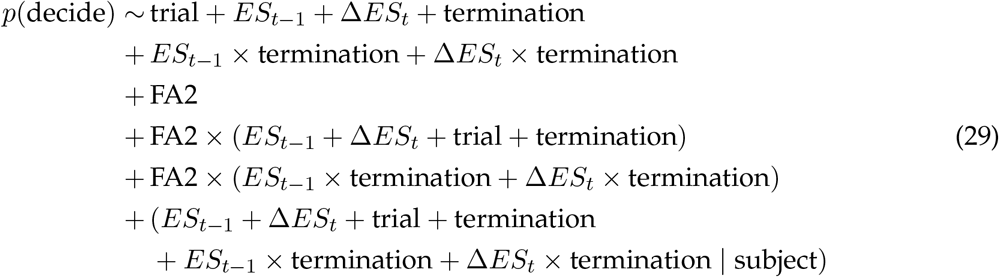

where all regressors are as defined in Table 6, and FA2 denotes the obsessive-compulsive (OC) factor score of [26]. This is the second of the three factors returned by an exploratory factor analysis (maximum likelihood estimation with oblimin rotation, on a heterogeneous correlation matrix) that del Río and colleagues ran on the 210 individual items of the seven psychiatric questionnaires completed by all 105 participants: the Obsessive-Compulsive Inventory-Revised (OCI-R), the revised Padua Inventory (PI-WSUR), the Frost Multidimensional Perfectionism Scale (FMPS), the Intolerance of Uncertainty Scale (IUS), the Beck Depression Inventory II (BDI-II), the Barratt Impulsiveness Scale (BIS), and the State-Trait Anxiety Inventory (STAI). Its highest item loadings come from the two obsessive-compulsive scales, OCI-R and PI-WSUR, while the remaining two factors capture anxious-depressive and intolerance-of-uncertainty/perfectionistic variance respectively. FA2 is therefore a continuous dimension estimated across the whole sample rather than a diagnosis or a clinical cut-off: every participant receives a score irrespective of which group they belong to.

## Results

### Model comparison

#### Short horizon models

Fig 9 and Table 7 summarise the comparison among the top five short-horizon candidate models. `SB-XT-RPh----` is the best-supported model according to BIC, although the four closest alternatives are preferred under AIC and AICc (Table 7). The criteria disagree because they penalise model complexity differently. For short-horizon models, overall fit quality is consistently high across candidates (mean per-subject *R*^2^ = 0.93, as shown in Fig 10b), leading us to prioritise the simplest model with the lowest cumulative BIC score.

**Table 7.** Model selection metrics for the short-horizon candidate models. Rows are ordered by ascending ΔBIC; the best-supported model (`SB-XT-RPh----`, ΔBIC = 0) is shown in bold. *k* = *N*_params_ is the number of free parameters per subject; sum logL and ΔAIC/ΔAICc/ΔBIC are summed across all 105 subjects, each computed from their own observation count (see Model comparison).

| Model | sum logL | $N_{\text{params}}$ | $\Delta AIC$ | $\Delta AICc$ | $\Delta BIC$ |
| --- | --- | --- | --- | --- | --- |
| **SB-XT-RPh----** | **-12791.67** | **5** | **0.00** | **0.00** | **0.00** |
| SB-XTGRPhC--- | -12234.63 | 7 | -694.08 | -676.02 | 93.21 |
| SBEXT-RPh---- | -12603.04 | 6 | -167.26 | -158.94 | 226.39 |
| SBEXT-RPhC--- | -12320.10 | 7 | -523.14 | -505.08 | 264.15 |
| S--XT-RPH--UK | -12347.08 | 7 | -469.18 | -451.12 | 318.11 |

Note: Showing the top 5 of 25 models by BIC. Rows are ordered by ascending $\Delta BIC$ ; the best-supported (BIC-winning) model ( $\Delta BIC = 0$ ) is shown in bold. $\Delta AIC$ , $\Delta AICc$ , and $\Delta BIC$ are all relative to this same BIC-winning model, so a negative $\Delta AIC/\Delta AICc$ means that row is actually preferred over the bolded row on that criterion specifically; $k = N_{\text{params}}$ .

**Fig 9.**
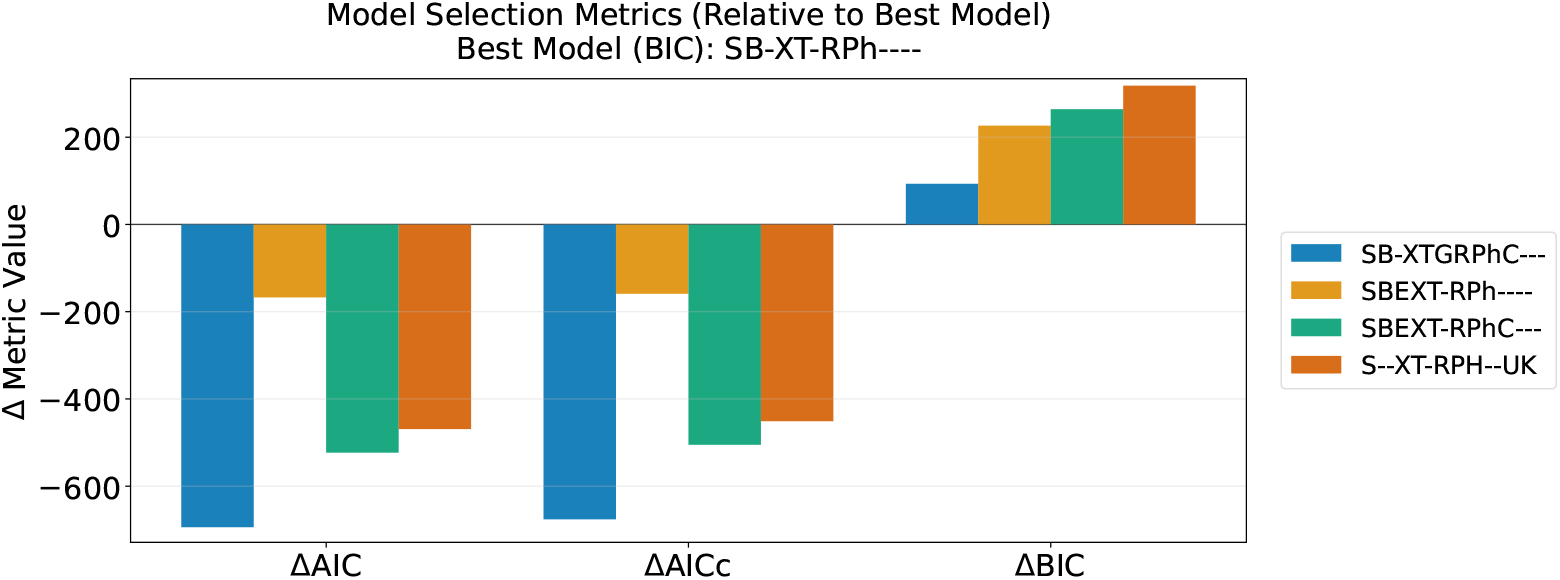
Comparison of short-horizon candidate models. Model selection metrics (ΔAIC, ΔAICc, ΔBIC) for each candidate, relative to the best-supported model; lower is better. `SB-XT-RPh----` is the best-supported model (ΔBIC = 0).

**Fig 10.**
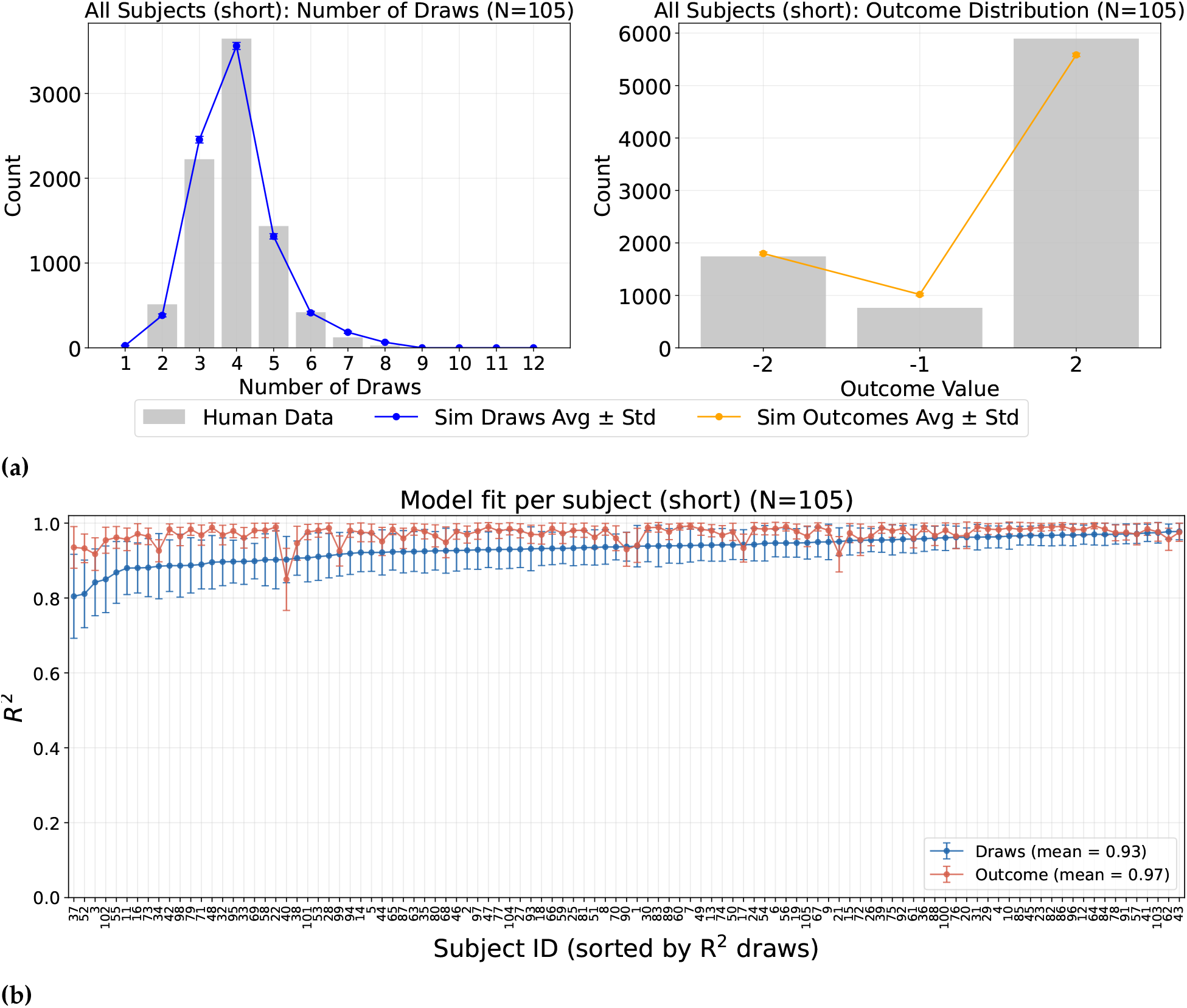
Model fit for SB-XT-RPh---- (short horizon). (a) Human (gray bars) vs. model-simulated (mean *±* SD over simulation ensembles) distributions of number of draws (left) and outcome (right), pooled across all *N* = 105 subjects. (b) Per-subject goodness of fit: *R*^2^ between human and simulated histograms for number of draws (blue) and outcome (red), subjects sorted by draws-*R*^2^; mean *R*^2^ given in the legend.

`SB-XT-RPh----` contains five free parameters estimated per subject (Table 4): belief bias *β*, lapse rate *ξ*, temperature *τ*, subjective risk cost *R*_risk_, and patience time *t*_*p*_. Notably, this winning model includes neither recency mechanism. This is consistent with expectations for the short horizon condition, where participants are less likely to exhibit forgetting or exaggeration and instead keep count of the evidence. The generative hazard mechanism remains active (*H* = True), reflecting participant awareness of the stochastic deadline, as evidenced by the short-horizon draw distribution (Fig 2). Furthermore, the inclusion of *R*_risk_ captures an elevated subjective penalty for incorrect choices, providing an explanation for why participants occasionally miss the deadline. The best-performing forgetting-based and exaggeration-based variants trail `SB-XT-RPh----` by ΔBIC = 93.2 and 226.4, respectively.

In `SB-XT-RPh----`, the shape parameters of the temporal regulation function are fixed (*ϕ*_max_ = 50, *ϕ*_min_ = −10, and slope *k* = −2), leaving only its midpoint, patience time *t*_*p*_, to be fitted per subject. These fixed values were obtained by taking the mean of each parameter across subject-level fits from a broader candidate model where they were left free (see the “Candidate models’ naming” section). To verify that this choice was near-optimal, we held all other fitted parameters constant for each subject while sweeping each of the three shape parameters across its original search range, evaluating the population-summed log-likelihood at each value (S1 Appendix, “Parameter and model recovery and sensitivity”). All three parameters landed at or near their population optima (*k* = −2 matched exactly; *ϕ*_max_ = 50 vs. an optimum of 48, costing 4.7 BIC points; *ϕ*_min_ = −10 vs. an optimum of −10.5, costing 6.5 BIC points). We therefore selected `SB-XT-RPh----` for subsequent analyses. Detailed fits for the best-supported models are presented below, and parameter recovery results are detailed in S1 Appendix, “Parameter and model recovery and sensitivity”.

Figure 10 summarises model fit for `SB-XT-RPh----`, the best-supported short-horizon model (Table 7); parameter and model recovery results are shown in S1 Appendix, “Parameter and model recovery and sensitivity”.

#### Long horizon models

Fig 11 and Table 8 summarise the comparison among the top 5 long-horizon candidate models. `LBE-T-RPhCL--` is consistently the best-fitting model across all comparison metrics (Table 8), outperforming the best-supported forgetting variant by ΔBIC = 91.1. `LBE-T-RPhCL--` has seven free parameters (Table 4): belief bias *β*, the exaggeration factor *E*, temperature *τ*, the subjective cost *R*_risk_, patience time *t*_*p*_, maximum regulation *ϕ*_max_, and hazard-lapse rate *L*. The winning recency mechanism here is exaggeration. The temporal regulation function’s ceiling (*ϕ*_max_) and hazard-lapse rate (*L*) are both fitted per subject, while its floor (*ϕ*_min_ = −10) and slope (*k* = −2) remain fixed at the same values used in the short-horizon model. As in the short horizon, we checked how close these fixed values were to optimal by holding every subject’s other fitted parameters constant and sweeping each one over the range used when it was fitted freely, evaluating the resulting population-summed log-likelihood (S1 Appendix, “Parameter and model recovery and sensitivity”). Both values proved optimal (*ϕ*_min_ = −10 and *k* = −2; ΔBIC = 0.0 for each). We therefore carried this model forward to the combined-vs-separate comparison below; detailed fitting for `LBE-T-RPhCL--` is presented below, and parameter recovery is shown in S1 Appendix, “Parameter and model recovery and sensitivity”.

**Table 8.** Model selection metrics for the long-horizon candidate models. Rows are ordered by ascending ΔBIC; the best-supported model (`LBE-T-RPhCL--`, ΔBIC = 0) is shown in bold.

| Model | sum logL | $N_{\text{params}}$ | $\Delta\text{AIC}$ | $\Delta\text{AICc}$ | $\Delta\text{BIC}$ |
| --- | --- | --- | --- | --- | --- |
| **LBE-T-RPhCL--** | **-16585.75** | **7** | **0.00** | **0.00** | **0.00** |
| LB--TGRPhCL-- | -16631.28 | 7 | 91.06 | 91.06 | 91.06 |
| LBEXT-RPhCL-- | -16486.05 | 8 | 10.60 | 17.14 | 460.13 |
| LB--T-RPhCL-- | -17181.29 | 6 | 981.07 | 975.36 | 531.54 |
| LB-XTGRPhCL-- | -16545.84 | 8 | 130.18 | 136.72 | 579.71 |

Note: Showing the top 5 of 31 models by BIC. Rows are ordered by ascending $\Delta\text{BIC}$ ; the best-supported (BIC-winning) model ( $\Delta\text{BIC} = 0$ ) is shown in bold. $\Delta\text{AIC}$ , $\Delta\text{AICc}$ , and $\Delta\text{BIC}$ are all relative to this same BIC-winning model, so a negative $\Delta\text{AIC}/\Delta\text{AICc}$ means that row is actually preferred over the bolded row on that criterion specifically; $k = N_{\text{params}}$ .

**Fig 11.**
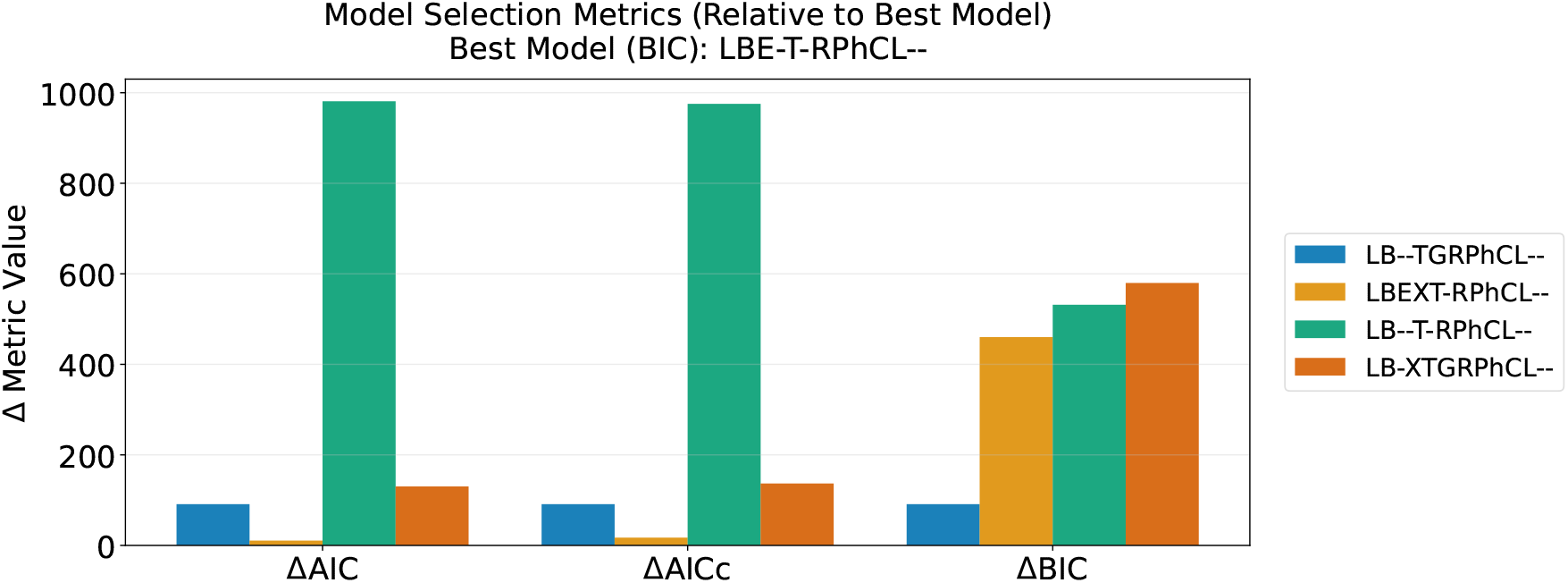
Comparison of long-horizon candidate models. Model selection metrics relative to the best-supported model. `LBE-T-RPhCL--` is the best-supported model (ΔBIC = 0).

Fig 12 summarises model fit for `LBE-T-RPhCL--`, the best-supported long-horizon model (Table 8); parameter and model recovery results are shown in S1 Appendix, “Parameter and model recovery and sensitivity”. Per-subject ensemble fits for the individual subjects with the most extreme fitted exaggeration factor, under this model and under `C-EXT-RPHC-UK`, are shown in S1 Appendix, “Example individual subject fits”.

**Fig 12.**
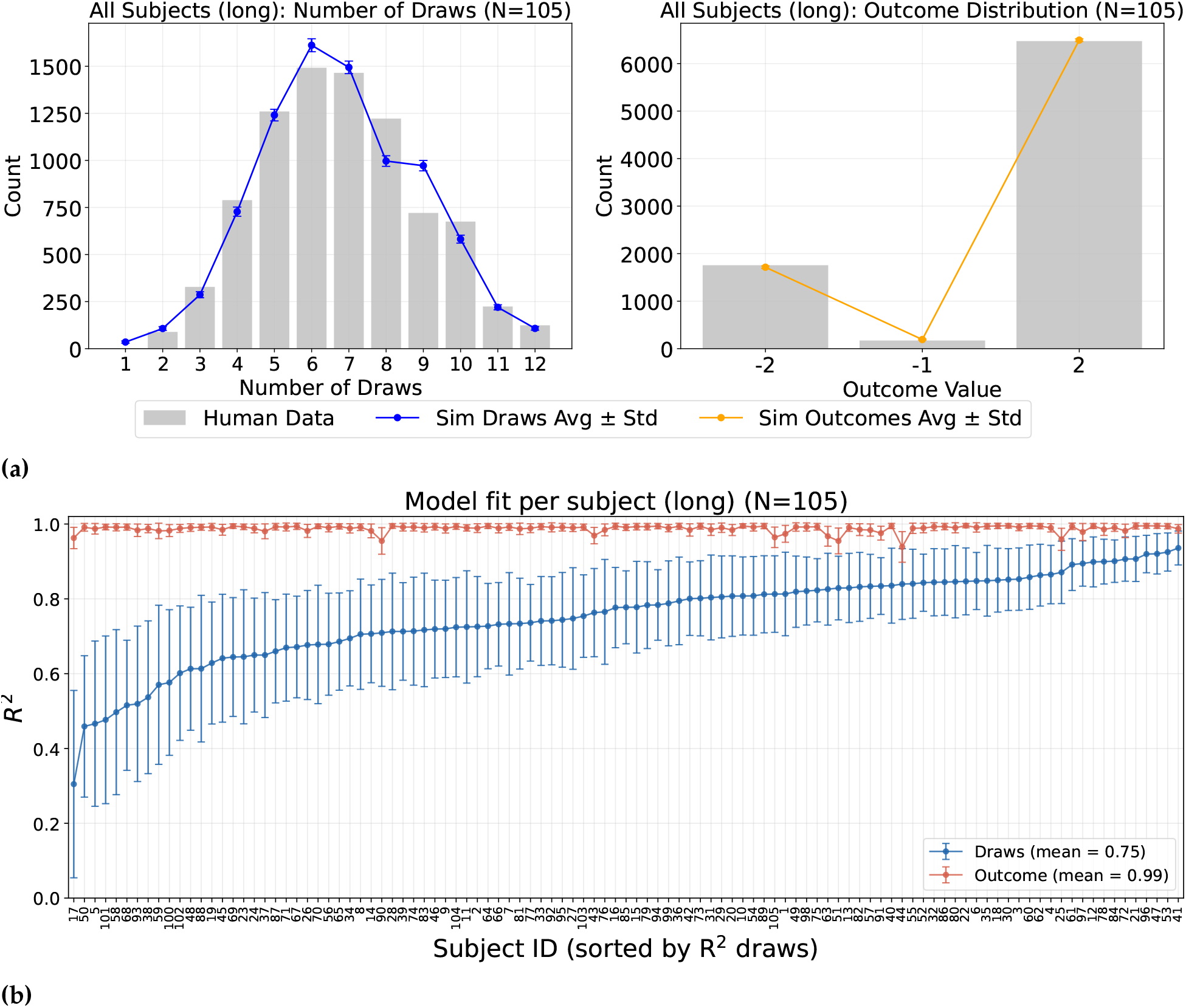
Model fit for LBE-T-RPhCL-- (long horizon). (a) Human vs. model-simulated distributions of number of draws (left) and outcome (right), pooled across all *N* = 105 subjects, as in Fig 10a. (b) Per-subject *R*^2^ goodness of fit for draws and outcome, as in Fig 10b.

#### Combined horizon models

Fig 13 and Table 9 present the comparison among the combined-horizons candidate models. Here the two model comparison criteria agree: `C-EXT-RPHC-UK` has the lowest (best) BIC and the lowest AIC (Table 9), ahead of the next candidate, `C-E-T-RPHC-UK` (ΔBIC = 315.9). The best forgetting-type candidate, `CB-XTGRPHCLUK`, trails by ΔBIC = 2559.5, so unlike the horizon-specific fits the combined horizon does not favour a forgetting mechanism; the margin against forgetting is far wider here than in either horizon-specific fit, where the best forgetting variant trails by only 93.2 and 91.1. We selected `C-EXT-RPHC-UK` for the subsequent comparison between the combined and separate fits below. `C-EXT-RPHC-UK` has nine free parameters (Table 4): the lapse rate *ξ*, temperature *τ*, the subjective cost *R*_risk_, patience time *t*_*p*_, the exaggeration factor *E*, the temporal regulation floor *ϕ*_min_, ceiling *ϕ*_max_ and slope *k*, and the hazard *H*. Unlike either horizon-specific model it fits no belief bias, and unlike `SB-XT-RPh----` it retains the exaggeration factor; the whole temporal regulation function is fitted per subject here rather than held at fixed values, and the hazard is fitted rather than fixed.

**Table 9.** Model selection metrics for the combined-horizons candidate models. Rows are ordered by ascending ΔBIC; the best-supported (BIC-winning) model (`C-EXT-RPHC-UK`, ΔBIC = 0) is shown in bold. ΔAIC, ΔAICc, and ΔBIC are all relative to this same model, which is also the AIC and AICc winner, so all three criteria agree here and no Δ is negative.

| Model | sum logL | $N_{\text{params}}$ | $\Delta\text{AIC}$ | $\Delta\text{AICc}$ | $\Delta\text{BIC}$ |
| --- | --- | --- | --- | --- | --- |
| **C-EXT-RPHC-UK** | **-30373.89** | **9** | **0.00** | **0.00** | **0.00** |
| C-E-T-RPHC-UK | -30886.02 | 8 | 814.26 | 809.70 | 315.93 |
| C--XT-RPHC-UK | -30892.12 | 8 | 826.46 | 821.90 | 328.13 |
| CB-XT-RPHC-UK | -30811.89 | 9 | 876.00 | 876.00 | 876.00 |
| CBEXT-RPhCLUK | -30474.87 | 10 | 411.97 | 417.04 | 910.30 |

Note: Showing the top 5 of 22 models by BIC. Rows are ordered by ascending $\Delta\text{BIC}$ ; the best-supported (BIC-winning) model ( $\Delta\text{BIC} = 0$ ) is shown in bold. $\Delta\text{AIC}$ , $\Delta\text{AICc}$ , and $\Delta\text{BIC}$ are all relative to this same BIC-winning model, so a negative $\Delta\text{AIC}/\Delta\text{AICc}$ means that row is actually preferred over the bolded row on that criterion specifically; $k = N_{\text{params}}$ .

**Fig 13.**
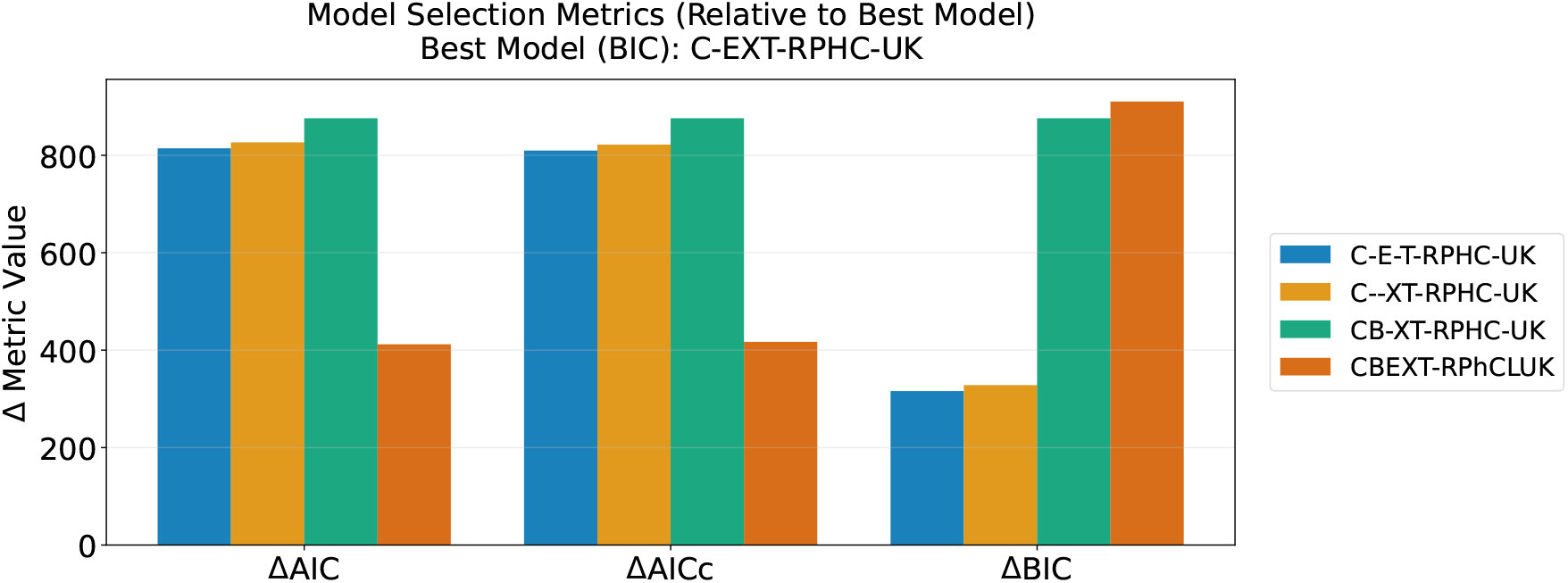
Comparison of combined-horizons candidate models. Model selection metrics relative to `C-EXT-RPHC-UK`, which is the best-supported candidate on all three criteria (ΔAIC = ΔAICc = ΔBIC = 0). A negative bar would mark a candidate preferred over it on that criterion specifically; none occurs here, so AIC, AICc and BIC agree.

Fig 14 summarises model fit for `C-EXT-RPHC-UK`, the best-supported combined-horizons model (Table 9), fit jointly to short- and long-horizon trials. The model achieves a mean per-subject draws-*R*^2^ of 0.82 and outcome-*R*^2^ of 0.99 (Fig 14b), between the short- and long-horizon fits on draws. Parameter and model recovery results are shown in S1 Appendix, “Parameter and model recovery and sensitivity”.

**Fig 14.**
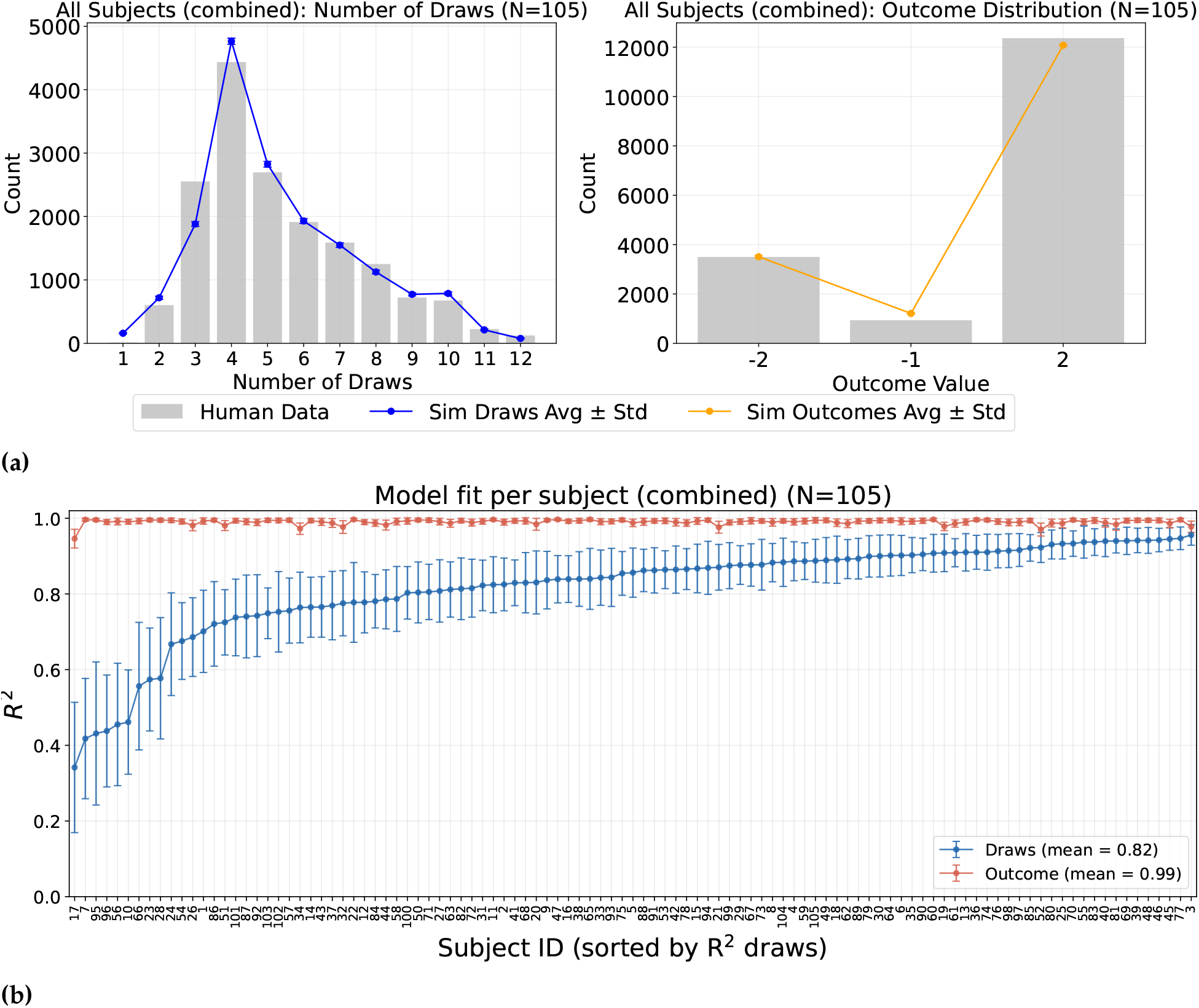
Model fit for C-EXT-RPHC-UK (combined horizons). (a) Human vs. model-simulated distributions of number of draws (left) and outcome (right), pooled across all *N* = 105 subjects and both horizons, as in Fig 10a. (b) Per-subject *R*^2^ goodness of fit for draws and outcome, as in Fig 10b.

#### Short + long vs. combined models

Fig 15 and Table 10 compare the best-performing separate short- and long-horizon models (`SB-XT-RPh----` + `LBE-T-RPhCL--`) against the best-performing combined-horizons model (`C-EXT-RPHC-UK`). The separate fit is consistently better across all comparison metrics than the combined-horizons fit.

**Table 10.** Model selection metrics: best short + long horizon models vs. the best combined-horizons model. *k* = *N*_params_ is the per-subject free-parameter count, summed across the short and long fits for the separate-model row; sum logL and AIC/AICc/BIC are summed across all 105 subjects, each computed from their own observation count.

| Model | sum logL | $N_{\text{params}}$ | AIC | AICc | BIC |
| --- | --- | --- | --- | --- | --- |
| SB-XT-RPh---- + LBE-T-RPhCL-- | -29377.42 | 12 | 61274.85 | 61318.09 | 66389.78 |
| C-EXT-RPHC-UK | -30373.89 | 9 | 62637.78 | 62660.45 | 67122.78 |

**Fig 15.**
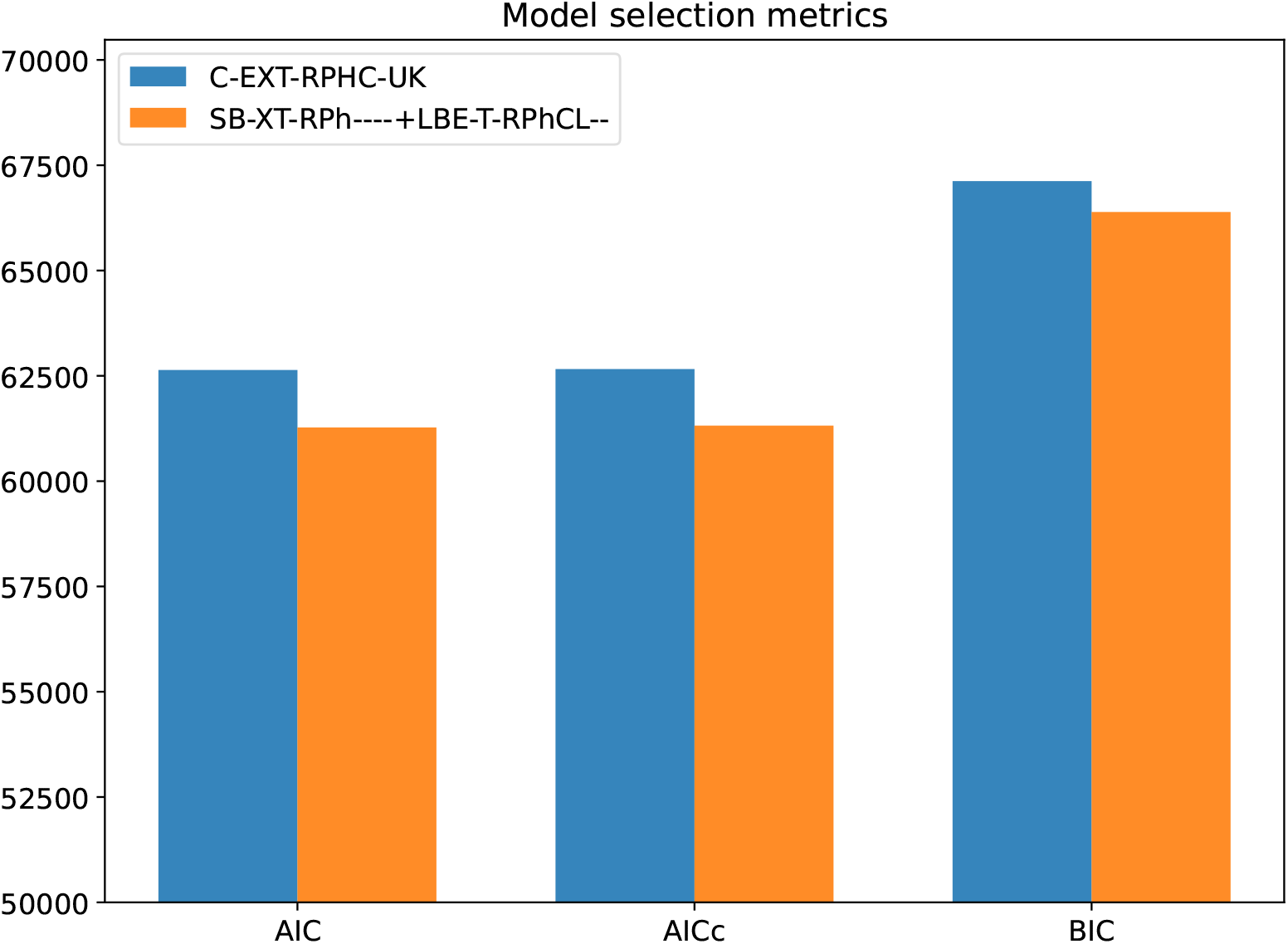
Separate (short + long) vs. combined model fits. Raw AIC, AICc, and BIC for the summed separate fits (`SB-XT-RPh----` + `LBE-T-RPhCL--`, orange) against the combined fit (`C-EXT-RPHC-UK`, blue). Lower is better on all three, so the separate fits are preferred on every criterion.

#### Model fit differs across horizons

Although all three models achieved a good overall fit (mean individual *R*^2^ = 0.93 for the short horizon, 0.75 for the long horizon, and 0.82 for combined horizons; Fig 10b, Fig 12b, Fig 14b), they did not fit equally well. One potential cause is that the long horizon features more reachable states than the short horizon. By task design, both conditions provided the same number of games (80 each): at each draw *t*, the belief state (*n*_*y*_, *n*_*b*_) can take 5*t* + 1 distinct values, yielding 540 reachable states across the 14 draws of the long horizon compared with only 189 across the 8 draws of the short horizon. More specifically, model fit is heavily influenced by states at the decision boundary, where the fitted policy transitions between waiting and committing. Pooling states adjacent to the boundary across all 105 subjects’ fitted policies yields 8 such states for the short horizon versus 20 for the long horizon. Each of the 8 short horizon boundary states was visited at least 15 times across the sample. In contrast, half of the 20 long horizon boundary states (concentrated from draw 5 onward, near and past the hazard onset at draw 10) were never visited. Consistent with this, fits for the short horizon were high for nearly all subjects, ranging from 0.80 to almost perfect. Fits for the long horizon exhibited greater variability, with *R*^2^ values ranging from approximately 0.31 for subjects with the worst fit to 0.94 for subjects with the best fit. Subject 17, who had the worst fit, reflects noisy distribution of data in the long horizon rather than a failure of the model: their long horizon draw distribution is scattered erratically across draws 2 to 14 with no clear structure. In contrast, this same subject displays clear behavioural dynamics in the short horizon, allowing the short horizon model to fit their data reliably as shown in S1 Appendix, “Example individual subject fits”.

#### The cost of departing from optimality

The model comparison above establishes which mechanisms the data require, but not what those mechanisms cost the subjects who use them. We therefore asked how far below the normative optimum the subjects and their fitted models fall, by scoring every policy against the objective payoff of the task. We constructed the normative agent by returning every departure to its neutral value (*E* = 1, *γ* = 1, *R*_risk_ = 0, *L* = 0, and Φ(*t*) ≡ 0), giving it the task’s own prior over the generative probability (S1 Appendix, “Generative probability and card sequence generation”) in place of the fitted belief bias *β*, with a near-deterministic policy (*τ* = 10^−8^, *ξ* = 0), and kept the generative hazard active. Each policy then replayed that subject’s own games up to the predetermined deadline. Human performance was read from the recorded outcomes.

Subjects fell short of the achievable payoff in every horizon condition (Fig 16a). In the short horizon they earned 71.8 of a possible 95.8 points, a shortfall of 24.0 points (SD 15.5, median 24.0, IQR 15 to 33; Wilcoxon *p* = 1.1 *×* 10^−17^), corresponding to 70.2% correct choices against 79.3%. In the long horizon they earned 88.2 of 131.8 points, a shortfall of 43.6 points (SD 12.5, median 43.0, IQR 35 to 51; *p* = 5.7 *×* 10^−19^) from which no subject escaped, corresponding to 77.1% correct against 90.9%. Pooling both conditions, the shortfall is 67.4 points (SD 20.3) over 160 games. Only 7 of 105 subjects exceeded the normative agent in the short horizon, which is expected on a finite sample given that the normative policy maximises expected rather than realised reward.

**Fig 16.**
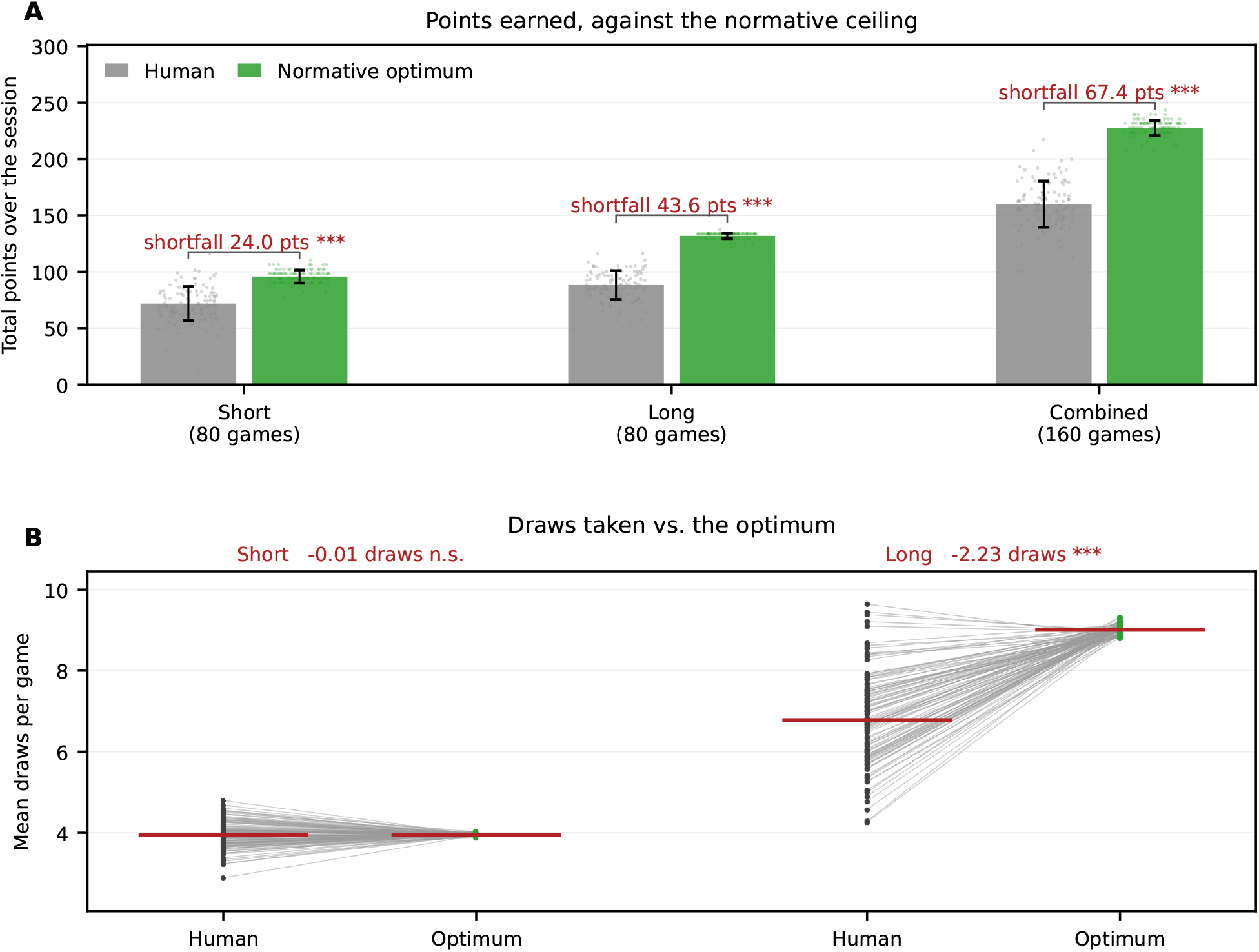
The cost of departing from optimality. (a) Total points earned over the session, for the subjects and the normative optimum, in each horizon condition. Bars give the mean across the 105 subjects with the between-subject standard deviation; each dot is one subject. The shortfall annotated above each group is the mean of the per-subject difference between the normative optimum and the subject. (b) Mean draws taken per game, with each subject’s own value joined to the normative agent’s value on the same card sequences. In the short horizon the two coincide, whereas in the long horizon almost every subject falls below the optimum. Red bars mark group means. Asterisks give paired Wilcoxon signed-rank tests across the 105 subjects (*n*.*s. p >* 0.05, ^∗^*p <* 0.05, ^∗∗^*p <* 0.01, ^∗∗∗^*p <* 0.001); in (a) the bracket compares the subjects with the normative optimum and is labelled by the shortfall it measures, and in (b) the annotation compares human and normative draw counts.

The shortfall does not reflect the same behaviour in the two conditions. In the long horizon, subjects undersampled systematically, taking 6.78 draws against the normative 9.01 (difference − 2.23, *p* = 1.3 *×* 10^−18^), with 100 of 105 subjects below the normative agent. In the short horizon, they took 3.94 draws against the normative 3.95 (Fig 16b), a difference of 0.01 draws that is indistinguishable from zero (*p* = 0.67, with 54 of 105 below). Subjects in the short horizon model (`SB-XT-RPh----`) therefore sampled as much as they should have while still losing 9 percentage points of accuracy. We detected at least two reasons: 1) they mostly misallocated draws across trials, stopping earlier than the normative agent on 29.1% of games and later on 21.1%, which left 9.5% of games undecided at the deadline ( ≈ 7.5 points); and 2) they occasionally chose against their own evidence, on 6.4% of games ( ≈ 1.5 points). On the other hand, the long horizon model (`LBE-T-RPhCL--`) does not require a lapse parameter as it is the condition in which the deadline leaves room to sample further, and it is there that premature commitment dominates. Measured against the optimum, these subjects therefore gather little evidence in the long horizon and the right amount in the short one.

#### Shared mechanisms across horizon conditions

Across both horizon conditions, the best-supported models retained a belief bias, a subjective risk cost on incorrect decisions, patience time, and a temperature parameter (`SB-XT-RPh----` and `LBE-T-RPhCL--`; Tables 7 and 8). They differed in recency: `LBE-T-RPhCL--` fitted an exaggeration factor, whereas `SB-XT-RPh----` fitted neither recency mechanism. The long horizon model additionally required a hazard lapse parameter fitted per subject, consistent with some participants confusing horizon conditions. Fitting the two horizon conditions separately provided a consistently better account than a single model fitted jointly across both (Table 10). This allows us to test whether a given subject’s short and long horizon strategies reflect a shared trait mechanism (where a subject with a high parameter value in one horizon also has a high value in the other) or independent horizon specific strategies. For each of the four shared parameters, we calculated the Spearman rank correlation across all 105 subjects between their short horizon and long horizon fitted values (Fig 17, which also reports Pearson correlations for reference). Two parameters showed significant positive correlations: temperature *τ* (*ρ* = 0.493, *p <* 0.0001) and subjective cost of error *R*_risk_ (*ρ* = 0.567, *p <* 0.0001). Neither patience (*ρ* = 0.136, *p* = 0.168) nor belief bias (*ρ* = 0.054, *p* = 0.581) correlated across horizons. Therefore, subjects maintain a stable tendency across horizons for risk and choice stochasticity (reflected by the temperature parameter), but adjust their patience and prior beliefs separately for each horizon, in addition to incorporating mechanisms such as recency in the long horizon.

**Fig 17.**
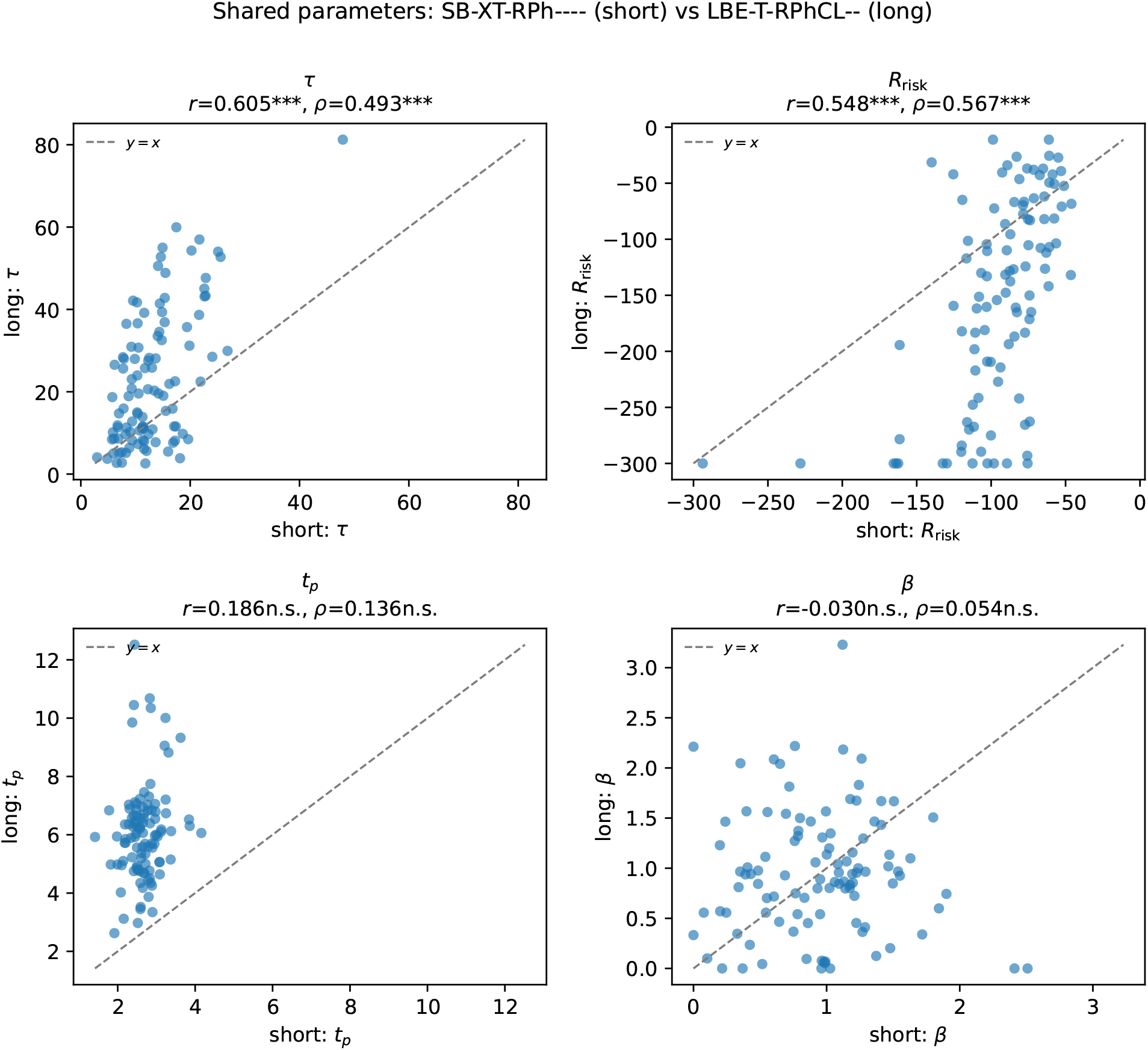
Per-subject correlation of shared parameters between SB-XT-RPh---- (short) and LBE-T-RPhCL-- (long). Each point represents one subject. Each panel depicts one of the four parameters shared between the two independently fitted models, with a dashed line marking equality (*y* = *x*). Each panel title reports the Pearson correlation coefficient *r* (linear agreement) and the Spearman rank correlation coefficient *ρ* (rank agreement; the statistic reported in the main text), alongside significance levels (^∗^*p <* 0.05, ^∗∗^*p <* 0.01, ^∗∗∗^*p <* 0.001).

#### Distinguishing exaggeration from forgetting as the recency mechanism

The two candidate implementations of recency competed directly within every horizon condition, since each candidate model fitted either the exaggeration factor or the forgetting factor but never both. Within the long horizon condition, the two mechanisms are close in BIC scores, whereas in the short horizon condition the runner up is forgetting, but `SB-XT-RPh----` has no recency mechanism (Tables 7 and 8). These margins are too small to separate a transient distortion of how new evidence is weighted from a persistent loss of old evidence. We therefore carried out GLM and GLMM analyses on simulated data from these models. We observed that the exaggeration mechanism from `LBE-T-RPhCL--`, combined with `SB-XT-RPh----`, better reproduces the GLM and GLMM results from human data; see the “Correspondence with GLM-described behaviour” section. On the other hand, as shown in Fig. L, the forgetting model tends to disagree on the relative importance of the most recent evidence strength update relative to total evidence. This was also quite clear when the models were fit exclusively to the commit decision; see Table E in S1 Appendix. Moreover, pooling the conditions separates them decisively: the best candidate based on forgetting trails `C-EXT-RPHC-UK` by more than 2500 BIC (Table 9). Exaggeration is therefore the better-supported implementation once the conditions are pooled, and the one that reproduces the regression signature more faithfully. Within the long horizon taken alone, however, the two cannot be separated, and neither can be separated from having no recency mechanism at all: subjects divide almost evenly between the three when each is allowed their own best-fitting structure, and a random-effects comparison over those fits returns protected exceedance probabilities close to 1*/*3 for all three families. That analysis is reported in the “Personalised model selection per subject” section below, and `LBE-T-RPhCL--` should be read accordingly as the best single description of the sample rather than as evidence that the participants exclusively use exaggeration.

#### Testing recency and urgency hypotheses

Two primary mechanisms have been proposed to explain the information gathering behaviour in OCD: attenuated urgency and mis-weighted recent evidence [25, 26]. To test whether one or a combination of both explains choice behaviour in OCD, we isolated each mechanism by fitting four models per horizon condition that differed solely in which mechanism was active, keeping all other parameters fitted across all four cells and the hazard state identical (Table 11). Removing temporal regulation while retaining exaggeration is by far the most damaging omission in every horizon condition, costing 9191.8 BIC in the short horizon, 21436.9 in the long horizon, and 24650.0 in the combined fit. Removing exaggeration while retaining temporal regulation costs 531.5 BIC in the long horizon and 328.1 in the combined fit. In the short horizon, removing exaggeration improves model fit by 226.4 BIC, as the extra parameter is not justified by the minor log likelihood gain (188.6 versus 595.5 in the long horizon and 518.2 in the combined fit). When standing alone in the long horizon, the exaggeration model performs worse than a model with neither mechanism by 288.2 BIC, demonstrating that amplifying recent evidence without temporal regulation actively degrades fit. In contrast, for short and combined horizons, exaggeration alone outperforms the null model by 402.8 and 446.8 BIC, respectively. Exaggeration nonetheless earns its place on a different criterion: without it the model does not capture how the importance of the most recent evidence update rises relative to total evidence. Both mechanisms are therefore jointly necessary to capture these distinct aspects of OCD behaviour. Overall, these results show that both mechanisms are required to account for information gathering, though with distinct relative contributions.

**Table 11.** Isolating the contribution of recency misweighting and temporal regulation. One quartet per horizon condition, each anchored on the winning model for that horizon and varying only the two mechanisms. Within a quartet, every other free parameter is fitted across all four cells and the hazard state is kept identical throughout, so differences in BIC between cells are attributable to the mechanism alone (summed across subjects as throughout). The background parameter set differs between quartets because each is anchored on a different winning model. Temporal regulation means patience time *t*_*p*_ is fitted and Φ(*t*) is active; switching it off sets *ϕ*_max_ = *ϕ*_min_ = *k* = 0, so Φ(*t*) = 0 at every draw. Exaggeration means the exaggeration factor *E* is fitted; switching it off fixes *E* = 1. BIC values are given relative to the best cell of each horizon. Removing temporal regulation is far more damaging than removing exaggeration in all three horizon conditions. Exaggeration earns its place in the long and combined horizons (removing it costs 531.5 and 328.1 BIC), but not in the short horizon, where adding it on top of temporal regulation is 226.4 BIC *worse* than leaving it out. All of these models except `C-EXT-R-H`, which was fitted only for this comparison, belong to the candidate set compared above.

| Horizon | Mechanisms present | Model | $N_{\text{params}}$ | $\Delta\text{BIC}$ |
| --- | --- | --- | --- | --- |
| Short | Temporal regulation only | SB-XT-RPh---- | 5 | **0.0** |
| Short | Exaggeration + temporal regulation | SBEXT-RPh---- | 6 | 226.4 |
| Short | Exaggeration only | SBEXT-R-h---- | 5 | 9418.1 |
| Short | Neither | SB-XT-R-h---- | 4 | 9821.0 |
| Long | Exaggeration + temporal regulation | LBE-T-RPhCL-- | 7 | **0.0** |
| Long | Temporal regulation only | LB--T-RPhCL-- | 6 | 531.5 |
| Long | Neither | LB--T-R-h-L-- | 4 | 21148.7 |
| Long | Exaggeration only | LBE-T-R-h-L-- | 5 | 21436.9 |
| Combined | Exaggeration + temporal regulation | C-EXT-RPHC-UK | 9 | **0.0** |
| Combined | Temporal regulation only | C--XT-RPHC-UK | 8 | 328.1 |
| Combined | Exaggeration only | C-EXT-R-H---- | 5 | 24650.0 |
| Combined | Neither | C--XT-R-H---- | 4 | 25096.8 |

#### Personalised model selection per subject

The comparisons above select a single fixed model structure for each horizon condition. Allowing each subject their own best fitting structure instead improved cumulative BIC across all horizon conditions, with similar margins in the short and long horizons (ΔBIC = 470.4 and 469.8, respectively) and a larger improvement in the combined condition (ΔBIC = 707.3, with 96 of 105 subjects showing improvement). `C-EXT-RPHC-UK` wins on summed BIC but is the best model for only 9 of 105 subjects, while C–XT-RPH–UK is best for 29. The aggregate winner is therefore carried by large improvements in a minority rather than by being the most common account (S1 Appendix, “Protected exceedance probabilities over the individualised models”). These improvements do not, however, compensate for the penalty of selecting a structure per subject: penalising each subject by one additional parameter for their model choice leaves personalised models behind the fixed model in the short and long horizons, and only at almost break-even in the combined horizon. Therefore, we report a single fixed structure per horizon condition throughout. Complete details of this procedure, full comparisons, and parameter accounting are provided in S1 Appendix, “Personalised per-subject model selection”.

Personalisation does, however, make clear that the evidence for a recency mechanism is weaker than a single winning structure suggests, and that this is where subjects differ most. Assigning each subject to the recency family containing their own best-fitting model splits the long horizon almost evenly: 37 of 105 subjects are best fitted by an exaggeration model, 36 by a forgetting model, and 32 by a model with no recency mechanism at all. A random-effects comparison over the same individualised fits reaches the same conclusion. Summing evidence within each family, the estimated population frequencies in the long horizon are 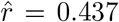 for exaggeration, 0.327 for forgetting and 0.235 for neither, but the Bayesian omnibus risk is BOR = 0.872, which pulls all three protected exceedance probabilities towards 1*/*3 (0.406, 0.303 and 0.291). The long-horizon data therefore do not identify which recency mechanism the population uses, and `LBE-T-RPhCL--` should be read as the best single description of the sample rather than as evidence that exaggeration is the mechanism its subjects employ. The other two horizon conditions are less ambiguous: models with no recency mechanism take 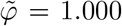 in the short horizon and 0.684 in the combined fit, against 0.316 for exaggeration and 0.000 for forgetting. Full tables are given in S1 Appendix, “Protected exceedance probabilities over the individualised models”. Whether the two mechanisms distinguish different kinds of subject is a further question to which the answer is largely negative. The long-horizon classes are not cleanly separated: the median subject is assigned to their family by only 4.5 BIC over the runner-up, and 24 of 105 subjects by less than 2. Nor do the classes track diagnosis or symptom severity. The proportion of subjects best fitted by exaggeration did not differ between the OCD patients and the rest of the sample in any horizon condition (long horizon 6*/*29 against 31*/*76, Fisher *p* = 0.068; short *p* = 1.000; combined *p* = 0.823), nor between the 20 most and 20 least compulsive subjects (*p* = 0.096, 1.000 and 1.000). Treating the preference as continuous rather than categorical, by taking each subject’s BIC difference between their best forgetting and best exaggeration model, gives one nominally significant association in six: in the long horizon a stronger preference for forgetting accompanies a higher obsessive-compulsive factor score (*ρ* = −0.192, *p* = 0.049) and OCI-R total (*ρ* = −0.190, *p* = 0.052), while the short horizon shows the opposite sign (*ρ* = +0.182, *p* = 0.063) and the combined fit nothing (*ρ* = −0.018, *p* = 0.852). Neither the categorical nor the continuous analysis survives correction, and the reversal between horizon conditions argues against reading the long-horizon result as a subgroup effect.

#### Correspondence with GLM-described behaviour

To check whether the fitted POMDP parameters relate to the same regressors the GLM uses to describe decision timing, and to keep this comparison on the same footing as the GLM (which predicts only whether a participant commits to a decision or continues waiting, not which option is chosen), we fitted the POMDP using the commit-vs-wait likelihood defined in the “Commit-versus-wait likelihood” section. The separate best short- and long-horizon commit-fit models, summed, beat the GLM on every fixed-model metric (AIC and BIC; S1 Appendix, “Commit likelihood and comparison with the GLM”). We simulated an ensemble of choice sequences from each subject’s own best-fitting commit-POMDP parameters, refit the GLM/GLMM to each simulated subject, and correlated each subject’s fitted POMDP parameters with their ensemble-averaged, model-simulated GLM coefficients. The refit GLMM captured the same overall regression signature as the human data, including the same significant terms (Fig I in S1 Appendix, Table E in S1 Appendix), and subject-level GLM coefficients fit to model-simulated choices correlated strongly with those fit to the real human choices across all seven regressors (*r* = 0.67–0.97; Fig J in S1 Appendix). At the parameter level, the exaggeration factor correlated significantly and negatively with the total-evidence regressor (*ES*_*t*−1_; *r* = −0.34, *p <* 0.001), consistent with its role as a mechanism that shifts weighting toward the most recent evidence (Fig K in S1 Appendix). We also correlated each subject’s full-fit and commit-fit parameter values directly and found them significantly correlated for nearly every parameter (Fig G in S1 Appendix, S1 Appendix, “Commit likelihood and comparison with the GLM”). See S1 Appendix, “Commit likelihood and comparison with the GLM” for the full results, including Table E in S1 Appendix and the complete per-subject parameter-coefficient correlation matrix. We tested both the forgetting-based and the no-recency models against the human GLMM. Neither reproduced the over-weighting humans place on the most recent evidence strength update relative to total evidence (Fig L in S1 Appendix and Fig M in S1 Appendix).

#### Group-level correspondence with OCD symptom severity

del Río and colleagues’ central model-agnostic finding in this dataset was that the GLM’s own weight on the most recent evidence update (Δ*ES*_*t*_) decreases with symptom severity along the obsessive-compulsive spectrum [26]. Refitting the GLM to choices simulated purely from each subject’s own fitted commit-POMDP parameters, with no access to questionnaire data at any point, reproduced this relationship at a magnitude comparable to the human data, for both the OCD-symptom factor score (FA2) and the OCI-R total score (Fig. 18). This indicates that the recency-weighting-vs-OCD-symptom relationship, reported in the GLM fitted on human data, is recoverable from a mechanistic account fitted purely to individual choice behaviour; see S1 Appendix, “Commit likelihood and comparison with the GLM” for the full statistics and the corresponding model comparison against the GLM.

**Fig 18.**
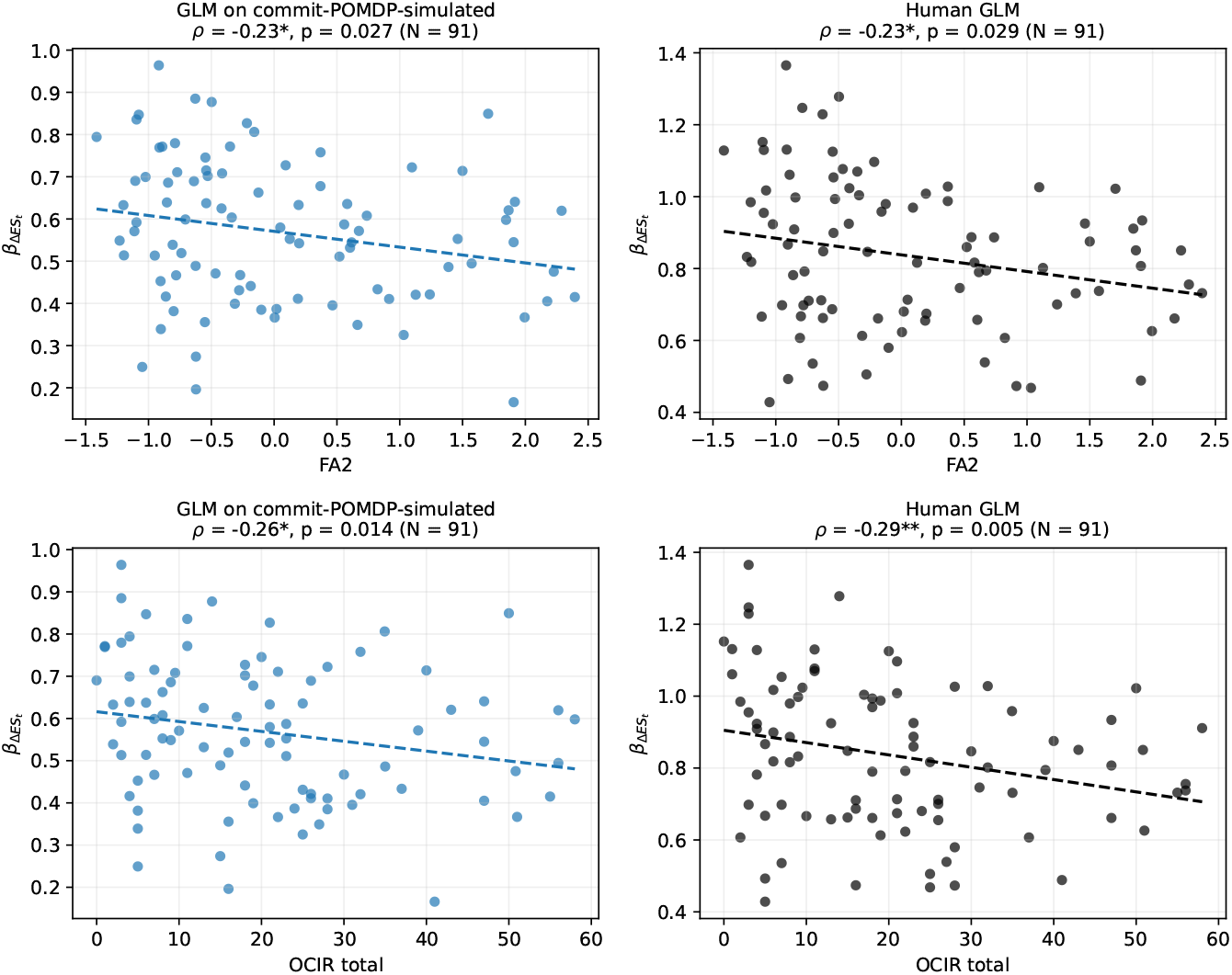
Recency-weighting regressor vs. OCD-related symptom measures. Each point is one subject. Each row is one questionnaire measure with the two fits side by side: the GLM refit to commit-POMDP-simulated choices on the left, the human GLM on the right. *Top:* vs. FA2, the obsessive-compulsive factor score. *Bottom:* vs. OCI-R total. Dashed lines are least-squares fits. Spearman *ρ* and its uncorrected *p* are annotated per panel (\**p <* 0.05, \*\**p <* 0.01).

#### Relating fitted parameters to obsessive-compulsive symptoms

We correlated every fitted parameter with every questionnaire measure held for this sample (the OC factor, OCI-R total and its six subscales, the composite compulsion score, PI-WSUR total and its five subscales, and the Y-OCS scales for the OCD patients), across all three model fits. The three scans comprise 190, 266 and 318 tests for the short, long and combined fits, of which 2, 4 and 7 reached *p <* 0.05 uncorrected, against the 9.5, 13.3 and 15.9 expected under the null. None survives Benjamini– Hochberg correction at *q* = 0.05, and every scan therefore returns fewer nominally significant associations than chance alone would produce. We accordingly treat the scan as null and do not interpret the individual nominally significant correlations; the full procedure is reported in S1 Appendix, “Parameter–symptom associations across horizon conditions”.

## Discussion

Participants were notably suboptimal in information gathering in a simple task. We therefore built a range of potential suboptimalities into a mechanistic model of the task, and examined which might be more responsible and whether they correlate with obsessive-compulsive symptoms. Two mechanisms are general: time-varying regulation of the value of waiting and an inflated subjective cost of error. Other mechanisms are more selective to the horizon of the task and individual differences. Thus, a recency bias is required by a majority of participants in the long, but not short, horizon, and equally favouring transient exaggeration and progressive forgetting. The two horizon conditions are described better by separate models than by one, with choice stochasticity and subjective cost, but not patience and belief bias, being stable within a participant across conditions.

One impetus for the original study was the fact that excessive information gathering, indexed by draws to a decision, is a standard behavioural marker in obsessive-compulsive samples [25, 26], although the literature is not unanimous [29, 34]. In our sample it is absent: the OCD patients take no more draws than the rest of the participants in either horizon condition, the 20 most compulsive take no more than the 20 least, and the number of draws is unrelated to symptom severity dimensionally. Scored against the optimal policy on the same card sequences, our subjects sample too little in the long horizon and take almost exactly the right number of draws in the short one, where they nonetheless lose accuracy by mostly sampling too much where the answer was already settled and too little where it was not.

Despite the overall similarity among the populations of participants, there could be latent differences that have gone undetected. Indeed, del Río and colleagues noted that information gathering in OCD, including their own, has concentrated on altered decision criteria such as differences in an urgency signal rather than differences in evidence integration [3, 25, 26]. This can also be motivated by the observation that recency biases are optimal when the underlying signal changes over time [18, 35, 36] – albeit a condition that does not actually apply here. The model-agnostic analyses of del Río et al. duly showed that integration does differ between healthy subjects and OCD patients.

In our model, we can compare the relative importance of urgency and recency. Temporal regulation [14], which encompasses the urgency hypothesis is indispensable in every horizon condition and its removal is by far the most damaging omission, for the goodness-of-fit, we tested, whereas recency misweighting is required only where the horizon is long enough for it to have an influence. One possible interpretation is that longer sequences make precise integration more demanding, so that participants either let old evidence decay or attend more strongly to the current draw.

The form of the recency bias is not uniform. Allowing each subject their own best-fitting structure divides the long horizon condition almost evenly between transient exaggeration, progressive forgetting, and no recency mechanism at all. A random-effects comparison over those fits returns protected exceedance probabilities close to chance for all three families. Nevertheless, the exaggeration-based model is the best single description of the pooled BIC scores for the fixed model structure across subjects.

The heterogeneity among the participants extends across symptom severity. Thus, although the relationship between recency weighting and symptom severity reported by del Río et al. for the behavioural data is recovered when the same regression is refit to choices simulated from our model, none of the factors in the model covaries with diagnosis or symptom severity at the level of the individual. This could also be a result of the limited sample: we have only 29 clinically diagnosed OCD participants, and a single session per participant. A larger clinical sample, with more trials across multiple sessions, would estimate each subject’s parameters more precisely and would be better placed to detect associations of this size.

Although fitting the horizon conditions separately within participants outperforms a combined fit, we do find some consistency. Subjective risk sensitivity *R*_risk_ and policy temperature *τ* are correlated across horizons, so a participant who treats an error as costly, or who chooses noisily, does so in both conditions. Patience and belief bias are not, and the long-horizon condition additionally reveals confusion in horizon perception, captured by the hazard lapse parameter *L*. We suggest that the individuals reparameterise their stopping rule when the horizon changes rather than employing a radically different computational structure.

The same parameter space that fails to separate individuals within one symptom dimension may still separate opposing ones. Excessive sampling has a mirror image in jumping to conclusions, the tendency to commit on very little evidence that characterises delusional thinking [4, 5, 31]. Whether the two arise from one mechanism operating in opposite directions or from distinct mechanisms is unsettled, and has been raised as an open question both for these data and for obsessive-compulsive disorder more broadly [26, 34]. A parameterised normative model makes the question concrete, because both would live in the same space: prolonged sampling as a raised subjective cost of error with weak temporal regulation, and premature commitment as the reverse. Fitting the same model to a sample spanning both symptom dimensions would say whether a single axis separates them.

We built a POMDP-based model of information gathering and added heuristic costs and mechanisms. A natural alternative would have been to have started from the classic stopping rule in the information-gathering paradigm, namely the drift-diffusion model (DDM) [9, 10, 37]. For a restricted problem with two choices, statistically-stationary evidence and no deadline, the sequential probability ratio test is the optimal stopping rule [2] as in the DDM [1, 8, 11]. Our task relaxes various of these assumptions: sequences terminate under a stochastic deadline, and the payoffs for a correct choice, an error and a missed deadline are explicit. Under a stochastic deadline, the optimal boundary is no longer fixed but collapses over time, in a way it has no closed form in general [14, 15, 38]. Rather than fit a collapsing bound to the choice-timing distributions [39, 40] we computed them using dynamic programming rather than fitted [7, 13, 41]. One benefit is that this explicitly identifies values for parameters such as subjective risk penalties (*R*_risk_) that are only implicit in the shape of collapsing bounds.

Several limitations should be noted. Trials from the two horizon conditions were intermixed throughout the session, making trial-to-trial carryover plausible, and we did not investigate or model these sequential dependencies. We evaluated transient exaggeration and progressive forgetting as mutually exclusive candidates and did not fit both within the same model, which precludes assessing whether individual participants rely on a combination of the two; given that the two are indistinguishable in the long horizon, a model allowing both is a natural next step. A further limitation is computational: the forgetting models were solved on a discretised belief grid queried by interpolation, whereas the other models were solved exactly, so a finer grid could improve their fits. The Y-BOCS was administered exclusively to the 29 participants in the OCD group, which is a relatively small clinical sample for detailed subscale analyses. Finally, our account rests on a single session per participant, and a single snapshot makes it difficult to isolate stable traits from state-dependent fluctuation; the claim that risk sensitivity and choice stochasticity are traits while patience is state-like would be properly tested by repeating the task across sessions.

## Conclusion

To answer the questions from the introduction: first, we found a number of mechanisms that were necessary to account for departures of optimality in human choices in an information gathering task. Second, urgency was a more significant contributor than recency. Third, recency was realised in structurally different ways in different participants, albeit with a rather minor effect on behaviour and behavioural suboptimality. Finally, in keeping with the subtlety of the effects on behaviour, the parameters in the model did not identify a clinical phenotype or covary substantially with symptom severity.

## Acknowledgments

We especially thank Magdalena del Río and Tobias U. Hauser for making the data of del Río and colleagues [26] available to us, and for very helpful suggestions, clarifications, and discussions throughout this work. We are also very grateful to Kenza Kadri and Aleya Marzuki for helpful discussions.

## Supporting information

**S1 Appendix**.

### Generative probability and card sequence generation

On each trial, evidence was presented sequentially over *T* discrete time steps. At each step *t*, a pair of token counts (*k*_*t*_, *n* − *k*_*t*_) was drawn from a binomial distribution,

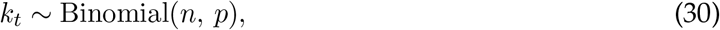

where *n* = 5 tokens were distributed at every step and *p* was the trial-specific generative probability favouring the correct option. The complementary count (*n* − *k*_*t*_) was assigned to the alternative option, so the total number of tokens per step was fixed.

The generative probability *p* was drawn independently for each trial from one of two ranges: a *high-probability* condition (*p* ∈ [0.60, 0.66], mean ≈ 0.63) and a *low-probability* condition (*p* ∈ [0.50, 0.56], mean ≈ 0.53). To ensure that the labelled correct option accumulated the majority of cumulative evidence, full sequences in which the alternative option received more total tokens were reflected (i.e. the two columns were swapped).

Sequences were generated under two horizon termination conditions. In the *short* horizon condition, trials ended after *T* ∈ {4, 5, 6, 7, 8} steps; in the *long* horizon condition, after *T* ∈ {10, 11, 12, 13, 14} steps. To equate cumulative evidence across contexts, short-context sequences were obtained by truncating the corresponding long-context sequences to the required length, rather than being sampled independently. This procedure yielded a 2 *×* 2 factorial design crossing probability level (high / low) with termination length (short / long), with 40 trials per cell. The full generation procedure is summarised in Algorithm 1.

#### Majority alignment and the truncation problem

Because short-horizon sequences are obtained by truncating their long-horizon counterparts *after* the reflection step, the guarantee that the correct option holds the cumulative majority applies only at the endpoint of the full long-horizon sequence, not at the truncation point. Formally, the reflection ensures

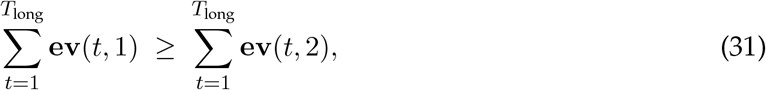

but makes no guarantee about any earlier prefix:

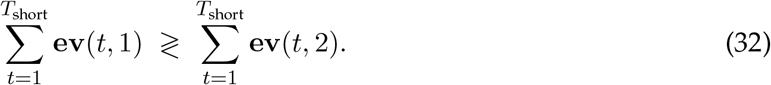

Consequently, short-horizon sequences can terminate at a point where the alternative option has accumulated more tokens than the correct option, creating a dissociation between the observed sample majority and the latent generative probability. We verified this in the sequence files: across all 160 trials in the sequences dataset, the correct option failed to hold the cumulative majority at the endpoint in 11 short-horizon trials (13.75%), whereas this never occurred in any long-horizon trial (0 of 80), consistent with the reflection guarantee in Eq. (31).

##### Algorithm 1

Trial sequence generation

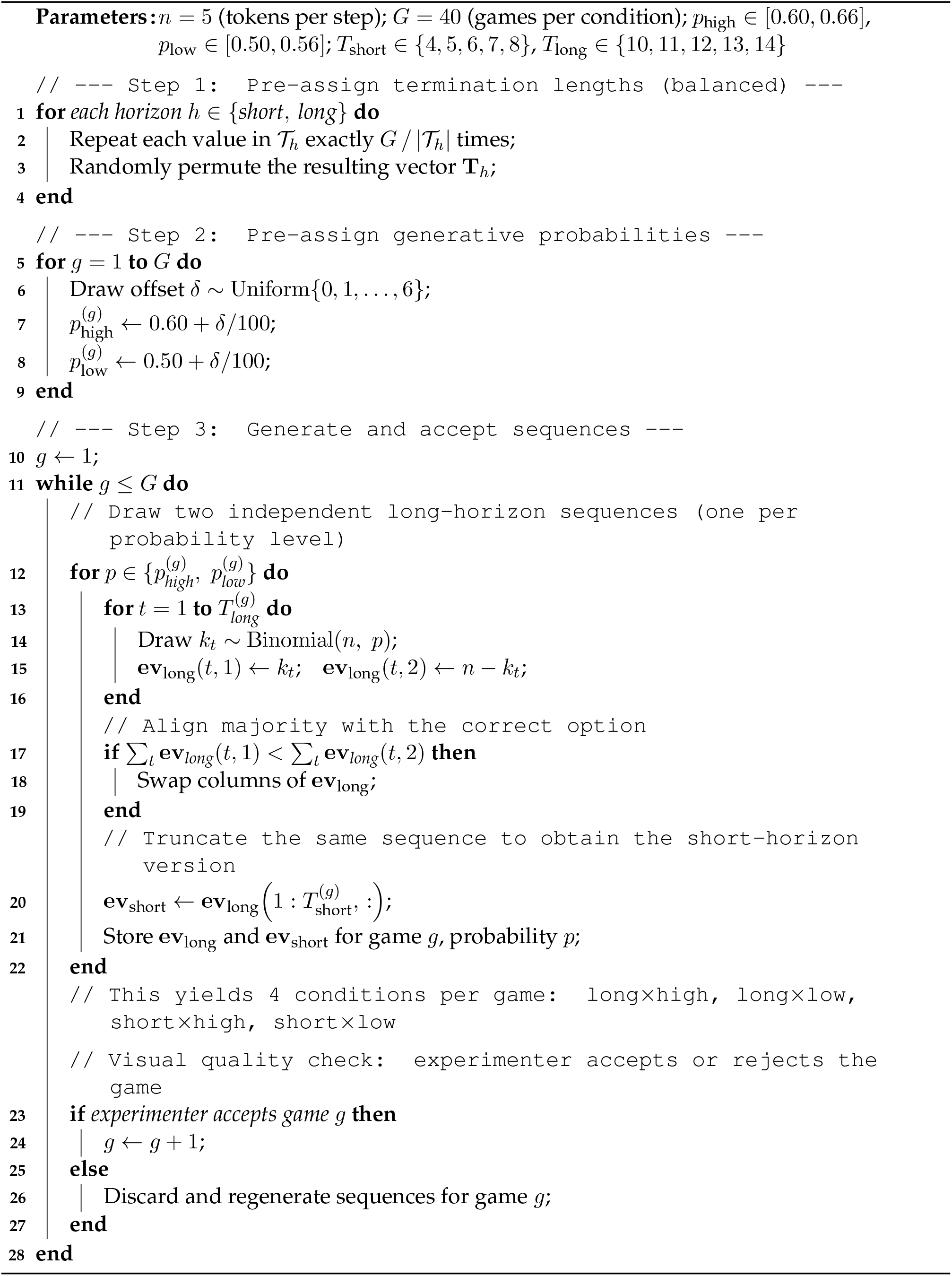

### Example *ES*_*t*_ and Δ*ES*_*t*_ calculations

Consider a trial with *N* = 5 draws, where each draw presents *n* = 5 cards. The per-draw counts of yellow cards at each draw *t* (with *t* ∈ *{*1, 2, 3, 4, 5*}*) are

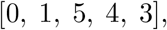

and, since five cards are shown per draw, the corresponding per-draw blue counts are

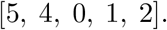

The cumulative counts 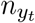 and 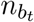 are obtained by summing the per-draw counts up to draw *t*:

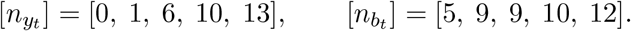

The total evidence at each draw is the absolute difference between the cumulative yellow and blue counts:

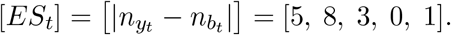

Shifting this sequence by one draw gives the total evidence at the previous draw,

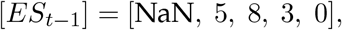

where *ES*_0_ is undefined (NaN) since no prior draw exists. Finally, the evidence strength update Δ*ES*_*t*_ is the difference between the total evidence at the current and previous draws:

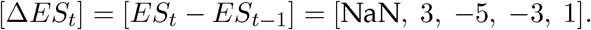

### Full candidate model set

Model specifications for the 78 candidate models compared in the Results (S1 Appendix, “Full candidate model set” and S1 Appendix, “Full candidate model set”), together with `C-EXT-R-H----`, which was fitted only to isolate a mechanism for the ablation analysis and was not entered into the model comparison (Table 4 reports only the three best-supported models, one per horizon condition). Columns as in Table 4; models are grouped by horizon condition (`S`, `L`, `C`) and generated as described in the “Candidate models’ naming” section.

**Table A.**
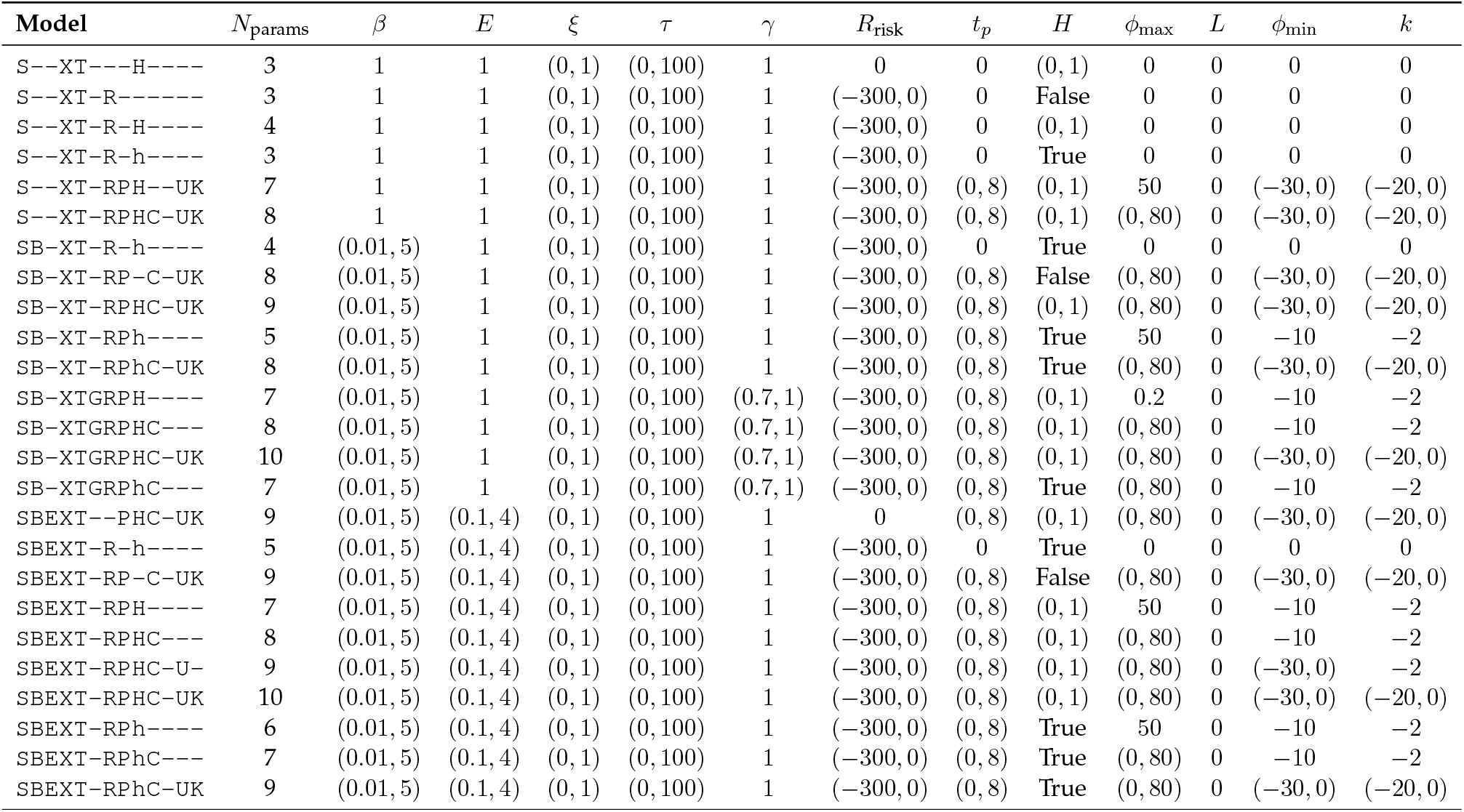
Full candidate model set (short horizon). Fitted range in parentheses; a bare value is the fixed setting for models that do not fit that parameter.

**Table B.**
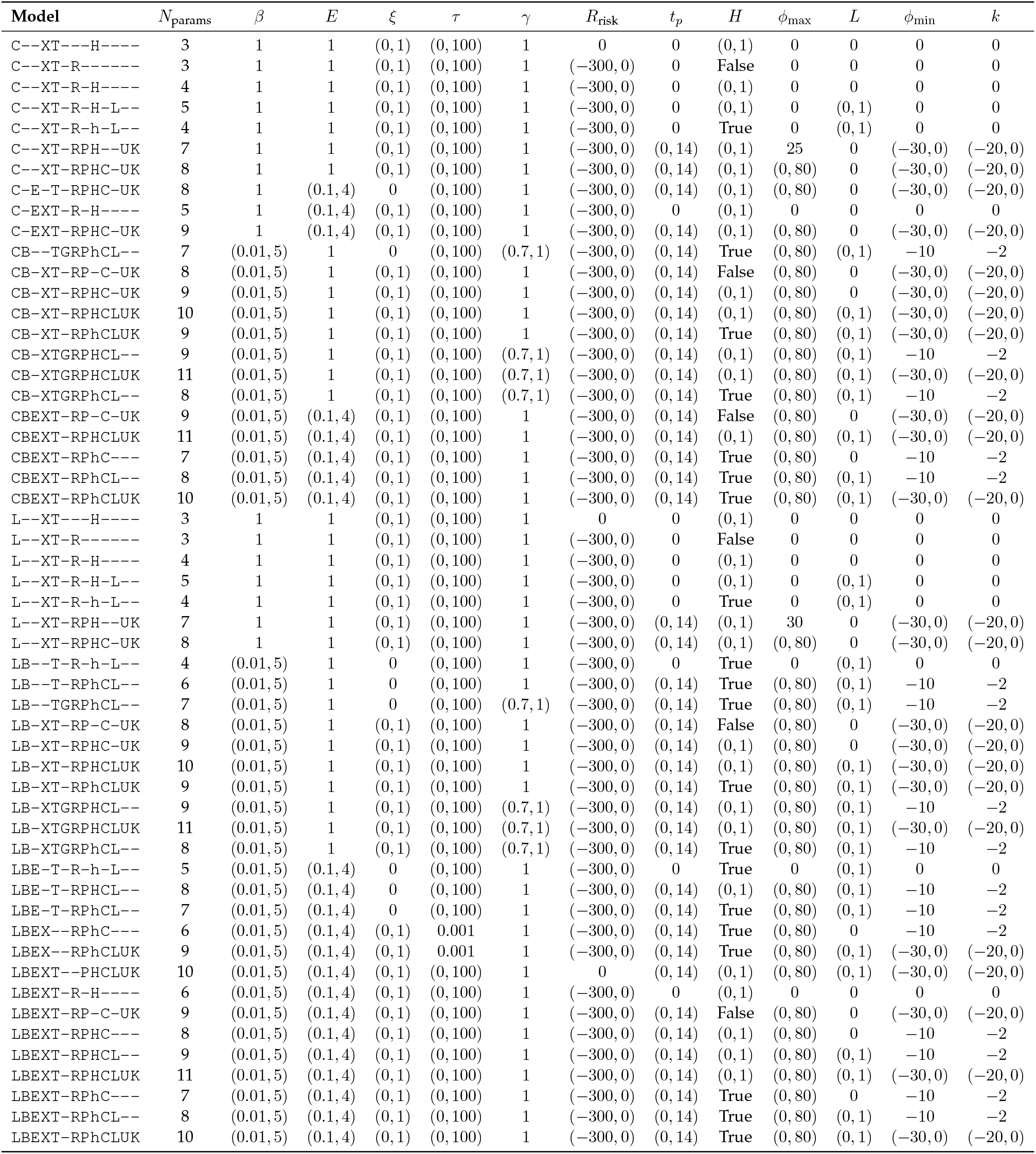
Full candidate model set (long and combined horizons). Conventions as above.

### Example individual subject fits

#### Subjects with the highest fitted exaggeration factor

Same convention as S1 Appendix, “Parameter and model recovery and sensitivity” (human, gray bars, vs. model-simulated, mean *±*SD over simulation ensembles, distributions of number of draws and outcome), shown separately, per subject, for the two subjects with the highest fitted exaggeration factor *E* under each of the two winning models that fit one (Fig A in S1 Appendix). `SB-XT-RPh----`, the best-supported short-horizon model, does not fit an exaggeration factor, so the short horizon contributes no panels. Each panel is a single subject’s own data. (a) Long-horizon subject 83 (*E* = 3.39, the highest under `LBE-T-RPhCL--`). (b) Long-horizon subject 98 (*E* = 3.37, second-highest). (c) Combined-horizon subject 17 (*E* = 2.91, the highest under `C-EXT-RPHC-UK`). (d) Combined-horizon subject 98 (*E* = 2.41, second-highest). The model continues to track these most-extreme-exaggeration individual subjects’ draw-count and outcome distributions reasonably well.

**Fig A.**
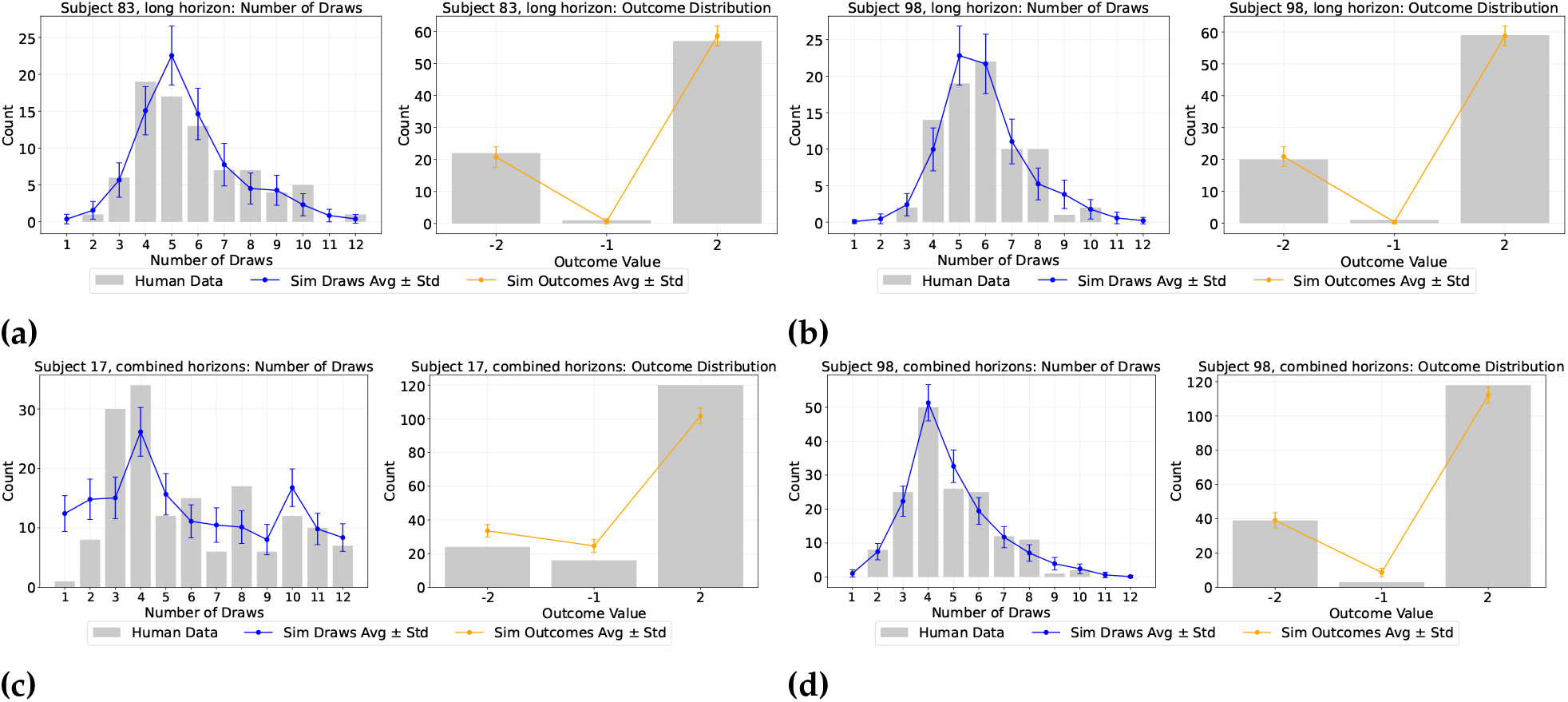
Ensemble fit for the individual subjects with the highest fitted exaggeration factor. Human (gray bars) vs. model-simulated (mean *±* SD over simulation ensembles) distributions of number of draws (left) and outcome (right). (a) Long-horizon subject 83 (*E* = 3.39, highest). (b) Long-horizon subject 98 (*E* = 3.37, second-highest). (c) Combined-horizon subject 17 (*E* = 2.91, highest). (d) Combined-horizon subject 98 (*E* = 2.41, second-highest). The short horizon is absent because `SB-XT-RPh----` does not fit an exaggeration factor.

#### Subject 17, the worst-fitting participant

The subject fitted least well by each horizon’s winning model is shown in Fig B in S1 Appendix. Each panel shows one horizon condition, fitted with that condition’s winning model. (a) `SB-XT-RPh----`, short horizon, draws-*R*^2^ = 0.94. (b) `LBE-T-RPhCL--`, long horizon, draws-*R*^2^ = 0.31. (c) `C-EXT-RPHC-UK`, combined horizons, draws-*R*^2^ = 0.34. The short-horizon distribution is well reproduced; the long-horizon one is nearly uniform across draws 2 to 14.

**Fig B.**
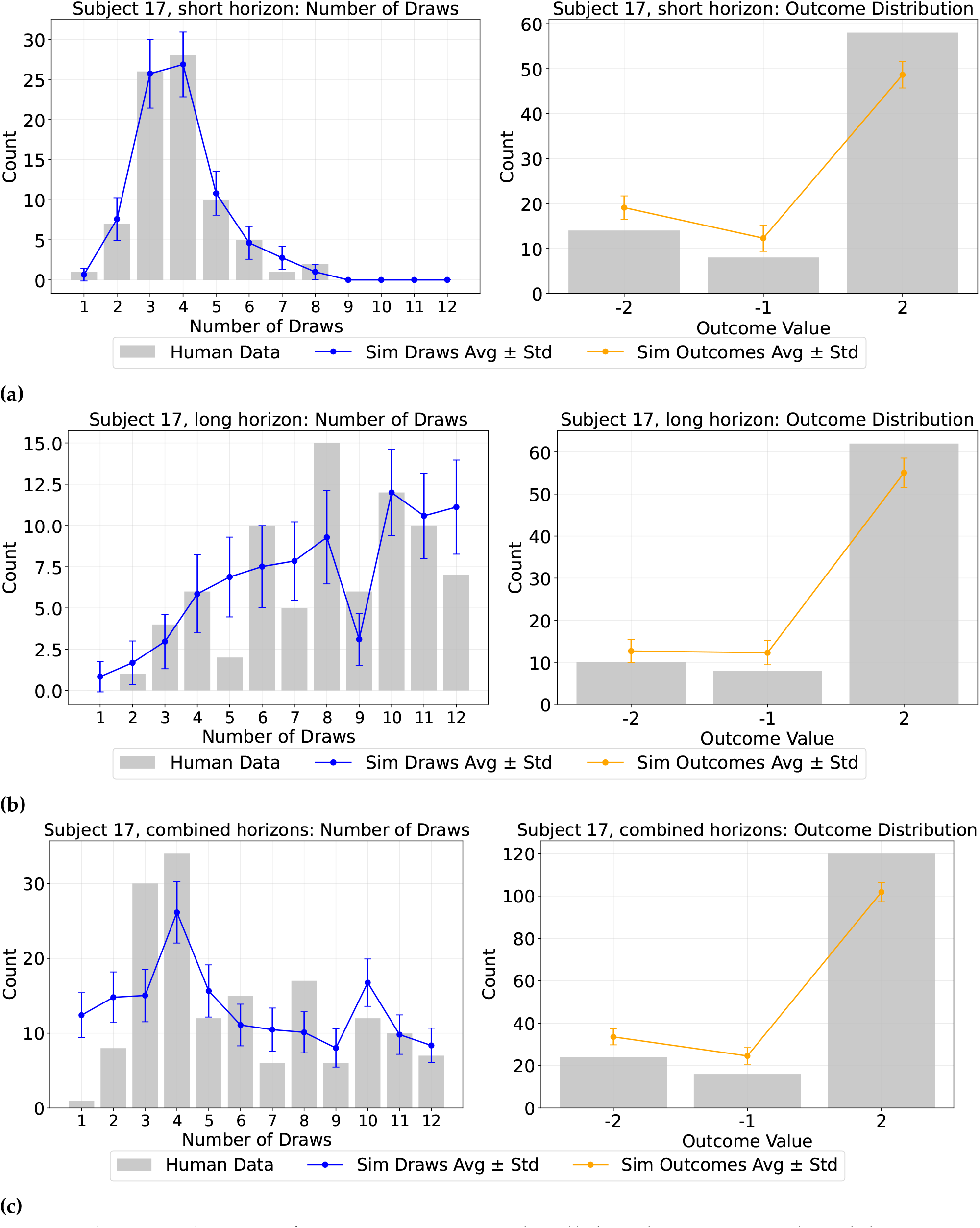
Subject 17, the worst-fitting participant, under all three best-supported models. Human (gray bars) vs. model-simulated (mean *±* SD over simulation ensembles) distributions of number of draws (left) and outcome (right), for this subject alone. (a) `SB-XT-RPh----`, short horizon. (b) `LBE-T-RPhCL--`, long horizon. (c) `C-EXT-RPHC-UK`, combined horizons.

### Parameter and model recovery and sensitivity

#### Parameter recovery

For each subject, synthetic data were simulated from that subject’s own fitted parameters (“True”) and refit blind with the same fitting procedure (“Recovered”). Each panel is one free parameter, with the identity line (perfect recovery), the fitted OLS regression line, and Pearson’s *r* (with significance) in the panel title. Fig C shows all three best-supported models: `SB-XT-RPh----` (five free parameters), `LBE-T-RPhCL--` (seven) and `C-EXT-RPHC-UK` (nine).

All five short-horizon parameters recover well (*r* = 0.72 to 0.93), the weakest being belief bias *β*. In the long-horizon parameter recovery is good for patience *t*_*p*_ (*r* = 0.96), the exaggeration factor *E* (*r* = 0.83) and the subjective cost *R*_risk_ (*r* = 0.82), and weakest for the hazard-lapse rate *L* (*r* = 0.44), which should therefore be read with caution. All nine combined-horizon parameters recover significantly, led by temperature *τ* (*r* = 0.93), the subjective cost (*r* = 0.90) and the exaggeration factor (*r* = 0.89); the two parameters of the linear temporal-regulation schedule are the weakest of the nine, with the slope *k* at *r* = 0.53 and the intercept *ϕ*_min_ at *r* = 0.62.

**Fig C.**
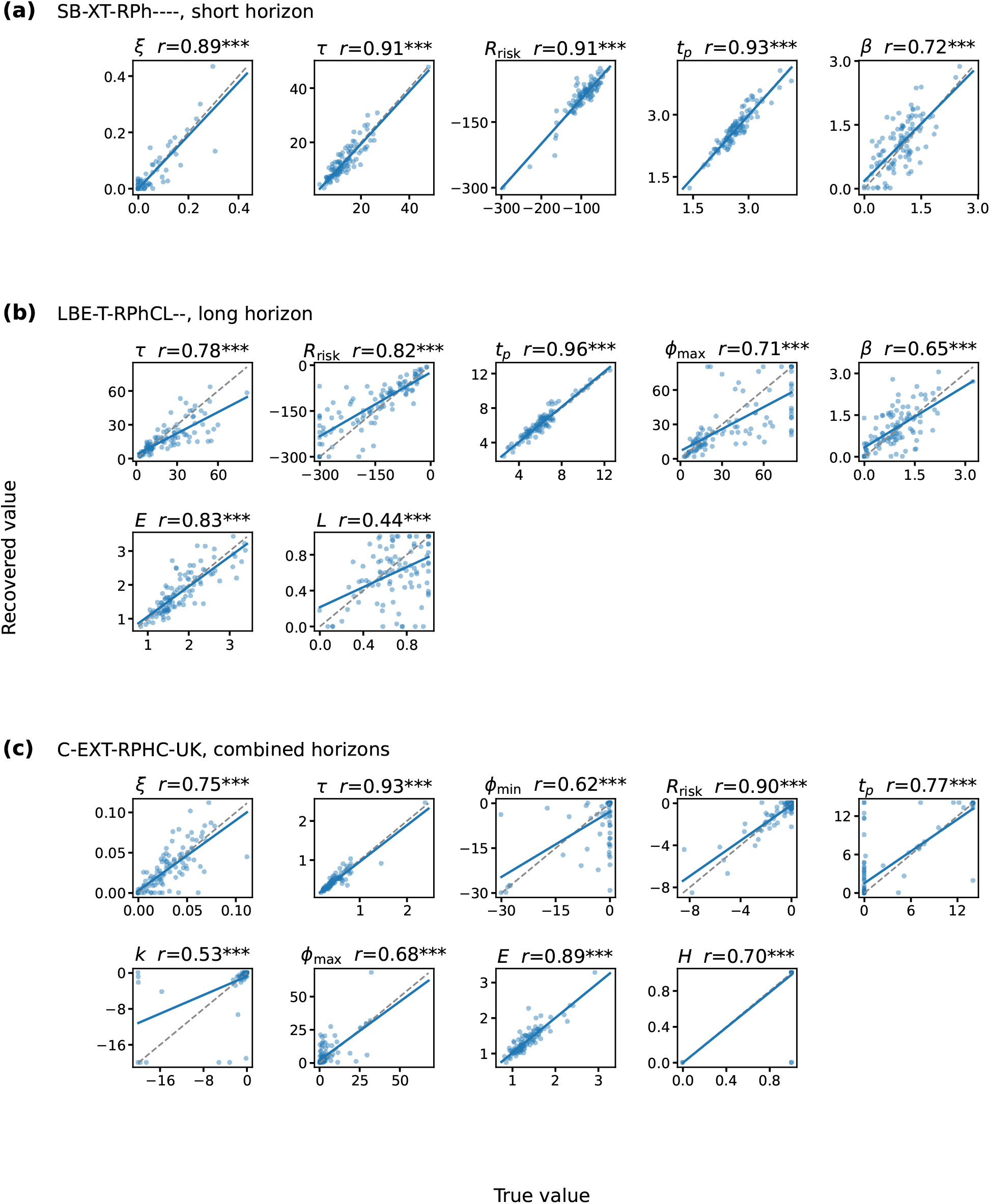
Parameter recovery for the three best-supported models. True vs. recovered values for every free parameter, from a blind refit of synthetic data simulated from each subject’s own fitted parameters (*N* = 105 per panel). Dashed line, identity; solid line, OLS fit; Pearson’s *r* with its significance in each panel title (\*\*\**p <* 0.001). **(a)** `SB-XT-RPh----`, short horizon. **(b)** `LBE-T-RPhCL--`, long horizon. **(c)** `C-EXT-RPHC-UK`, combined horizons. Axes are in each parameter’s own units.

#### Model recovery

Parameter recovery asks whether a model’s own parameters can be read back from its behaviour. Model recovery asks the separate question of whether the comparison can identify which model produced the data at all, since a model can have well-recovered parameters and still be indistinguishable from a rival. For each of the five best-supported models per horizon, we simulated data from parameters resampled from that model’s own fitted population, refit all five candidates blind to those data, and scored them with the same per-subject-summed BIC used for model selection throughout. This gives a 5 *×* 5 grid per horizon, each cell holding 105 independent subject-level fits.

Fig D reports the subject-level view: the percentage of the 105 simulated subjects whose own BIC was lowest for each candidate. Thirteen of the fifteen generating models are recovered when models are compared on the population-summed BIC, the statistic used for selection (red outlines): four of five in each of the short and long horizon conditions, and all five when the conditions are pooled. Recovery is close to complete for `SB-XT-RPh----` (100% of subjects), `LB--T-RPhCL--` (99%), `C-E-T-RPHC-UK` (95%) and `C--XT-RPHC-UK` (89%), and lower for the denser models, including `LBE-T-RPhCL--` (75%) and `C-EXT-RPHC-UK` (58%), which are nonetheless still recovered.

Both remaining failures share one structure: the winning candidate is a smaller model, and the BIC gap is carried by the parameter penalty rather than by fit quality. In the short horizon condition, `SBEXT-RPH----` loses to `SBEXT-RPh----` while fitting its own data better, by 257 BIC units of likelihood against a penalty saving of 749. In the long horizon condition, `LBE-T-RPHCL--` differs from `LBE-T-RPhCL--` only in whether the hazard indicator is fitted or fixed on the generative hazard, and its indicator sits at “on” in 99% of subjects. The two are therefore numerically the same model on these data, and the smaller one wins on a penalty saving of 789.

**Fig D.**
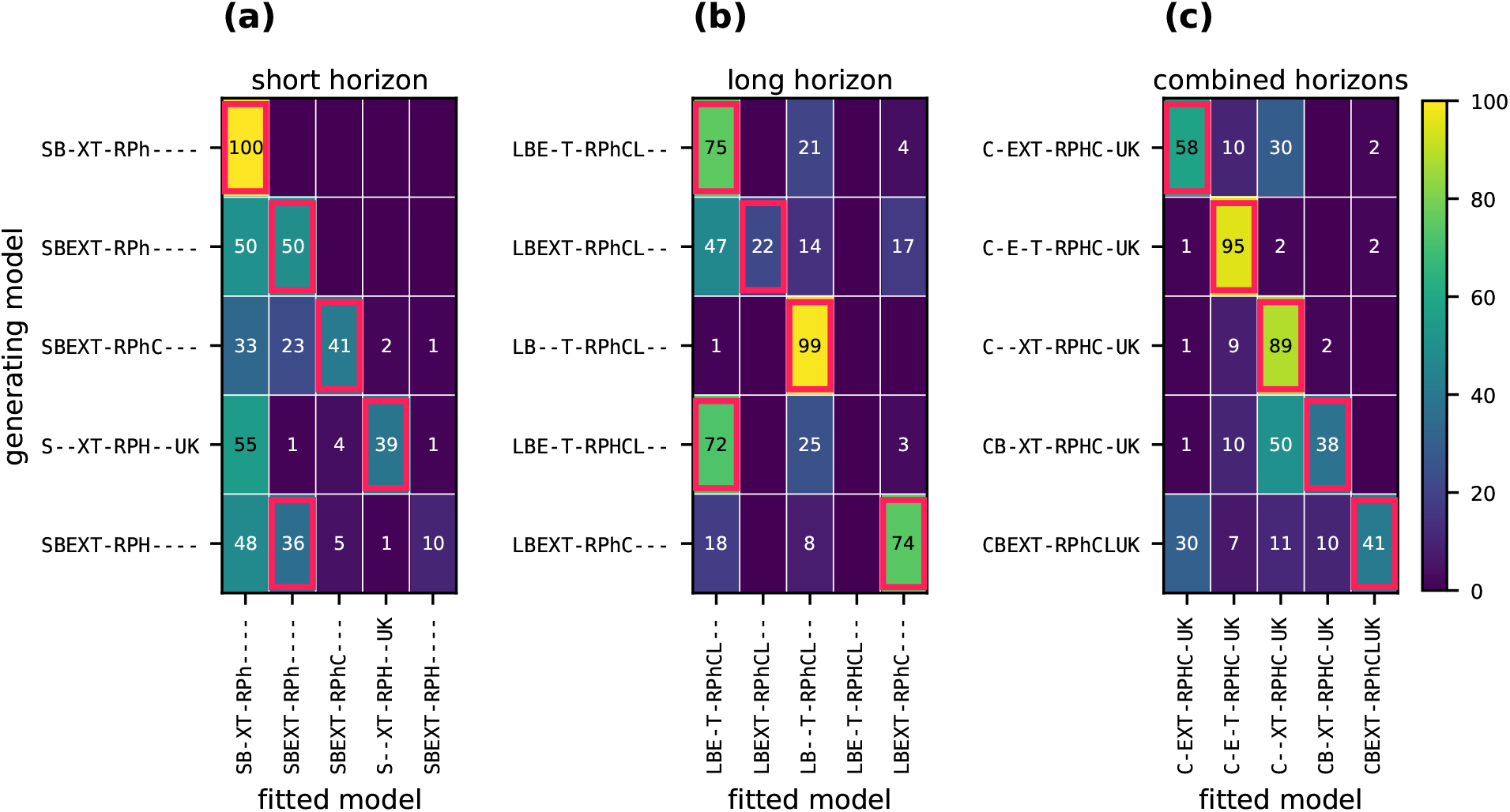
Model recovery among the five best-supported models in each horizon condition. Rows are the generating model, columns the fitted candidate, both ordered by BIC rank. Cell values give the percentage of the 105 simulated subjects whose own BIC was lowest for that candidate; empty cells are below 0.5%. Red outlines mark the candidate selected by the population-summed BIC, the statistic used for model selection. **(a)** Short horizon. **(b)** Long horizon. **(c)** Combined horizons.

#### Sensitivity of the model comparison to charging for fixed regulation parameters

The comparisons in the main text follow the standard convention of charging no parameter-count penalty for a fixed value. This convention is well justified when the fixed value is a purely structural choice (e.g. *γ* = 1 for no forgetting, *E* = 1 for no exaggeration), but is less well justified for the temporal-regulation parameters fixed to a data-informed value, as described above: some information from the sample was used to set the value, even though it was not fitted per subject.

As a sensitivity check, we recomputed BIC for every candidate charging each such parameter its exact cost. That cost is log *N*, where *N* is the pooled observation count for that horizon condition, because each fixed regulation value is a single number shared by all 105 subjects, that is, one population-level parameter estimated once, rather than a parameter each subject receives their own copy of. This comes to 10.4, 10.9 and 11.4 BIC per fixed parameter in the short, long and combined horizon conditions respectively. Candidates with the regulation function switched off entirely are not charged.

Fig E reports, for each horizon condition, the winning model, how many regulation parameters it fixes, and its margin over the closest competitor with and without this charge. The winning model is unchanged at every horizon, and so is the identity of the closest competitor. The short-horizon margin falls from ΔBIC = 93.2 to 82.8, the whole of the difference being the one extra regulation parameter `SB-XT-RPh----` fixes relative to `SB-XTGRPhC---`. The long-horizon margin is unchanged at 91.1, since `LBE-T-RPhCL--` and its closest competitor fix the same two parameters and the charge cancels between them, and the combined-horizon margin is unchanged at 315.9 because `C-EXT-RPHC-UK` fits all three regulation parameters per subject and so is not charged at all.

**Fig E.**
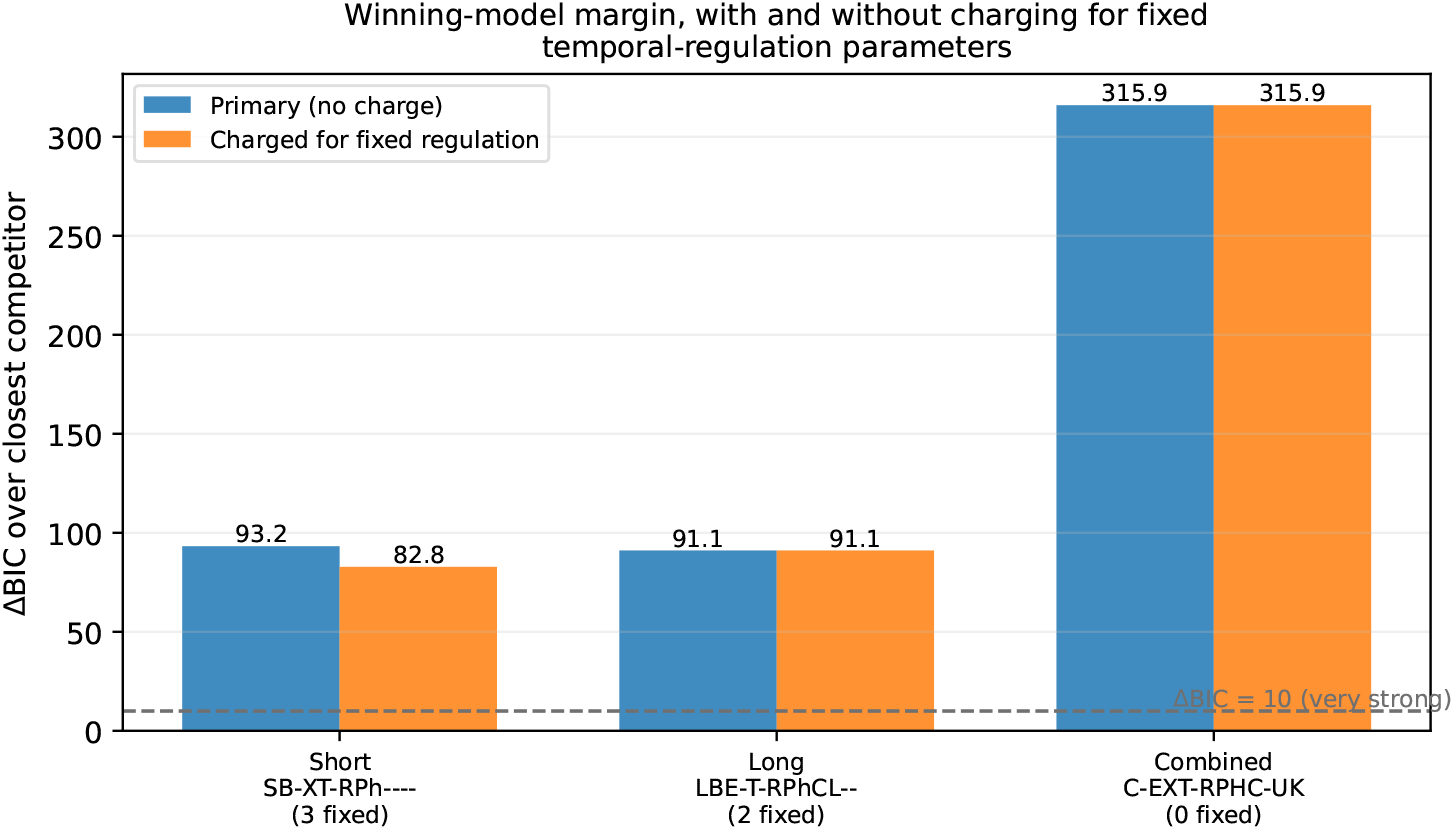
Winning-model margin with and without charging for fixed regulation parameters. ΔBIC of each horizon’s winning model over its closest competitor, under the convention used in the main text (blue) and once every candidate is charged log *N* for each regulation parameter it fixes (orange). The number of parameters each winner fixes is given under its name. The dashed line marks ΔBIC = 10, the conventional threshold for very strong evidence. Only the short-horizon margin changes, by the cost of the single extra parameter its winner fixes relative to the runner-up.

#### Sensitivity of the fixed temporal-regulation values to reoptimisation

The temporal regulation function’s shape parameters – maximum regulation *ϕ*_max_, minimum regulation *ϕ*_min_, and slope *k* – are fixed rather than fitted per subject in several best-supported models (the “Candidate models’ naming” section). Each fixed value was set by averaging that same parameter’s own fitted distribution from a closely related candidate model in which it was instead left free per subject. In this section, we check how close that choice is to the value that directly maximises log-likelihood.

For each regulation parameter that a winning model fixes rather than fits, we held every subject’s other already-fitted parameters constant and evaluated the population-summed log-likelihood at a grid of candidate values for that one parameter, holding the other regulation parameters at their originally chosen fixed values. Which parameters this covers differs by horizon condition. All three shape parameters are fixed in `SB-XT-RPh----`, so all three were swept. `LBE-T-RPhCL--` fits *ϕ*_max_ per subject, so only *ϕ*_min_ and *k* were swept. `C-EXT-RPHC-UK` fits all three per subject, so nothing about it is fixed and no sweep applies; the combined horizon is therefore absent from this analysis. *ϕ*_max_ was swept over [0, 80] and *ϕ*_min_ and *k* over [−30, 0] and [−20, 0] respectively, in each case a range at least as wide as the one used when that parameter was fitted freely in the corresponding candidate model. Because a sweep changes only the value of a parameter and not how many are free, the BIC cost of the chosen value is twice its log-likelihood deficit.

Table C and Fig F report, for each swept parameter, the chosen value, the value found by this direct search, and the BIC cost of using the chosen value instead of the found optimum. Every fixed value is at or very near its optimum. In the long horizon both swept parameters are exactly optimal (*ϕ*_min_ = −10 and *k* = −2, ΔBIC = 0.0 for each). In the short horizon the slope *k* = −2 is likewise exactly optimal, while *ϕ*_max_ = 50 against an optimum of 48 costs 4.7 BIC units and *ϕ*_min_ = −10 against an optimum of −10.5 costs 6.5. Both of those are below the conventional threshold for even positive evidence (ΔBIC = 10; Kass & Raftery) and roughly an order of magnitude below the margins separating the winning models from their closest competitors (Tables 7–10). In every case the chosen value sits within a shallow, single-peaked region of the log-likelihood surface (Fig F), so none of these choices leaves a substantial amount of fit quality on the table.

**Table C.**
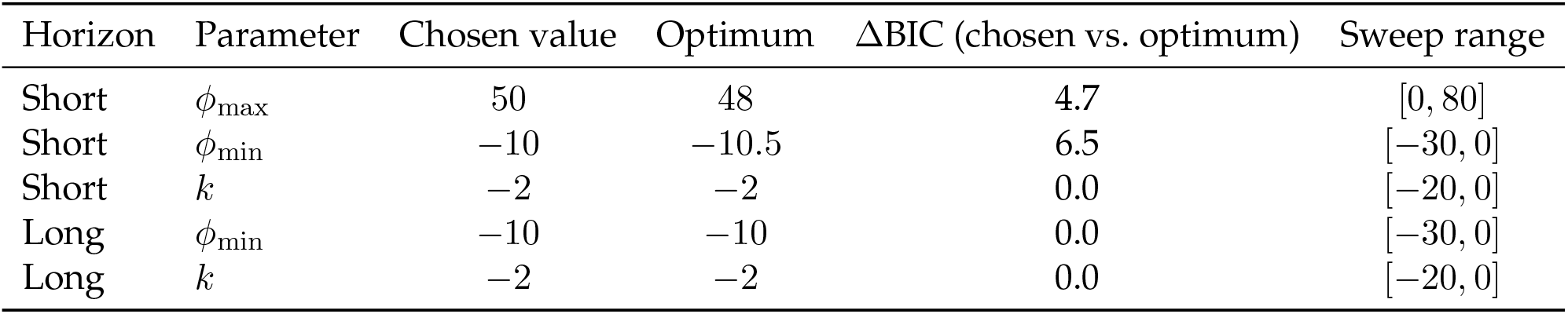
Chosen vs. directly reoptimised fixed temporal-regulation values. For each regulation parameter that `SB-XT-RPh----` or `LBE-T-RPhCL--` fixes rather than fits, the value it was fixed to, the value found by directly maximising population-summed log-likelihood over the swept range (holding every subject’s other parameters fixed), and the resulting BIC cost of the chosen value relative to that optimum, equal to twice its log-likelihood deficit. `C-EXT-RPHC-UK` fits all three regulation parameters per subject and so has nothing fixed to sweep.

**Fig F.**
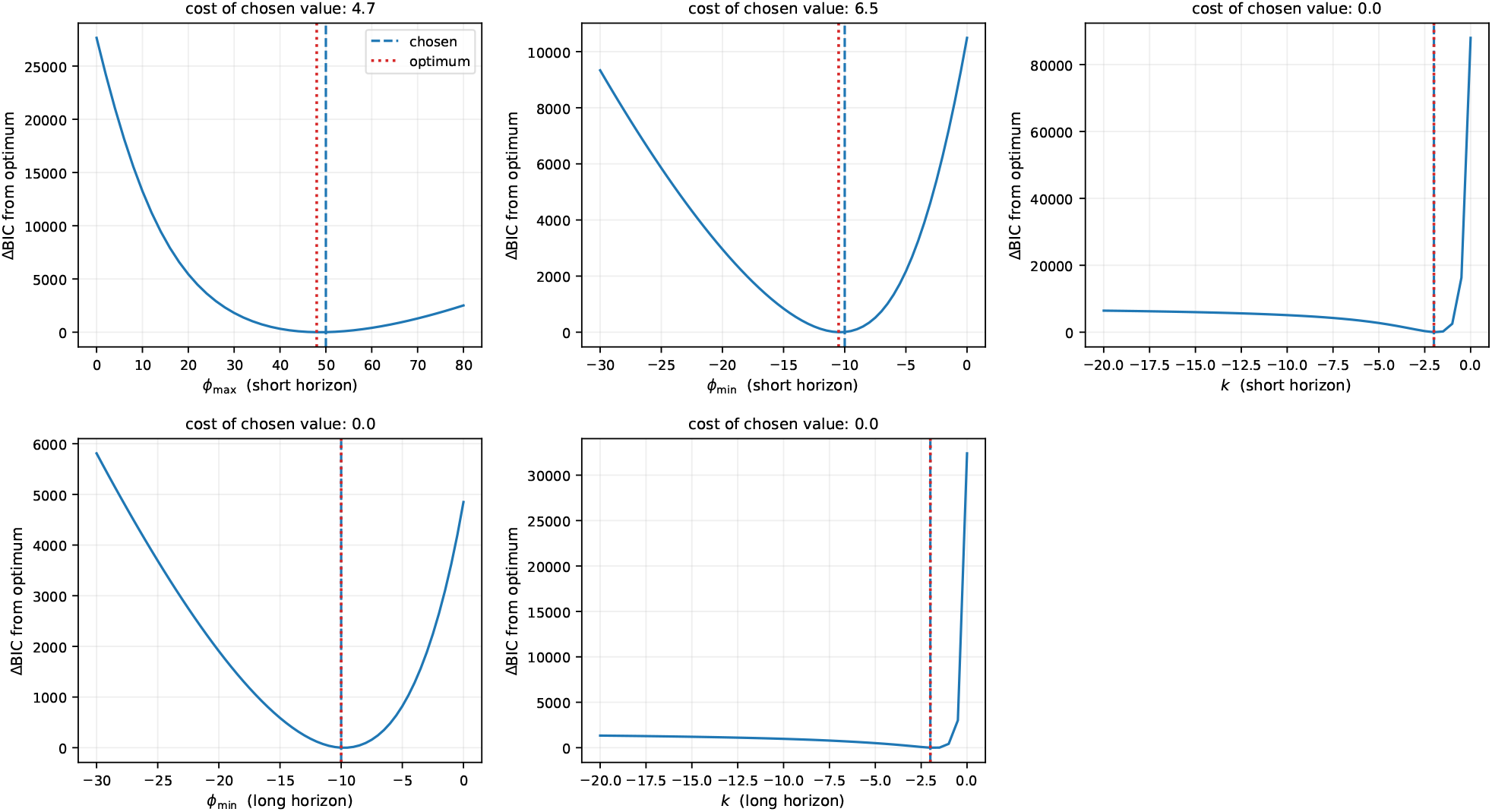
Population-summed ΔBIC as a function of each fixed temporal-regulation parameter, relative to its optimum. Each panel sweeps one parameter (holding every subject’s other parameters, including the other two regulation parameters, fixed) and plots ΔBIC relative to the minimum found over the sweep. Dashed blue line: the value the parameter was actually fixed to. Dotted red line: the value found by this search.

### Commit likelihood and comparison with the GLM

#### Commit-versus-wait likelihood

In the “Model fitting” section, we discuss fitting the models to a likelihood (Eq. 25) that solely considers when a decision is made rather than also which decision. This was to match results reported by del Río and colleagues. In this section, we describe the results using this restricted likelihood.

#### Are the full-fit and commit-fit parameter estimates the same?

To understand the relation between the full-fit vs. commit-fit POMDP models, we refit each horizon’s winning *structure* under the commit objective and compared, for every free parameter, each subject’s full-fit value against their own commit-fit value from that same-structure refit, using the Pearson correlation across subjects. Fig G reports both. In the short models, temperature *τ* and subjective cost *R*_risk_ agree closely under either objective in both horizons (*r* = 0.94 and 0.88 short; *r* = 0.92 and 0.88 long), and patience *t*_*p*_ is similarly consistent (*r* = 0.74 short, *r* = 0.95 long), as is the long-horizon ceiling *ϕ*_max_ (*r* = 0.93). Belief bias is the weakest shared parameter in both horizons (*r* = 0.45 short, *r* = 0.28 long), and also the weakest-recovering of the short-horizon parameters (*r* = 0.72, Fig Ca). The long-horizon exaggeration factor (*r* = 0.33) and hazard lapse (*r* = 0.47) agree only moderately. The short-horizon lapse rate *ξ* is the clear exception: full-fit and commit-fit values are essentially uncorrelated (*r* = 0.10, *p* = 0.31). This is justified, since the commit likelihood does not pay attention to the correct choice but only whether to continue or commit (Fig Ga).

**Fig G.**
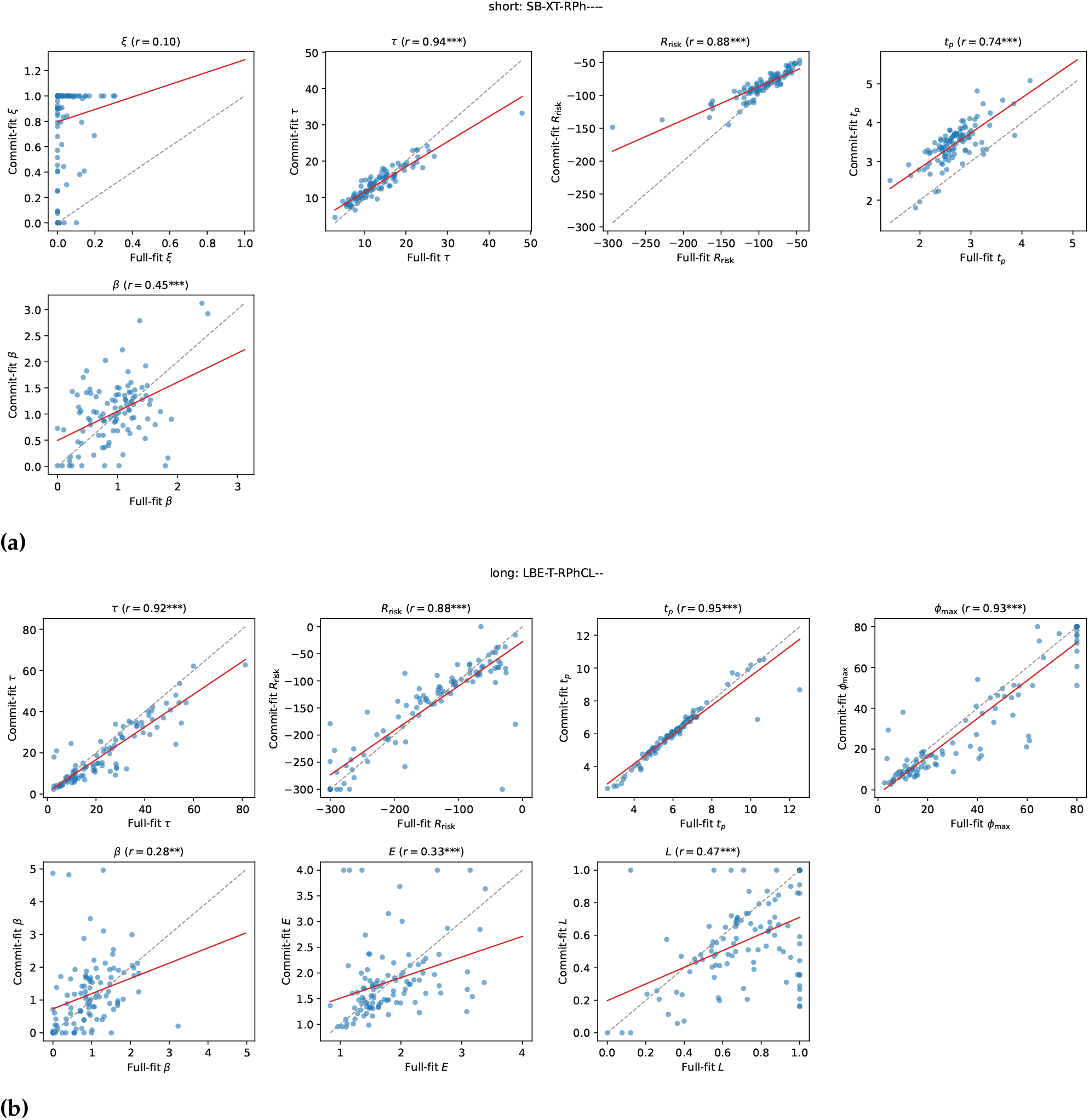
Full-fit vs. commit-fit parameter values, per subject, for the same model structure refit under each objective. Each point is one subject; the dashed line is *y* = *x* (perfect agreement); the red line is a linear regression fit with a 95% confidence band. Pearson *r* is annotated per panel. (a) Short horizon, `SB-XT-RPh----`. (b) Long horizon, `LBE-T-RPhCL--`.

#### Comparison between the POMDP-commit models and the GLM

We compare the commit-likelihood POMDP models against the GLM/GLMM (fitted on the human behavioural data [26]) using the same per-subject BIC/AIC (each subject’s own BIC/AIC computed from their own *n*_obs_ and log-likelihood, then summed across subjects).

Table D and Fig H compare the best-supported combined-horizons commit model, the best-supported separate short+long commit models, and the GLM, applying one fixed model to every subject and using the per-subject-summed AIC/BIC convention used throughout this manuscript (each subject’s own *n*_obs_; Model comparison). The separate short+long commit fit has the lowest (best) value on both criteria (AIC = 52230.5, BIC = 57739.1), ahead of the GLM (AIC = 54720.0, BIC = 58208.3) and the combined-horizons fit (AIC = 54094.6, BIC = 58579.6). The margin over the GLM is comfortable on AIC (ΔAIC = 2489.5) but narrower on BIC (ΔBIC = 469.2), reflecting the separate fit’s larger per-subject parameter count (*k* = 13, vs. the GLM’s *k* = 7). The combined-horizons fit is the only one of the three that GLM beats on BIC, by 371.3. The mechanistic account, for the separate model, is therefore preferred by both criteria in this comparison.

**Table D.** Fixed-model selection metrics: POMDP (commit, combined), POMDP (commit, separate short+long), and GLM. sum logL, *k*, and *n*_obs_ are summed/pooled across the short and long fits for the separate-model row, matching Table 10’s convention.

| Model | sum logL | $N_{\text{params}}$ | AIC | AICc | BIC |
| --- | --- | --- | --- | --- | --- |
| C-EXT-RPHC-UK | -26102.28 | 9 | 54094.56 | 54117.23 | 58579.56 |
| SBEXT-RPh---- + LBEXT-RPhC--- | -24750.24 | 13 | 52230.48 | 52282.03 | 57739.05 |
| GLM | -26624.98 | 7 | 54719.96 | 54734.03 | 58208.29 |

**Fig H.**
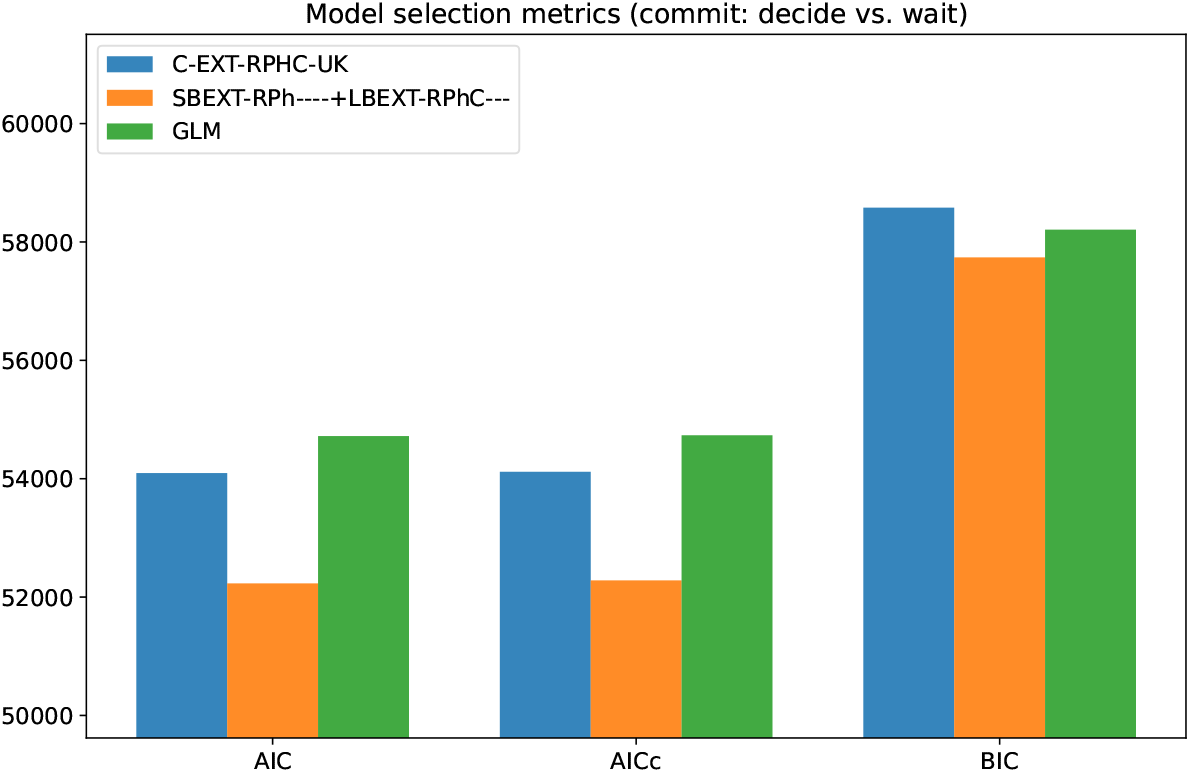
Fixed-model BIC: POMDP (commit) vs. GLM. Raw AIC, AICc, and BIC for the combined-horizons commit fit, the separate short+long commit fit, and the GLM.

#### Behavioural signature: GLMM and per-subject GLM correspondence

The comparisons above all use BIC/AIC, which score how well a model predicts the exact timing of each decision. Here we instead ask whether the best-supported commit POMDP (`SBEXT-RPh----` for short, `LBEXT-RPhC---` for long, combined into one short+long fit) reproduces the same *regression signature* the GLMM/GLM analyses describe in the human data. We refit the same GLMM to synthetic choice sequences generated by this commit-fit POMDP pair and overlaid the resulting fixed effects on the human estimates (Fig I, Table E). The model captures the same overall pattern as the human GLMM, including the same significant terms.

**Table E.** GLMM fixed-effect estimates: human data vs. commit-POMDP-simulated choices. Same convention as Fig I; “Difference” is the model estimate minus the human estimate.

| Predictor | Human | LBEXT-RPhC--- | Difference |
| --- | --- | --- | --- |
| $ES_{t-1}$ | 0.462 | 0.417 | -0.045 |
| $\Delta ES_t$ | 0.795 | 0.541 | -0.254 |
| trial | 1.895 | 1.938 | 0.044 |
| termination | 1.523 | 1.544 | 0.021 |
| $FA2 \times ES_{t-1}$ | -0.007 | -0.002 | 0.005 |
| $FA2 \times \Delta ES_t$ | -0.042 | -0.034 | 0.007 |
| $FA2 \times \text{trial}$ | -0.042 | -0.011 | 0.030 |
| $FA2 \times \text{termination}$ | -0.042 | -0.020 | 0.022 |

We additionally fit a (non-mixed-effects) GLM separately to each subject’s human choices and to that same subject’s choices simulated from their own best-fitting commit-POMDP parameters (averaged over the simulation ensemble), then correlated the two sets of subject-level coefficients across all six regressors plus the intercept (Fig J). All seven coefficients showed strong positive correlations (*r* = 0.67–0.97), with the strongest correspondence for the model’s intercept (*r* = 0.97) and the weakest for *ES*_*t*−1_ (*r* = 0.67).

Correlating each subject’s fitted commit-POMDP parameters with the GLM regressors fitted on that subject’s simulated data (Fig K) shows that the exaggeration factor correlates significantly and negatively with the total-evidence regressor (*ES*_*t*−1_; *r* = − 0.48, *p <* 0.001), consistent with it acting as a mechanism that shifts the relative weighting of the last piece of evidence against the total accumulated evidence. The patience parameter shows the strongest overall associations, dominating the intercept (*r* = −0.62, *p <* 0.001) and termination (*r* = 0.39, *p <* 0.001) weights.

del Río and colleagues’ central model-agnostic finding was that the GLM weight on the most recent evidence update (Δ*ES*_*t*_) decreases with symptom severity along the obsessive-compulsive spectrum [26], quantified using the obsessive-compulsive factor score (FA2) defined in Model fitting above. We asked whether this same relationship is reproduced when the GLM is instead fit to choices simulated from the commit-fit POMDP rather than to the real human choices, using each subject’s own FA2 score and OCI-R total score (OCIR total). In the human data, 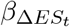 correlated negatively with both FA2 (*ρ* = −0.23, *p* = 0.029) and OCIR total (*ρ* = −0.29, *p* = 0.005); the GLM refit to commit-POMDP-simulated choices reproduced both relationships at comparable magnitude (FA2: *ρ* = −0.23, *p* = 0.027; OCIR total: *ρ* = −0.26, *p* = 0.014; Table F, Fig 18), even though no questionnaire information was used when fitting the POMDP to each subject’s own choices. This indicates that the recency-weighting-vs-OCD-symptom relationship reported in the human data is not an artifact specific to the GLM’s own fitting procedure, but is recoverable from a mechanistic account fitted purely to individual choice behaviour.

**Fig I.**
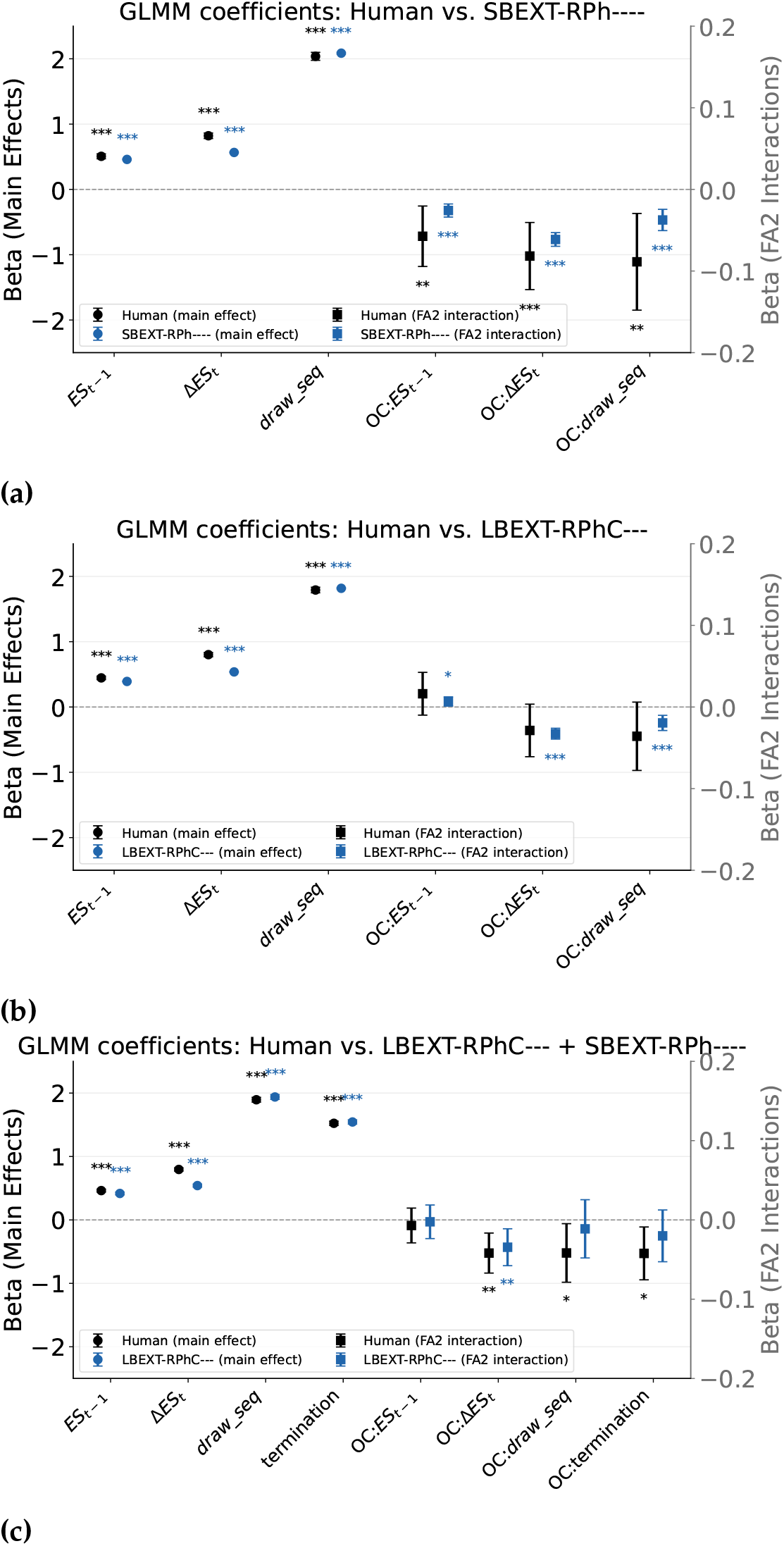
GLMM coefficients: human data vs. commit-POMDP-simulated choices. Fixed-effect estimates (*±*95% CI) refit on simulated choices (blue) and overlaid on the human estimates (black); main effects on the left axis (circles), FA2 interactions on the right axis (squares); asterisks give each estimate’s own significance. **(a)** Short horizon, against `SBEXT-RPh----`. **(b)** Long horizon, against `LBEXT-RPhC---`. **(c)** Both horizons combined into one fit, which adds the termination regressor; termination is constant within a single horizon and so is dropped in (a) and (b). Model intervals are narrower because they come from the pooled simulated ensemble.

**Fig J.**
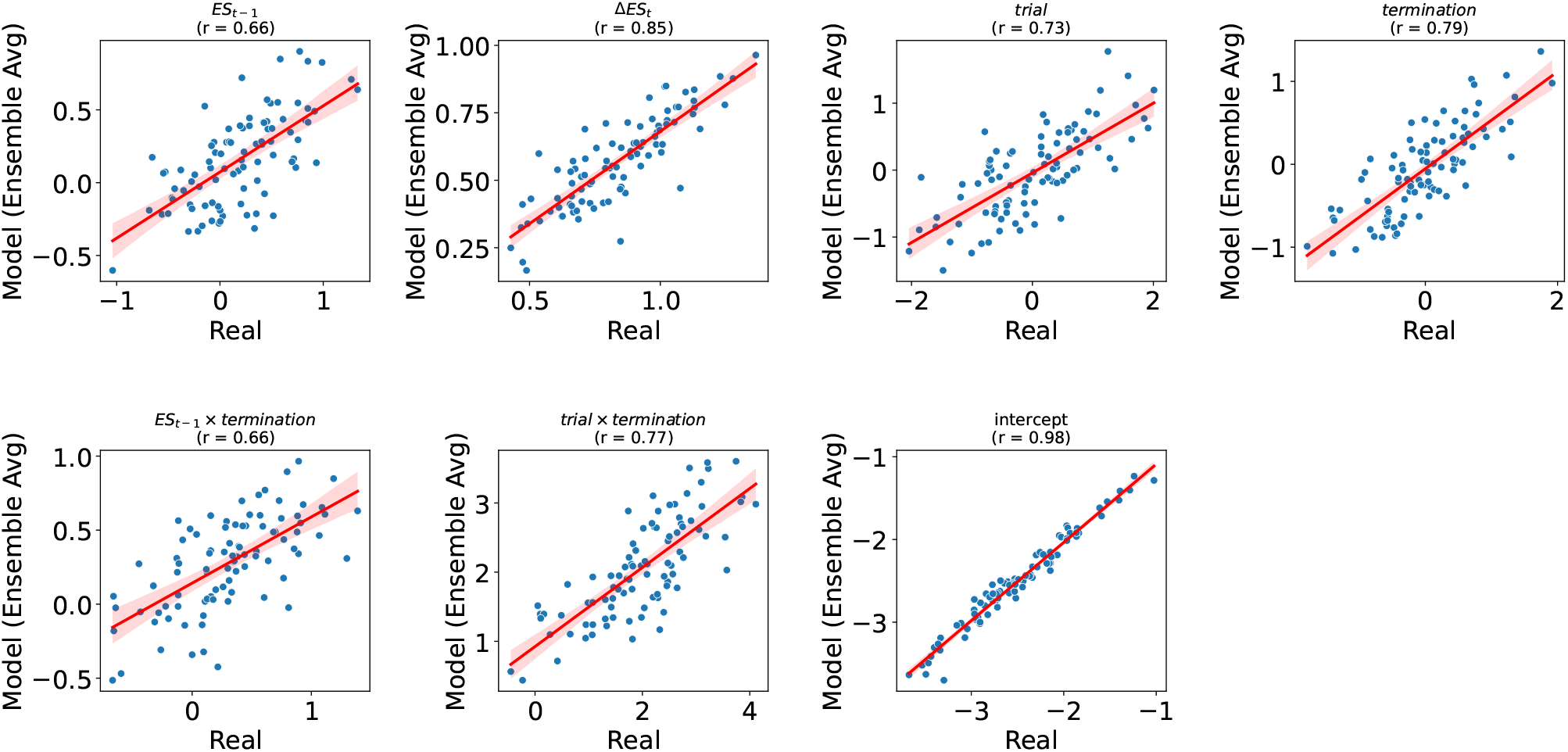
Per-subject correspondence between human and commit-POMDP-simulated GLM coefficients. Each point is one subject’s GLM coefficient fit to real human choices (*x*-axis) vs. the same subject’s coefficient fit to choices simulated from their own best-fitting commit-POMDP parameters (*y*-axis). Pearson’s *r* for each regressor is given in the panel title.

**Fig K.**
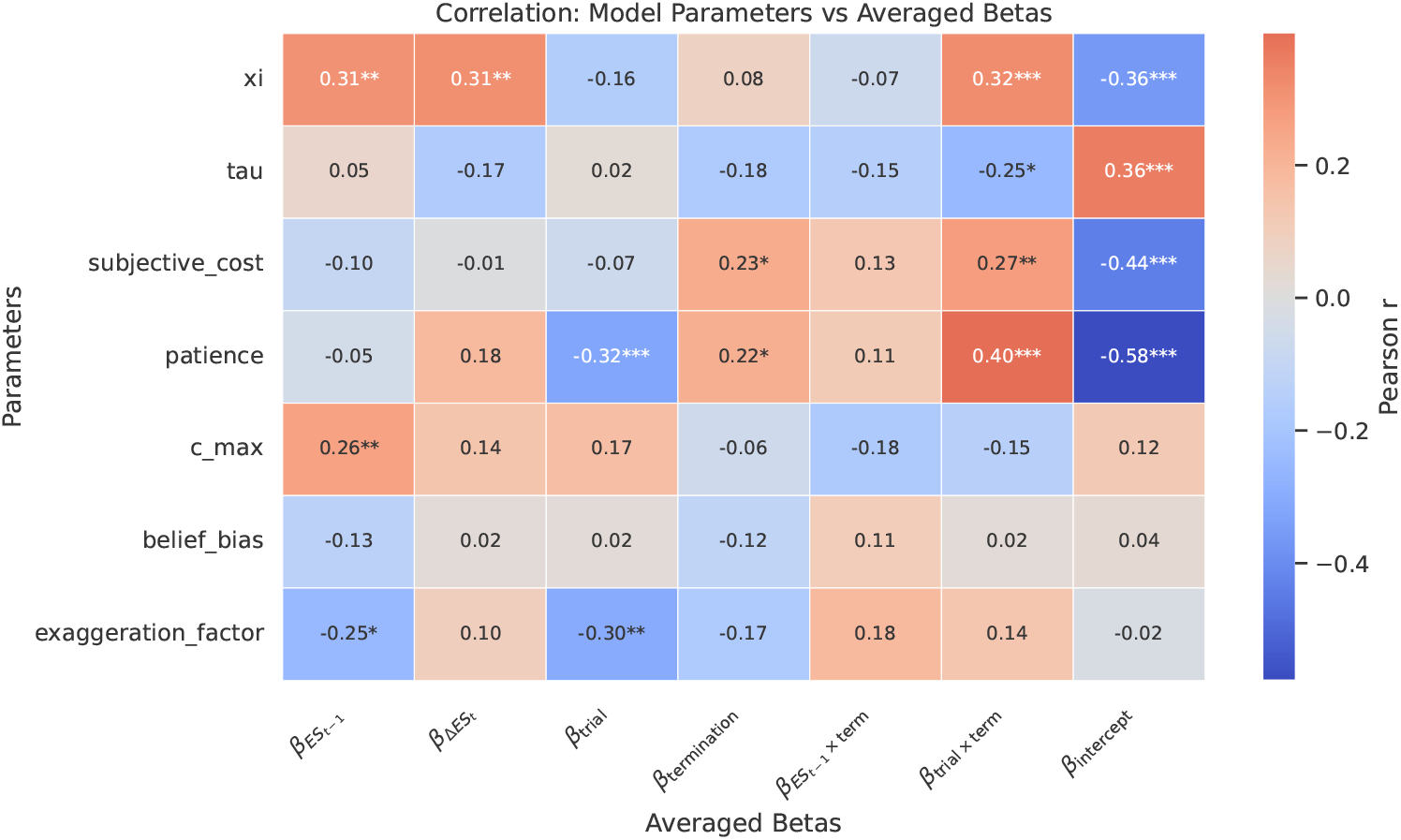
Correlation between commit-POMDP parameters and simulated GLM coefficients. Pearson correlation (annotated; \**p <* 0.05, \*\**p <* 0.01, \*\*\**p <* 0.001) between each subject’s fitted commit-POMDP parameters (rows) and that subject’s ensemble-averaged, model-simulated GLM coefficients (columns).

**Fig L.**
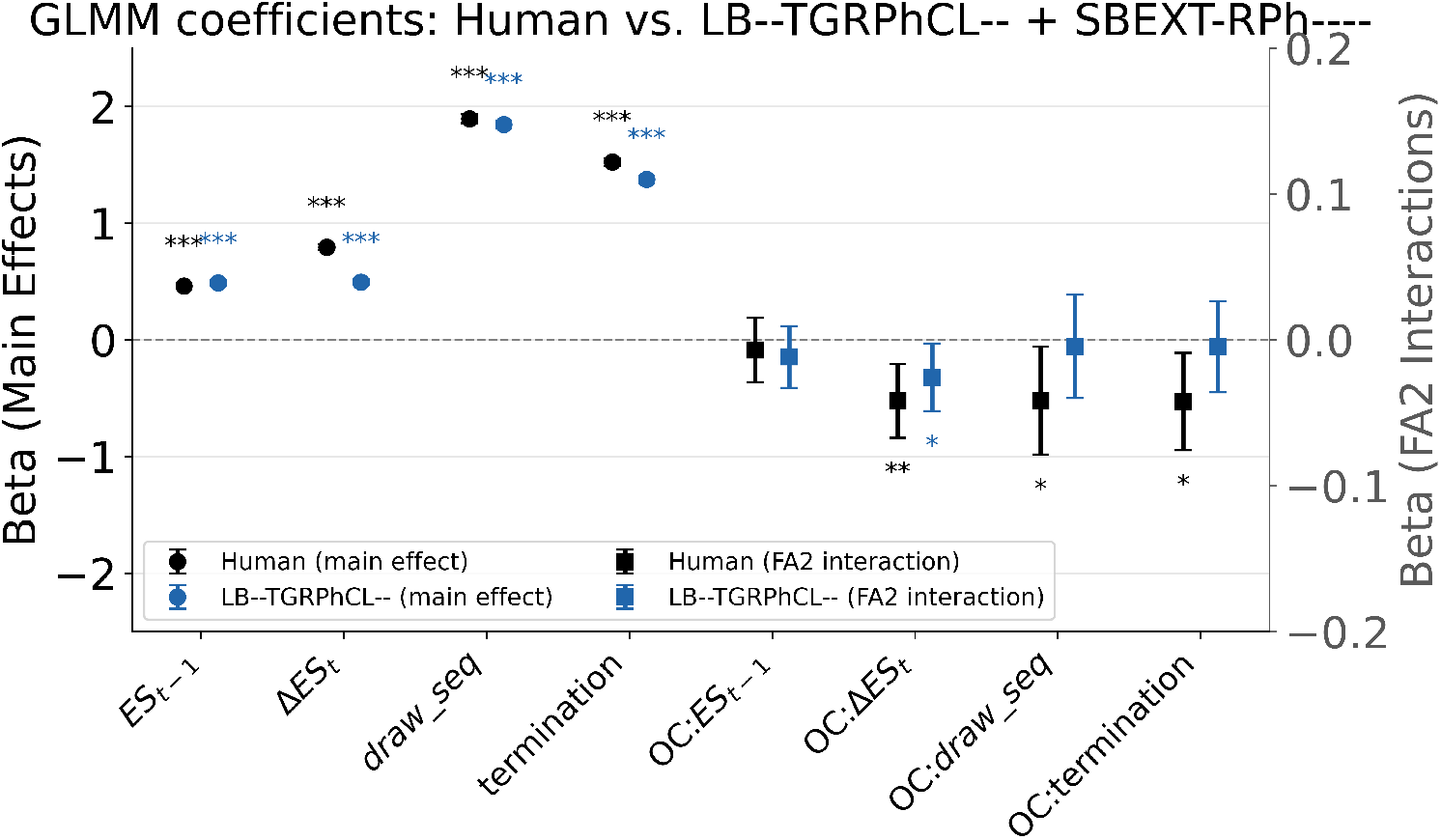
GLMM coefficients: human data vs. the best forgetting-based commit-POMDP models. Fixed-effect estimates (*±*95% CI) refit on choices simulated by `LB--TGRPhCL—`with `SBEXT-RPh----` (blue) and overlaid on the human estimates (black); main effects on the left axis (circles), FA2 interactions on the right axis (squares); asterisks give each estimate’s own significance. Both horizons are fitted together, as in Fig Ic. Forgetting matches the weight on total evidence *ES*_*t*−1_ but recovers only 0.49 of the human 0.80 on the most recent update Δ*ES*_*t*_.

**Fig M.**
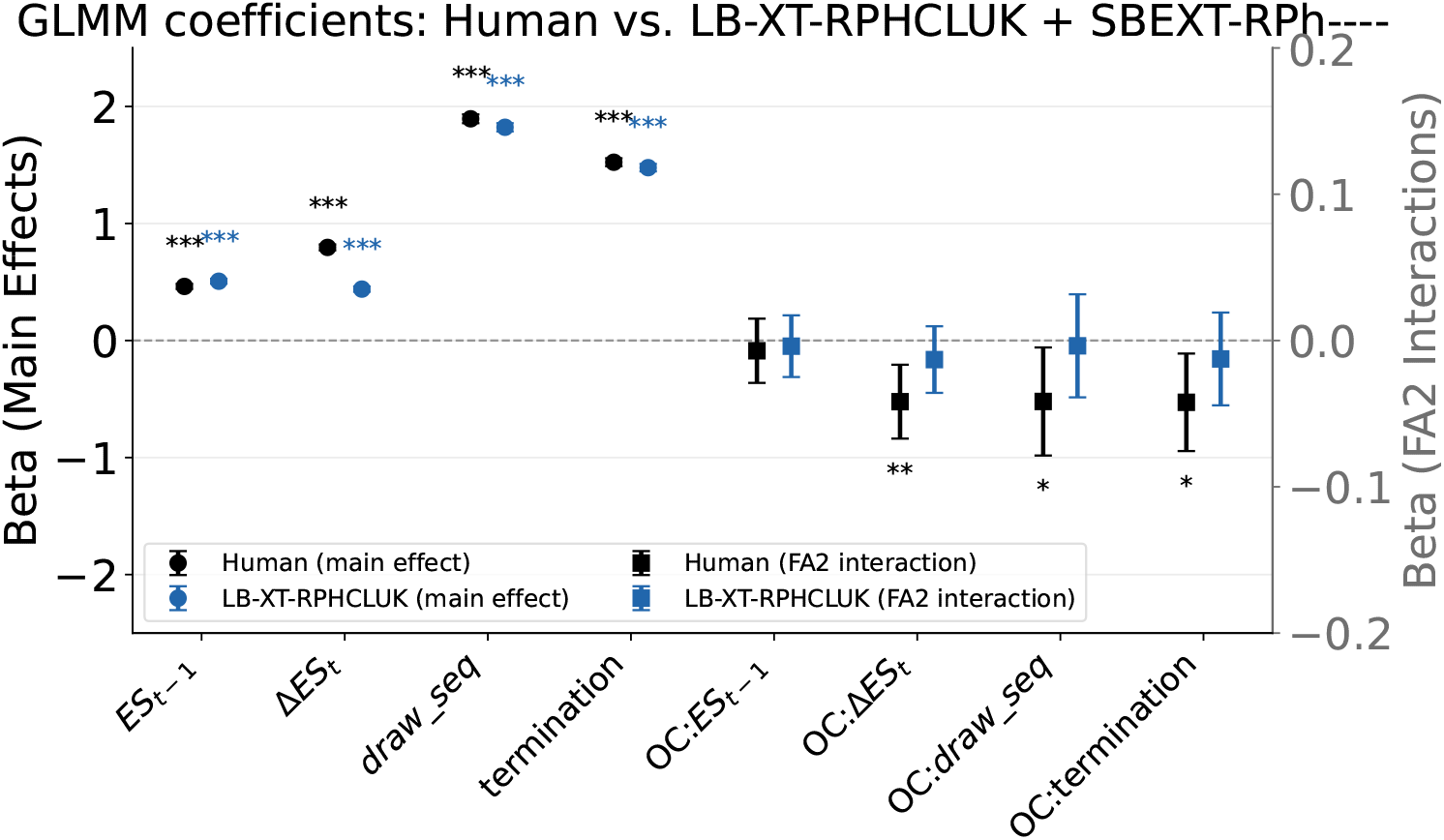
GLMM coefficients: human data vs. a commit-POMDP model with no recency mechanism. As Fig L, for `LB-XT-RPHCLUK` with `SBEXT-RPh----` (blue) against the human estimates (black). Removing recency leaves *ES*_*t*−1_ and the time-pressure terms near their human values but drops Δ*ES*_*t*_ further, to 0.44 against 0.80, and none of the three FA2 interactions significant in humans reaches significance here.

**Table F.** Recency-weighting regressor 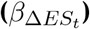 vs. OCD-related symptom measures: human GLM vs. GLM on commit-POMDP-simulated choices. Spearman *ρ* between each subject’s Δ*ES*_*t*_ coefficient and their OCD-symptom factor score (FA2) or OCI-R total score (OCIR total; \**p <* 0.05, \*\**p <* 0.01).

| Source | Questionnaire measure | Spearman $\rho$ | $p$ |
| --- | --- | --- | --- |
| GLM on commit-POMDP-simulated | FA2 | −0.232* | 0.027 |
| Human GLM | FA2 | −0.229* | 0.029 |
| GLM on commit-POMDP-simulated | OCIR total | −0.256* | 0.014 |
| Human GLM | OCIR total | −0.292 * * | 0.005 |

#### Per-horizon GLMM on a single simulated dataset

The GLMM comparisons in S1 Appendix, “Commit likelihood and comparison with the GLM” is repeated against a single randomly drawn simulation per horizon (Fig N in S1 Appendix) (instance 255 of 300, seed 0), which has the same number of games per subject as the human data. Human and model trial counts are then 27,205 vs 28,984 in the short horizon, 52,192 vs 55,444 in the long, and 79,397 vs 84,428 combined. Termination is constant within a single horizon and so is dropped in (a) and (b), and retained in (c).

### Parameter–symptom associations across horizon conditions

Every fitted parameter of each horizon’s winning model was correlated with every measure in the set defined in the Results, both across the whole sample and within the 20 subjects with the highest composite compulsion score, giving 190 tests for the short horizon, 266 for the long and 318 for the combined fit. Two, four and seven of these reached *p <* 0.05 uncorrected, against the 9.5, 13.3 and 15.9 expected under the null, so every scan returned fewer nominally significant associations than chance alone would produce, and none survived Benjamini–Hochberg correction at *q* = 0.05. Individual associations are therefore not tabulated and are not interpreted. Two of the nominally significant combined-horizons correlations are shown in Fig O to illustrate the scale of the effects the scan returns.

**Fig N.**
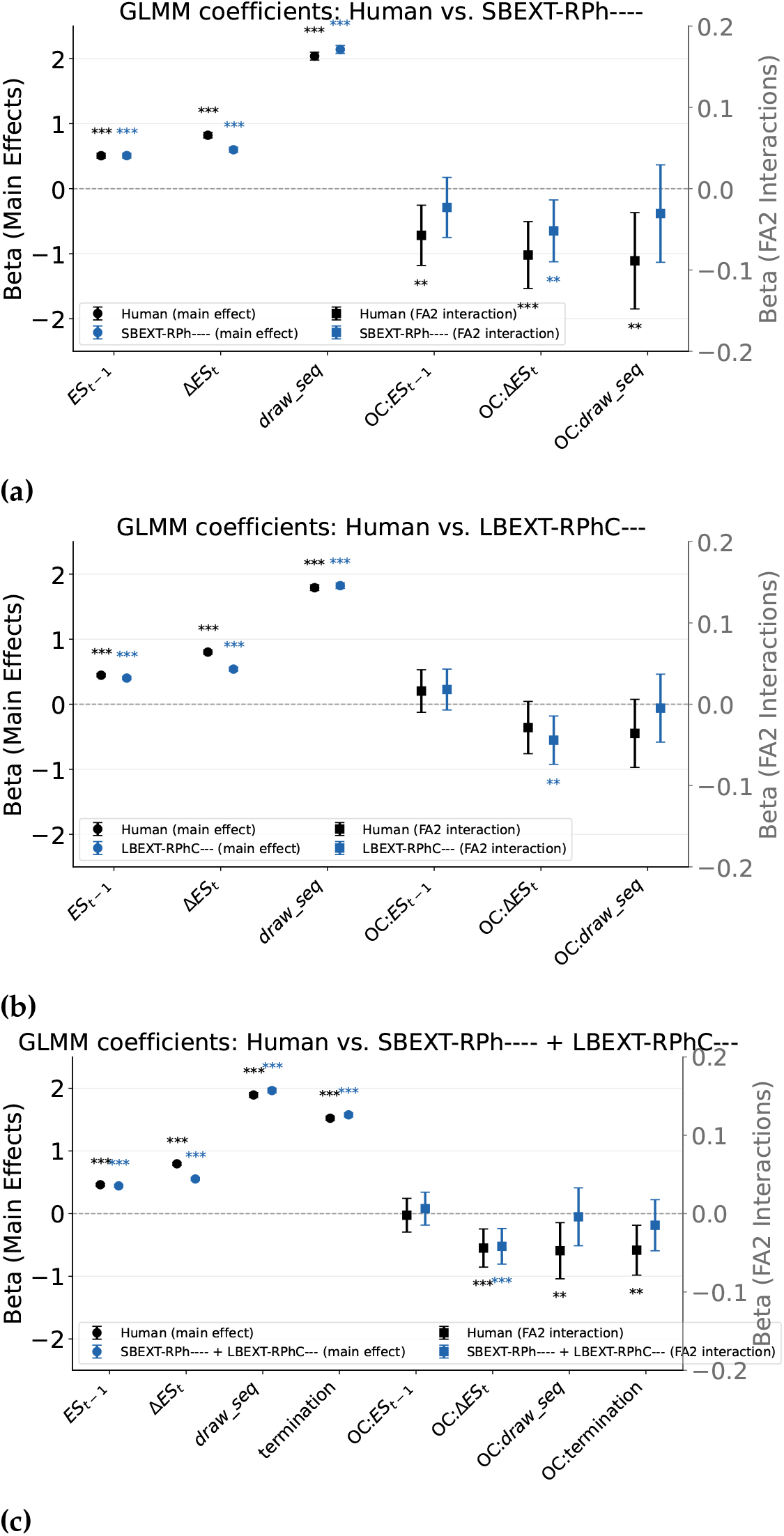
GLMM coefficients on a single simulated dataset matched in size to the human data. Conventions as in Fig I: fixed effects (*±*95% CI), main effects on the left axis (circles), FA2 interactions on the right axis (squares), asterisks giving each estimate’s own significance. The model is a single randomly drawn simulation rather than an ensemble average, so its intervals are directly comparable to the human ones. **(a)** Short horizon, `SBEXT-RPh----`. **(b)** Long horizon, `LBEXT-RPhC---`. **(c)** Both horizons combined.

**Fig O.**
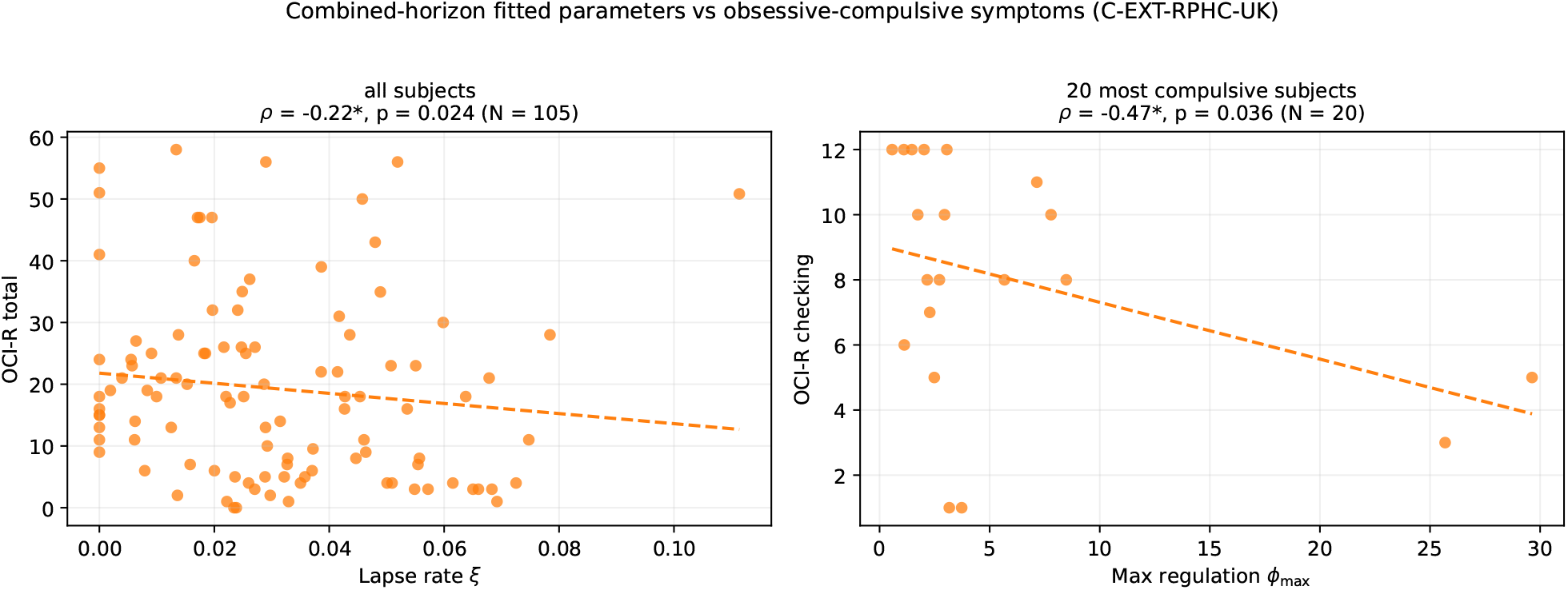
Combined-horizons fitted parameters vs. obsessive-compulsive symptoms. Each point is one subject; the dashed line is a least-squares fit shown for visual trend only, and Spearman *ρ* is annotated per panel (^∗^*p <* 0.05). *Left:* the lapse rate *ξ* vs. OCI-R total across all subjects. *Right:* maximum regulation *ϕ*_max_ vs. OCI-R checking within the 20 most compulsive subjects. Neither survives correction for the 318 tests in this scan, and neither is interpreted.

### Personalised per-subject model selection

Whether the single model structure selected for each horizon condition is representative of individual subjects, and what is gained by letting every subject take their own.

Two analyses elsewhere in the appendices bear directly on this:S1 Appendix, “Protected exceedance probabilities over the individualised models” asks the same question in random-effects terms, estimating how frequently each structure is used in the population rather than which one has the lowest total: it confirms the short-horizon result, finds the long horizon unable to distinguish the mechanisms at all (BOR = 0.872), and finds that the combined-horizon structure with the lowest cumulative BIC is not the one most subjects use. That last point is the sharpest statement of the heterogeneity this section documents. The cost accounting below then asks whether that heterogeneity is large enough to pay for identifying it subject by subject, and finds that it is not. The two are consistent: subjects do differ in which structure suits them, and the differences are still too small to justify fitting each subject their own.

Every comparison in the main text selects a single model structure per horizon condition whose concatenated BIC score values across the subjects was the lowest. To test whether that population-level winner is representative of individual subjects, we additionally computed one BIC per subject under every candidate model within a horizon condition (using that subject’s own observation count, BIC_*i*_ = *k* log *n*_obs,*i*_ −2 log ℒ_*i*_) and asked, independently for each subject, which structure minimises their own BIC. Summing each subject’s own best-fitting BIC gives the cumulative BIC of a “personalised” ensemble of 105 individually-selected models; we compare this against the cumulative BIC of the single population-level winning model *structure* applied to everyone – every subject is still fit with their own parameter values within that one shared structure, exactly as in the main-text comparisons; only the choice of *which* structure is used differs between the two cumulative BIC numbers, computed with the identical per-subject-summed formula used throughout this manuscript (Tables 7–10), so the two numbers differ only in whether model structure is personalised. Personalising model structure improved cumulative BIC in every horizon condition (Table G, Fig P), but by very different margins: the improvement was modest for the short and long horizons (ΔBIC = 470.4 and 469.8; 42/105 and 65/105 subjects improved by more than 2 BIC points, respectively) and larger still for the combined horizon (ΔBIC = 707.3; 88/105 subjects improved; Fig Q). This personalisation sweep spans the full candidate set: `C-EXT-RPHC-UK` is itself both the population-level (aggregate-BIC) winner *and* the single most common individual winner (40/105 subjects).

**Table G.** Personalised per-subject model selection vs. a single fixed model, by horizon condition. All 105 subjects, full candidate set (including forgetting-model variants). “Fixed model” is the population-level winner from the comparisons above; cumulative BIC values use the per-subject-summed convention described in the text. “N subjects improved” counts subjects whose personalised BIC is more than 2 points lower than under the fixed model (positive evidence for a structural difference, Kass & Raftery scale).

| Horizon | N subjects | Fixed model | Personalized cumulative BIC | Fixed cumulative BIC | Delta (fixed - personalized) | N subjects improved ($\geq 2$ BIC) |
| --- | --- | --- | --- | --- | --- | --- |
| Short | 105 | SB-XT-RPh---- | 28131.13 | 28601.57 | 470.43 | 42 |
| Long | 105 | LBE-T-RPhCL-- | 37318.45 | 37788.21 | 469.76 | 65 |
| Combined | 105 | C-EXT-RPHC-UK | 66415.46 | 67122.78 | 707.32 | 88 |

**Fig P.**
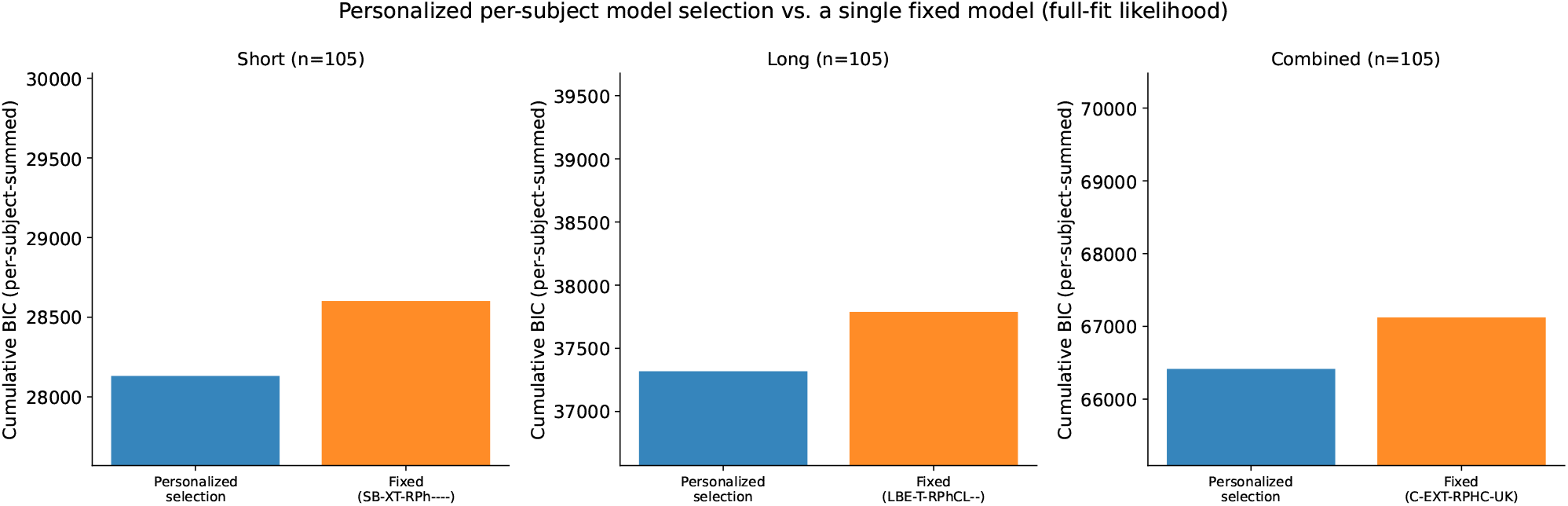
Cumulative BIC: personalised per-subject model selection vs. a single fixed model. Each panel is one horizon condition; bars give the cumulative (per-subject-summed) BIC for the personalised ensemble of individually-selected models vs. the single population-level winner applied to all subjects.

#### Protected exceedance probabilities over the individualised models

To check whether the models and mechanisms that win on the summed criteria are the ones most subjects actually use, we calculated the protected exceedance probability over the individualised fits. Every comparison above is a fixed-effects one, in which the per-subject criteria are summed and a model wins by having the lowest total; a large gain in a minority of subjects can therefore outweigh a small loss in the majority.

The exceedance probability is the posterior probability that a given candidate is more frequent in the population than any other, obtained by treating the model that generated each subject as a random variable and using 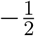 BIC as the per-subject log evidence [42]. On its own it can favour a candidate even when the data cannot separate the alternatives, since it is a statement about ordering that stays defined however weak the evidence. The protected exceedance probability removes that risk by mixing it with the uniform distribution in proportion to the Bayesian omnibus risk, the posterior probability that the frequencies are in fact all equal, so that every value is drawn towards 1*/K* when no candidate is genuinely distinguishable [43]. We ran it at two levels: at the model level every candidate for that horizon condition enters separately (25, 31 and 23 models), and at the family level the candidates are grouped by which recency mechanism they carry, with evidence summed within a family.

**Fig Q.**
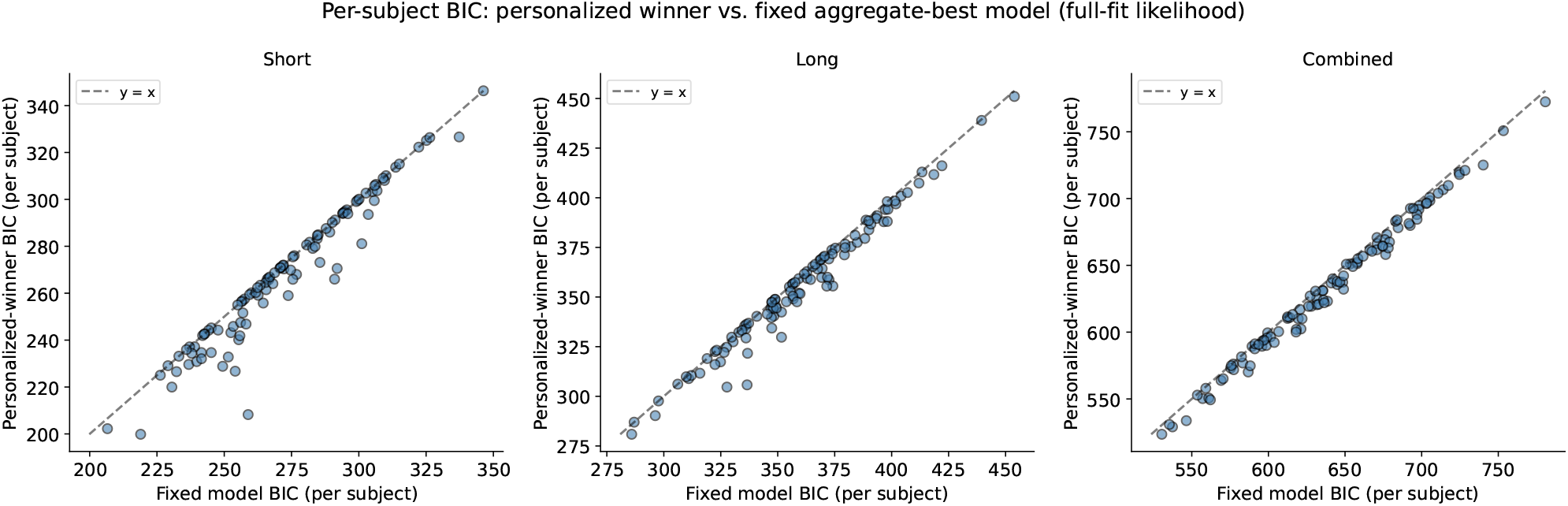
Per-subject BIC: personalised winner vs. fixed aggregate-best model. Each point is one subject; points below the identity line are better described by their own individually-selected model than by the population-level winner.

The three horizon conditions give three different answers (Table H, Fig R; the model-level results are in Table I). The short horizon is decisive and agrees with the fixed-effects result: `SB-XT-RPh----` has an estimated frequency of 0.49 and 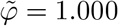, and at the family level the models carrying no recency mechanism take 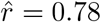 and 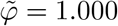 with BOR = 0.000. That the short horizon needs no recency mechanism is therefore not an artifact of summing criteria across subjects.

The long horizon is the opposite. At the model level the two leading candidates are a forgetting-based and an exaggeration-based model separated by almost nothing (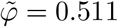 and 0.487), and at the family level the omnibus risk is BOR = 0.872, so all three protected exceedance probabilities collapse towards 1*/*3 (0.406, 0.303, 0.291). Read strictly, the long-horizon data do not identify which recency mechanism the population uses. This is the same conclusion the criterion-based comparison reaches when the two implementations are separated by under 100 BIC without agreeing on an ordering, stated in a form that makes the ambiguity explicit rather than leaving it to be inferred from a small margin.

The combined horizon is the one place where the random-effects and fixed-effects analyses disagree, and the disagreement is informative rather than contradictory. The random-effects winner is `C--XT-RPH--UK` 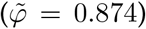, not `C-EXT-RPHC-UK`, which receives 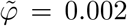. The reason is visible in the per-subject fits: `C--XT-RPH--UK` is the best model for 29 of 105 subjects while having a cumulative BIC of 73020.0, whereas `C-EXT-RPHC-UK` is best for only 9 subjects yet has the lowest cumulative BIC at 67122.8. The aggregate winner is therefore carried by large improvements in a minority of participants rather than by being the most common account, which is exactly the distinction a random-effects analysis exists to expose. At the family level the combined horizon divides between no recency mechanism 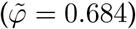 and exaggeration 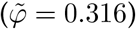, with forgetting excluded 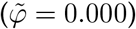.

Two limits are worth stating. The Laplace approximation 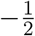 BIC stands in for the log evidence, so these results inherit its accuracy. And with 105 subjects spread over 23 to 31 candidates the model-level analysis is thinly supported, which is why the family-level result is the one we would rest an argument on.

**Table H.**
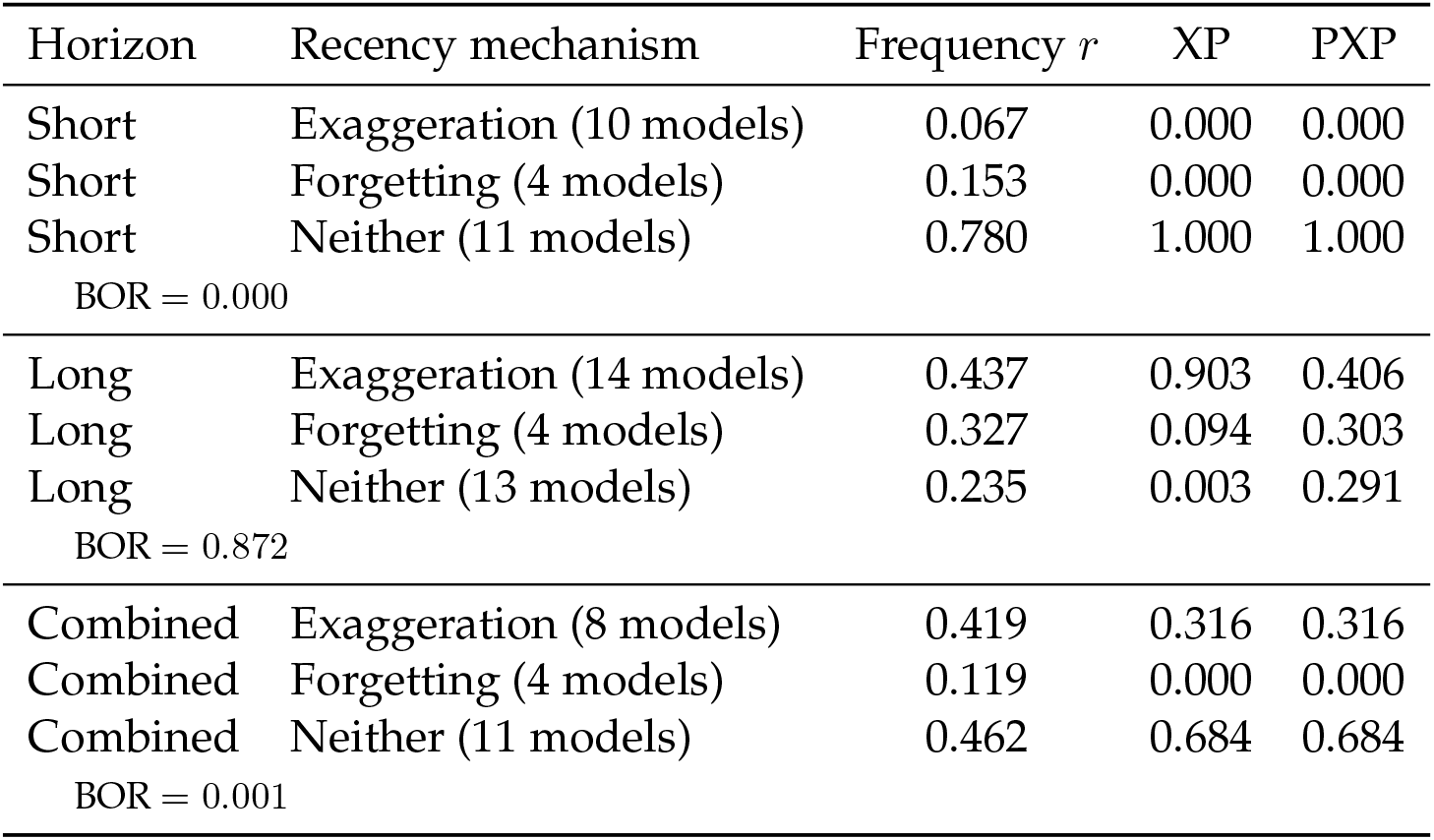
Protected exceedance probabilities by recency mechanism. Candidates are grouped by which recency mechanism they carry, with the number of models per family in parentheses, and the evidence summed within a family so that the comparison is between mechanisms rather than between individual models. 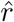 is the estimated population frequency, XP the exceedance probability, and PXP the protected exceedance probability. BOR is the Bayesian omnibus risk for that horizon condition; the long horizon’s 0.872 is what pulls its three PXPs towards 1*/*3.

**Fig R.**
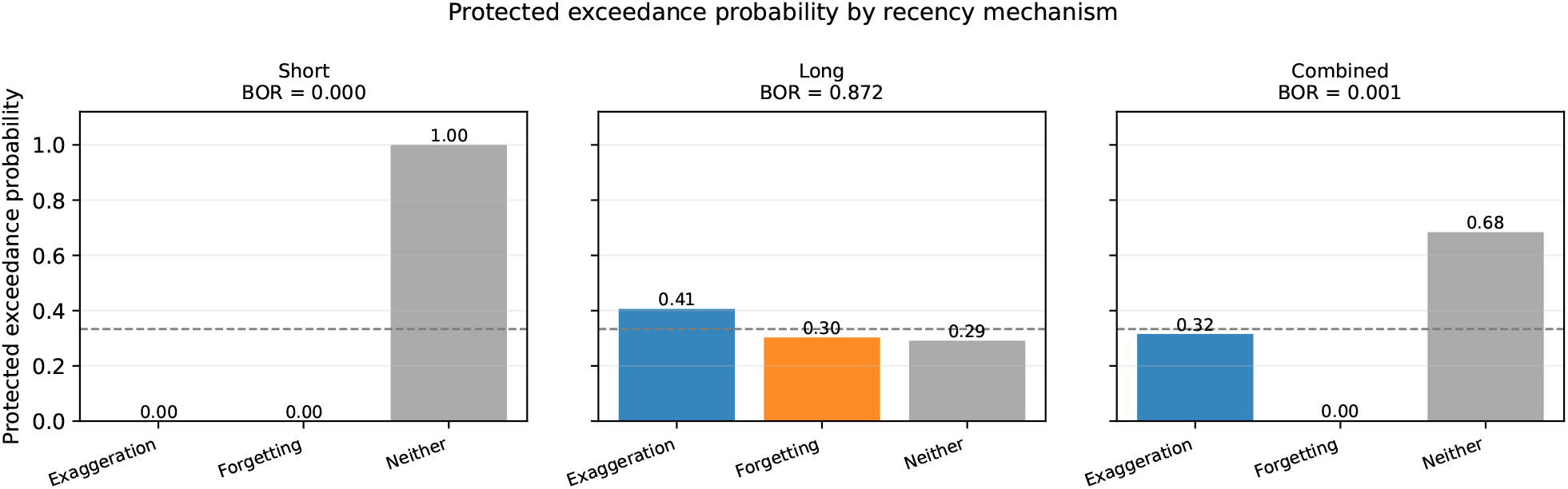
Protected exceedance probability by recency mechanism, per horizon condition. Dashed line, 1*/*3, the value every family takes when the data cannot distinguish them. The Bayesian omnibus risk is given above each panel: it is negligible in the short and combined horizons and 0.872 in the long horizon, where the three families are correspondingly indistinguishable.

**Table I.**
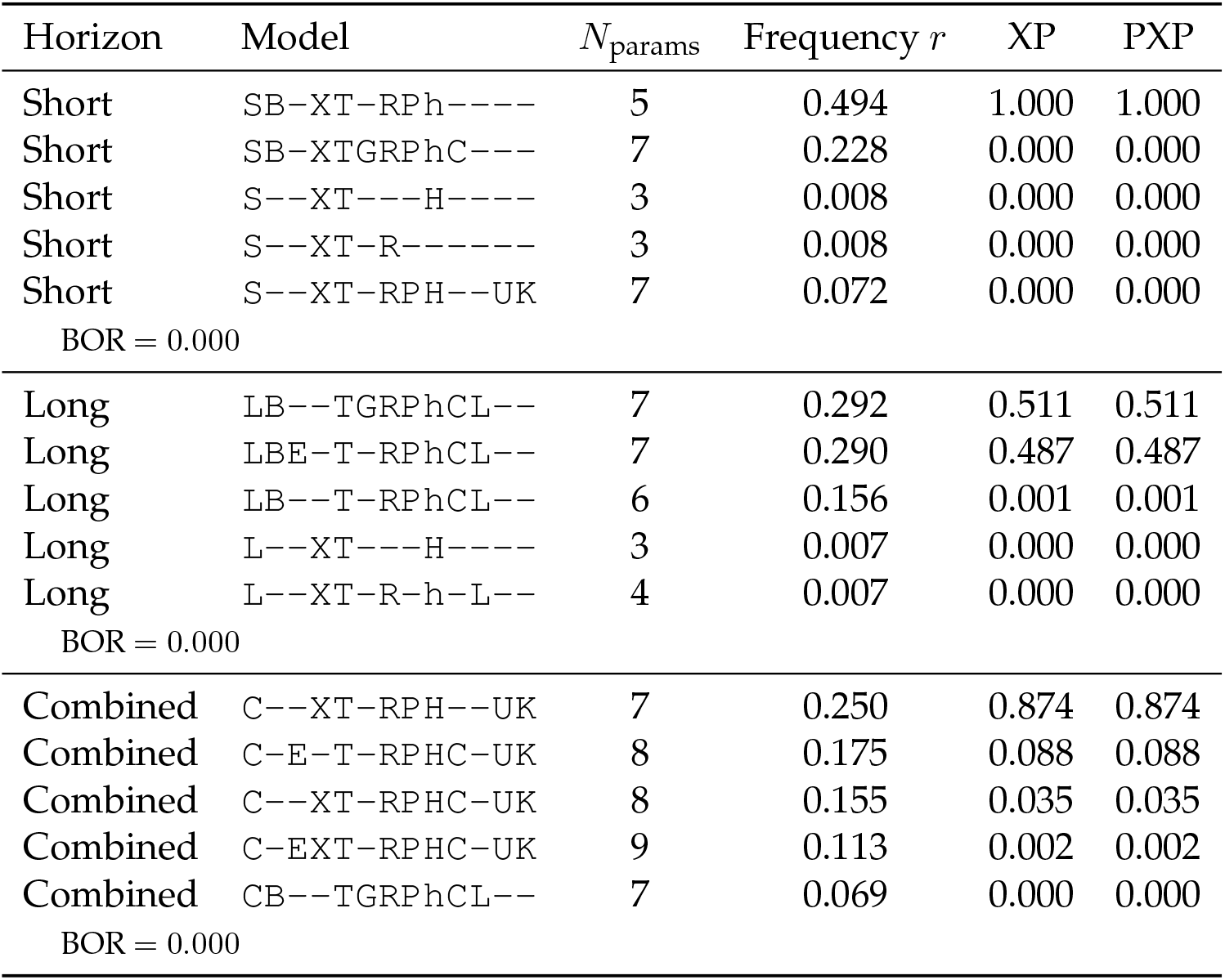
Protected exceedance probabilities, model level. The five leading candidates in each horizon condition, out of the 25, 31 and 23 models fitted for each condition respectively, ordered by PXP. Each row is a single model rather than a family, which is the comparison Table H deliberately does not make. Conventions as in Table H. The combined-horizon leader is not the model with the lowest cumulative BIC; see the text.

#### Does personalising pay for its own selection cost?

The comparison above credits personalisation with the whole BIC improvement while charging nothing for the choice that produced it. That choice is not free. Selecting, for each subject independently, whichever of the *M* candidate structures fits that subject best is itself an extra parameter, so each subject carries one extra parameter beyond those of the structure they were assigned. Personalisation is worth adopting only if the BIC it saves exceeds what the choice costs.

Under the *parameter* charge, each subject’s extra parameter costs log *n*_obs,*i*_ in the per-subject-summed convention used throughout, so ∑_*i*_ log *n*_obs,*i*_ in total. Under the *selection* charge, the choice is charged its information cost under a uniform prior over the *M* candidates available for that horizon, 2 log *M* per subject. Note the contrast with the fixed regulation values of S1 Appendix, “Parameter and model recovery and sensitivity”, which are single numbers shared by the whole sample and so cost log *N* once: here every subject makes their own choice, so the charge is per subject and roughly two orders of magnitude larger.

Table J (full likelihood), Table K (commit likelihood) and Fig S report both. The two accounts give similar magnitudes, between 603.6 and 721.1 BIC, and they agree on the short and long horizons: in neither does personalisation recover its cost, under either likelihood. For the full-fit short horizon the choice costs 603.6–676.0 against a gain of 470.4, and for the long horizon 659.5–721.1 against 469.8, leaving deficits of 133.2–205.5 and 189.8–251.4 respectively. The commit-likelihood versions are worse still, the short horizon most of all, where a gain of only 47.7 is set against a cost above 600.

The combined horizon is the one case that lands on the boundary, and the two accounts disagree about which side of it. Under the parameter charge personalisation falls just short (gain 707.3 against cost 708.3, a deficit of 1.0 for the full fit; 660.4 against 708.3 for the commit fit), while under the selection charge it just clears it (a surplus of 58.2 and 11.3 respectively, the combined horizon having the smallest candidate set at *M* = 22). A result that changes sign between two equally defensible ways of charging for the same choice is not evidence for personalisation; the honest reading is that the combined horizon breaks even and the short and long horizons do not.

This tempers the interpretation of the improvements reported above rather than overturning the observation behind them. That 96 of 105 subjects are individually better fit by some structure other than the population-level winner remains true, and remains evidence that subjects differ in which mechanisms they require. What the accounting shows is that this heterogeneity is not by itself strong enough to justify reporting a personalised model as the account of the data: the per-subject improvements, summed, are about the same size as the cost of having chosen per subject at all.

**Table J.** Does personalising model structure pay for its own selection cost? Full likelihood. “ΔBIC gained” is the improvement from personalising reported in Table G. The parameter charge costs each subject one extra parameter for their own choice, _*i*_ log *n*_obs,*i*_; the selection charge costs the choice 2 log *M* per subject, with *M* the number of candidate structures for that horizon (given in parentheses). “Net” is the gain minus the cost, positive when personalisation pays. “Borderline” marks a horizon where the two accounts disagree in sign.

| Horizon | Fixed model | ΔBIC gained | Parameter charge | | Selection charge | | Worth it? |
| --- | --- | --- | --- | --- | --- | --- | --- |
| | | | Cost | Net | Cost ($M$) | Net | |
| Short | SB-XT-RPh---- | 470.4 | 603.6 | -133.2 | 676.0 (25) | -205.5 | No |
| Long | LBE-T-RPhCL-- | 469.8 | 659.5 | -189.8 | 721.1 (31) | -251.4 | No |
| Combined | C-EXT-RPhC-UK | 707.3 | 708.3 | -1.0 | 649.1 (22) | +58.2 | Borderline |

**Table K.** Does personalising model structure pay for its own selection cost? Commit likelihood. Conventions as in Table J, applied to the personalised selection carried out under the commit likelihood.

| Horizon | Fixed model | ΔBIC gained | Parameter charge | | Selection charge | | Worth it? |
| --- | --- | --- | --- | --- | --- | --- | --- |
| | | | Cost | Net | Cost ($M$) | Net | |
| Short | SBEXT-RPh---- | 47.7 | 603.6 | -556.0 | 649.1 (22) | -601.4 | No |
| Long | LBEXT-RPhC--- | 432.1 | 659.5 | -227.4 | 699.8 (28) | -267.7 | No |
| Combined | C-EXT-RPhC-UK | 660.4 | 708.3 | -47.9 | 649.1 (22) | +11.3 | Borderline |

**Fig S.**
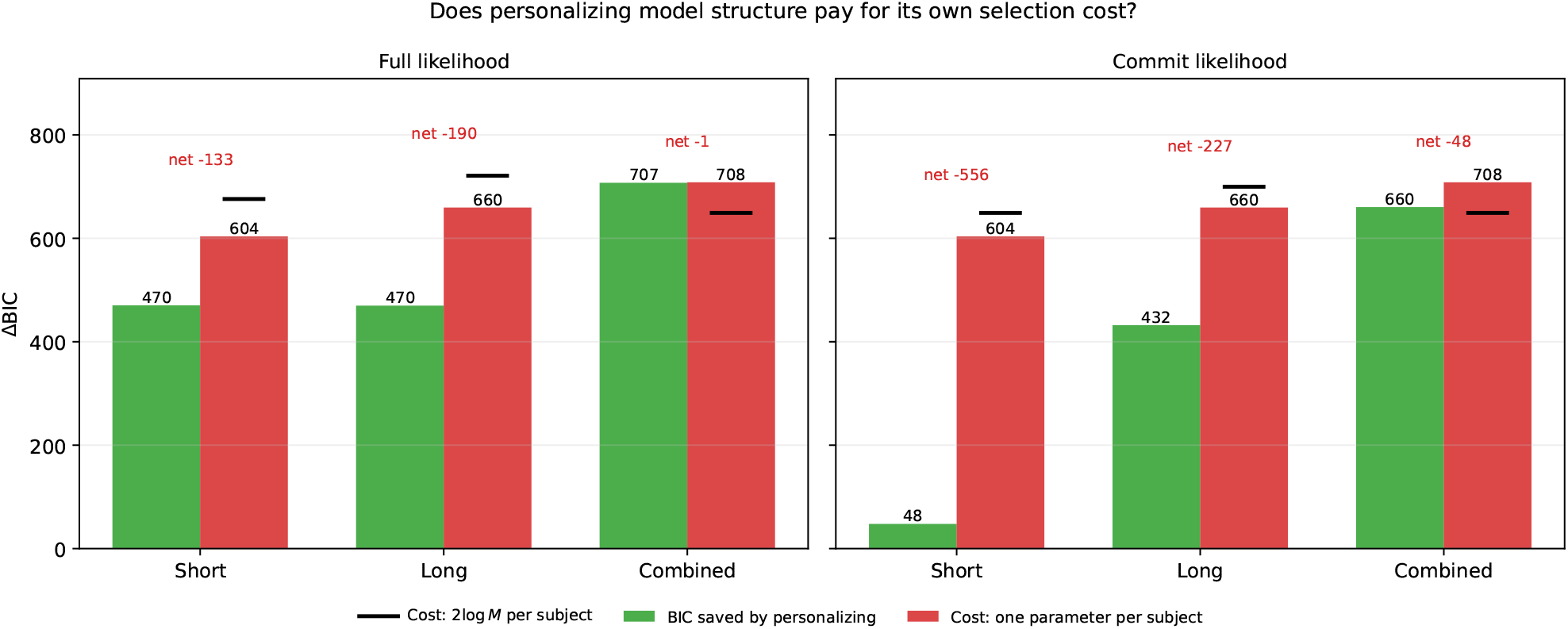
BIC saved by personalising, against the cost of the choice. Green bars give the cumulative BIC improvement from letting each subject take their own structure; red bars give the cost of that choice charged as one extra parameter per subject, and black dashes the same cost charged as 2 log *M* per subject. Personalisation pays only where the green bar exceeds both. *Left:* full likelihood. *Right:* commit likelihood. The number above each pair (‘net’) is effective cost of the parameter.

## Notes

### Competing Interest Statement

The authors have declared no competing interest.

